# Long-term biodiversity responses to anthropogenic change are spatially and taxonomically heterogeneous across coastal wetlands

**DOI:** 10.64898/2026.11.03.763952

**Authors:** Elena Baños, Javier de la Casa, Andrew B. Cundy, Bastian Hambach, Enrique González-Ortegón, Erik J. de Boer, Jesús Zarcero Pimentel, Xavier Benito, Xavier Turon, Luke E. Holman, Marc Rius, Sandra Nogué

## Abstract

Coastal wetlands have undergone centuries of anthropogenic modification, yet their long-term biodiversity trajectories remain poorly understood because ecological monitoring rarely extends beyond recent decades. Sedimentary ancient DNA (*seda*DNA) offers a historical perspective for evaluating how biodiversity patterns vary across environmental contexts and timescales. We combined COI and 18S *seda*DNA metabarcoding with ²¹⁰Pb and radiocarbon chronologies, sediment geochemistry and stable-isotope records from eight coastal wetlands across four regions of the Iberian Peninsula. We reconstructed changes in eukaryotic amplicon sequence variant (ASV) richness and taxonomic composition and examined their relationships with an Anthropogenic Enrichment Index (AEI) based on ten elements potentially associated with human activities. Richness-AEI relationships differed among sediment records and genetic markers. Higher enrichment was associated with lower richness for both markers in one Cádiz record, whereas COI richness increased with enrichment in the Ebro Delta. However, after accounting for calendar time, only the positive association in the Ebro Delta remained statistically significant, indicating that several initial relationships were partly explained by shared temporal trends. Exploratory rolling correlations further suggested that the direction of richness-enrichment associations varied across historical periods within individual records. Taxonomic profiles differed between markers and wetlands, while community composition was strongly differentiated among sediment records, highlighting ecological variation not captured by richness alone. Multiproxy changes also followed locally specific configurations rather than a common regional trajectory. Together, these findings show that long-term relationships between sedimentary enrichment and biodiversity are context-dependent and sensitive to the temporal perspective considered. Integrating molecular, geochemical and isotopic records can therefore improve the historical interpretation of biodiversity change and inform conservation approaches that account for taxonomic composition, environmental history and locally appropriate reference conditions.

## Introduction

Coastal wetlands, including estuaries, saltmarshes and coastal lagoons, support biodiversity and provide important ecosystem functions, including nutrient cycling, carbon sequestration, shoreline protection, and nursery habitats for numerous species (Barbier et al., 2011; Beck et al., 2001; Mcleod et al., 2011; Mitsch et al., 2015). At the same time, they are among the environments most heavily affected by human activities. Industrialisation, urban development, agricultural expansion, and maritime activity have transformed coastal systems over recent centuries, modifying sediment dynamics, nutrient inputs, and ecological communities (Halpern et al., 2008; Lotze et al., 2006; Nogué et al., 2025; Syvitski et al., 2005). Many estuarine environments have consequently undergone substantial biodiversity change, often accompanied by habitat degradation and biodiversity loss (Barbier et al., 2011; Jackson et al., 2001; Lotze et al., 2006).

Assessing the long-term consequences of these transformations remains difficult because most ecological observations cover only recent decades (Willis et al., 2010). Monitoring programmes therefore provide limited information on ecosystem conditions before major anthropogenic modification (Kopf et al., 2015; McClenachan et al., 2012). This limited historical context complicates the identification of ecological baselines and restricts understanding of how biological communities respond to environmental pressures over longer timescales (Nogué et al., 2017; Willis & Birks, 2006). The problem is particularly pronounced in coastal ecosystems, where climatic variability, sea-level change, sediment dynamics, land-use change and anthropogenic inputs interact across multiple temporal and spatial scales (Dalrymple et al., 1992; Nogué et al., 2021).

Sedimentary archives extend ecological observations beyond the period covered by direct monitoring. Biological remains and geochemical signatures preserved in sediments retain evidence of past environmental conditions and ecosystem dynamics (Battarbee, 2000; R. S. Bradley, 1999). Sedimentary ancient DNA (*seda*DNA) has emerged as a promising approach for reconstructing past biological communities in aquatic and terrestrial environments (Capo et al., 2021; Holman et al., 2023; Willerslev et al., 2003). By recovering DNA fragments preserved within sediments, metabarcoding can characterise historical community composition across a broad range of taxonomic groups (Heintzman et al., 2023; Pedersen et al., 2013). Applications in coastal wetlands nevertheless remain comparatively scarce, particularly at regional scales, where multiple records with contrasting ecological and environmental histories can be compared (Campbell et al., 2025; Nguyen et al., 2023).

Coastal-wetland biodiversity comprises several major eukaryotic lineages, including fungi, metazoans, plants, and phylogenetically diverse protists. Because these groups differ in their ecological roles and responses to environmental change, analyses focused on a single lineage provide a partial view of long-term community dynamics. Broad-range metabarcoding can recover several components of eukaryotic diversity simultaneously, although taxon detection depends partly on primer affinity, marker resolution, and reference-database coverage. As a consequence, COI and 18S provide complementary rather than interchangeable perspectives on past eukaryotic communities (Casey et al., 2021; Deiner et al., 2017).

Anthropogenic pressures can substantially alter biodiversity without necessarily producing consistent declines in local richness. Large-scale biodiversity syntheses indicate that local richness trends are frequently centred around zero, even when community composition and species turnover change markedly through time (Dornelas et al., 2014; Hillebrand et al., 2018). Hydrodynamic conditions, habitat structure, nutrient availability, and local environmental history may alter both the magnitude and direction of biodiversity responses. As a result, richness and community composition may follow nonlinear and site-specific trajectories rather than a common response to increasing anthropogenic pressure (Ficetola et al., 2018; Halpern et al., 2008; Holman et al., 2025). Here, we characterised historical variation in sedimentary elemental enrichment using a multielement geochemical index designed to capture changes in elements potentially associated with anthropogenic inputs.

We combined COI and 18S *seda*DNA metabarcoding with ^210^Pb and radiocarbon dating, X-ray fluorescence (XRF) geochemistry, and stable-isotope analyses (δ^13^C and δ^15^N)across eight sediment records from Atlantic and Mediterranean coastal wetlands of the Iberian Peninsula. We reconstructed long-term variation in overall and taxon-specific ASV richness, characterised differences in community composition among sediment records, and examined associations between richness and sedimentary multielement enrichment. By comparing records with contrasting environmental histories, we assessed whether richness–enrichment relationships were consistent across wetlands and metabarcoding markers, whether these associations persisted after accounting for calendar time, and whether their direction varied across historical periods. We also evaluated how taxonomic and compositional differences, together with concurrent changes in biological, geochemical and isotopic proxies, contributed to the interpretation of biodiversity trajectories. This approach provides a long-term perspective on the extent to which biodiversity change can be characterised by richness trends and sedimentary enrichment alone, or instead requires consideration of taxonomic composition, temporal context and local environmental history.

## Methods

### Study sites and sampling strategy

Eight sediment cores were collected between 2022 and 2024 from eight coastal wetland sites distributed among four estuarine systems of the Iberian Peninsula, encompassing both Atlantic and Mediterranean regions (Fig. S1; Table S1). Northern sites included three saltmarsh locations within the Santoña estuarine system (Cantabria): Cicero (CCO), Colindres (COL) and Santoña (STO) (Table S1). Southern Atlantic sites included two cores from Saltés Island (Huelva, Andalucía), representing contrasting hydrodynamic settings: HIT, located along the Tinto River margin, and HIO, facing the open marine sector. Two additional sites were located within the Bay of Cádiz (Andalucía): CMB, situated in saltmarshes within a restricted military area, and CSD, located in the Salinas Dolores complex. Finally, the Mediterranean site DEE was sampled from the Encanyissada lagoon in the Ebro Delta (Delta de l’Ebre, Catalunya) (Fig. S1). At each site, a sediment core was retrieved using a 50-cm Russian-type peat corer. Immediately after collection, sediments were transferred into semi-cylindrical PVC tubes (5 cm internal diameter), sealed to minimise contamination, and transported at 4 °C to the CEAB within 72 h.

Core subsampling was conducted in a dedicated clean DNA laboratory (PCR-free, positive-pressure environment) at CEAB under strict contamination-control procedures. After removing the outer sediment layer, 2–5 g of sediment were collected at 1 cm intervals using sterile tools and transferred to sterile containers. All subsamples were stored at −20 °C until DNA extraction.

### Chronology

Sediment chronologies were constrained using ^210^Pb dating conducted at GAU-Radioanalytical, in the School of Ocean and Earth Science at the University of Southampton (UK), and accelerator mass spectrometry (AMS) radiocarbon (^14^C) dating at the Poznań Radiocarbon Laboratory (Poland). Recent sediment accumulation rates were estimated from a total of 92 ^210^Pb measurements distributed among the eight sediment cores. Five samples from the lower sections of DEE, CSD, CMB, HIT, and HIO were additionally dated by AMS ^14^C to provide independent chronological information beyond the unsupported-^210^Pb dating horizons.

Core-specific sediment accumulation rates were estimated using the Constant Flux–Constant Sedimentation (CF:CS) model (Krishnaswamy et al., 1971), based on linear regressions of ln-transformed unsupported ^210^Pb activity against sediment depth. The lower limit of the ^210^Pb chronology was defined as the depth at which unsupported ^210^Pb declined to background activity. Calendar ages used in the temporal analyses were derived from the corresponding core-specific CF:CS accumulation rates. Ages assigned below the unsupported-^210^Pb dating horizon therefore represent linear extrapolations that assume continuation of the recent accumulation rate and were interpreted with caution.

AMS ^14^C determinations were calibrated using the IntCal20 Northern Hemisphere atmospheric calibration curve (Reimer et al., 2020) in R using the *rcarbon* package (v1.5.2; Crema & Bevan, 2021). No marine-reservoir or empirical old-carbon correction was applied. Because the dated carbon fraction was not independently characterised, the calibrated ages were interpreted as estimates of the age of the dated carbon rather than direct estimates of the depositional age of the surrounding sediment. Incorporating these determinations as depositional constraints could therefore have introduced an unquantified offset into integrated age–depth models. The ^14^C ages were instead retained as independent chronological comparisons, while the primary working chronologies were derived consistently from the core-specific ^210^Pb models. This choice does not imply that the ^210^Pb chronologies were intrinsically more accurate: ages extrapolated below the unsupported-^210^Pb horizons also remain uncertain and were interpreted cautiously. Further analytical and calibration details are provided in Method S1.

### Stable-isotope and geochemical characterisation

Stable carbon and nitrogen isotope compositions (δ^13^C and δ^15^N) were measured in 132 freeze-dried and homogenised sediment samples using an elemental analyser coupled to an isotope-ratio mass spectrometer. The C:N ratio was calculated on a molar basis after converting elemental mass percentages to molar quantities. Isotope values are reported in standard δ notation relative to Vienna Pee Dee Belemnite for carbon (Hoffman & Rasmussen, 2022) and atmospheric N2 for nitrogen (Brand et al., 2014). Analytical precision was better than ± 0.2‰ for both isotopes. Analyses were conducted at the Stable Isotope Laboratory from the National Oceanography Centre Southampton at the University of Southampton, UK.

Elemental composition was measured at contiguous 1-cm intervals throughout each sediment core using a Niton XL3p+ handheld X-ray fluorescence analyser at Universitat Autònoma de Barcelona/CREAF. Each measurement was performed on approximately 0.5 g of freeze-dried and homogenised sediment for 180 s, and elemental values were reported by the instrument in parts per million (ppm). To identify isolated analytical artefacts, elemental profiles were screened in R using a moving-window Hampel-type filter combined with a nearest-neighbour criterion (Hampel, 1974; Pearson et al., 2016). Measurements flagged by either criterion were treated as missing. Full filtering and replacement procedures are described in Methods S2 and S3.

### Selection of elements of anthropogenic enrichment

Eleven candidate elements were defined a priori as potential indicators of metal contamination or nutrient enrichment: Pb, Zn, Cu, Sb, Cd, Hg, Sn, Cr, Ni, As and P. Their documented associations with industrial, urban, mining, maritime, agricultural and wastewater-related inputs, together with the rationale and supporting references for their inclusion, are summarised in Table S2a. P was included as a potential indicator of nutrient enrichment; its inclusion did not imply an expected negative relationship with biodiversity because its ecological effects may depend on the intensity of enrichment. Sb was excluded because measurements were unavailable for some sediment cores, preventing its consistent inclusion in cross-record comparisons. The remaining ten elements were used to calculate the enrichment indices, and site-specific means and standard deviations of their cleaned XRF measurements are reported in Table S2b.

XRF-derived elemental values were treated as compositional data and centred log-ratio (CLR) transformed before enrichment was quantified (Aitchison, 1982; Bertrand et al., 2024; Weltje & Tjallingii, 2008). Because the CLR transformation requires complete and strictly positive compositions, missing and non-positive measurements were replaced separately for each element by half the minimum positive value observed for that element before transformation. This replacement was applied as an operational preprocessing step and was not interpreted as an estimate of the true elemental concentrations (Martín-Fernández et al., 2003). Further details and the CLR equation are provided in Method S3.

### Definition of baseline conditions and element-specific enrichment

A separate reference distribution was defined for each element within each sediment record. In CMB, CSD, DEE and STO, all ten elements had at least three observations assigned to dates before 1600 CE, and these observations were used as an early historical reference. The year 1600 Common Era (CE) was selected as an operational cutoff intended to provide reference conditions preceding large-scale industrialisation across the study region; it was not interpreted as a boundary between pristine and anthropogenically affected conditions. Earlier sediments may already record human influences associated with activities such as mining, metallurgy, agriculture, urban development or maritime activity (Mil-Homens et al., 2016), and the pre-1600 observations were therefore treated as an operational historical reference rather than as an unimpacted natural background. CCO, COL, HIT and HIO contained no observations dated before 1600 CE, and the complete corresponding elemental series were therefore used as internal statistical references. Consequently, enrichment scores were expressed relative to site- and element-specific reference conditions and did not represent enrichment above a common pre-industrial baseline across all sediment records. As a diagnostic assessment of the temporal cutoff, robust elemental variability was compared among the consecutive 200-year windows 1400–1599 CE, 1600–1799 CE and 1800–1999 CE (Method S3; Fig. S2).

Enrichment was quantified from the CLR-transformed values using a robust z-score:

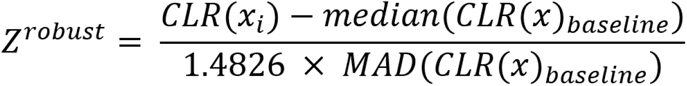

where CLR(xᵢ) is the CLR-transformed value of a given element, and the median and MAD were calculated from the corresponding site- and element-specific baseline. The MAD was multiplied by a scaling factor of 1.4826 to make it comparable to the standard deviation under a normal distribution (Rousseeuw & Croux, 1993; Rousseeuw & Hubert, 2018). This robust standardisation reduces the influence of extreme values when estimating the central tendency and variability of geochemical reference conditions, with positive scores indicating enrichment above the reference median relative to its robust variability (Reimann et al., 2005).

### Complementary selection of site-specific enrichment elements

As a complementary analysis, enrichment magnitude and recurrence were evaluated separately for each candidate element and sediment record (Method S3; Fig. S5). An element was retained in the site-specific set when strong enrichment, operationally defined as z^robust^ > 3, occurred in at least two observations from the corresponding record. This criterion was used to identify locally recurrent enrichment signals and was not interpreted as a universal threshold for contamination. Magnitude and recurrence were kept separate so that large but isolated anomalies could be distinguished from repeated enrichment. All candidate elements, including those not retained, are shown in Fig. S5.

### Calculation of general and site-specific Anthropogenic Enrichment Index

Multielement enrichment was summarised using a composite Anthropogenic Enrichment Index (AEI) based on the element-specific robust z-scores. Negative scores were first set to zero:

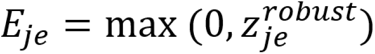

where Eje is the positive enrichment score of element e in sample j. Consequently, values at or below the site- and element-specific reference median did not offset the contributions of positively enriched elements.

The general AEI was calculated as the equal-weight arithmetic mean of the positive enrichment scores for the ten candidate elements:

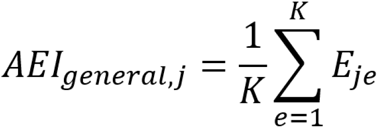

The general AEI was calculated only when valid enrichment scores were available for all ten elements, thereby maintaining a constant element composition, weighting scheme and denominator across sediment samples and records.

The site-specific AEI was calculated using the same aggregation rule but was restricted to the elements retained for the corresponding site:

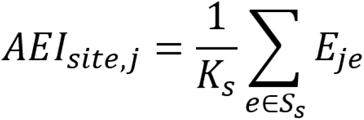

where Ss is the set of elements retained at site s, and Ks is the number of elements in that set. Element-specific robust z-scores placed the elements on a common relative scale before aggregation, and arithmetic averaging assigned equal weight to every included element (Greco et al., 2019; OECD et al., 2008). Both formulations were interpreted as relative measures of multielement enrichment above the corresponding site-specific reference distributions, rather than as established measures of contaminant concentration, toxicity or ecological risk. Positive enrichment scores indicate deviations in the relative representation of selected elements within the sedimentary composition and do not, by themselves, establish an anthropogenic origin. The AEI was therefore used as a geochemical indicator of potentially human-associated environmental change, whose interpretation requires consideration of local sedimentary processes and environmental history.

### Sensitivity of the AEI to site-specific element selection

The robustness of the reconstructed enrichment histories to element selection was evaluated by comparing the general and site-specific AEIs graphically and statistically at the site and pooled levels (Fig. S6). Similarity in the temporal ranking of samples was assessed using Spearman’s rank correlation coefficient, whereas numerical agreement was evaluated using Lin’s concordance correlation coefficient (CCC; Lin, 1989) and mean paired differences. Concordance was recalculated after separately standardising both indices within each site to assess similarity in temporal trajectories after removing site-specific differences in location and scale (Method S3).

The general AEI was defined a priori as the primary formulation for subsequent cross-record analyses because it maintained a common element set, equal-weighting scheme and denominator. The site-specific AEI was retained as a complementary sensitivity analysis. As a complementary sensitivity analysis, the same site- and marker-specific analyses were repeated separately using the element-specific robust z-scores of Pb and Hg instead of the AEI (Method S3).

### DNA extraction

DNA was extracted from 122 sediment samples, using the DNeasy PowerMax Soil Kit (QIAGEN) according to the manufacturer’s instructions and eluted in 1.2 mL of the kit-provided elution buffer. All extractions were conducted in a dedicated clean DNA laboratory at CEAB under stringent contamination-control procedures.

Contamination was monitored using 12 subsampling blanks, 16 extraction blanks and eight no-template PCR controls in each of the 11 libraries, yielding 116 negative controls per marker. DNA concentrations were quantified using a Qubit High Sensitivity assay, and extracts were stored at −20 °C prior to amplification.

### Library preparation and sequencing

Two metabarcoding markers were targeted: a fragment of the mitochondrial cytochrome c oxidase I (COI) gene (∼313 bp) amplified with the Leray-XT primers (Wangensteen et al., 2018), and the V7 region of the nuclear 18S rRNA gene (∼110 bp) amplified with 18S_allshort primers (Guardiola et al., 2015) (Table S3). Amplicons were generated using a twin-tagging dual-indexing approach (Bohmann et al., 2022; Kircher et al., 2012), with identical 8-bp barcode sequences (minimum Hamming distance ≥ 3) appended to both forward and reverse primers to reduce tag-switching artefacts (Schnell et al., 2015).

For each marker, eight independent PCR replicates were performed per sample in 40 µL reactions containing 4 µL of *seda*DNA template, 20 µL AmpliTaq Gold 360 Master Mix (Thermo Fisher Scientific), 11.68 µL laboratory water, 0.32 µL BSA (20 mg/mL; Fisher Scientific), and 2 µL of each primer (5 µM). Thermal cycling conditions differed between markers. For COI, amplification consisted of an initial denaturation at 95 °C for 10 min, followed by 35 cycles of denaturation at 94 °C for 1 min, annealing at 45 °C for 1 min and extension at 72 °C for 1 min. A final extension was performed at 72 °C for 5 min, followed by a hold at 4 °C. For 18S, amplification consisted of an initial denaturation at 95 °C for 10 min, followed by 45 cycles of denaturation at 95 °C for 30 s, annealing at 45 °C for 30 s and extension at 72 °C for 30 s. A final extension was performed at 72 °C for 5 min. PCR products were purified using the MinElute PCR Purification Kit (QIAGEN, Hilden, Germany), quantified fluorometrically (Qubit), normalised to equimolar concentrations and pooled. Sequencing libraries (n = 11) were prepared using the TruSeq PCR-Free protocol (Illumina) and sequenced on an Illumina NovaSeq 6000 platform (COI: 2 × 250 bp; 18S: 2 × 150 bp). Negative controls were included in sequencing libraries without normalisation due to low DNA yield.

### Bioinformatic processing

Raw paired-end reads were demultiplexed using Cutadapt v2.3 (Martin, 2011) with sample-specific dual barcodes and combined tag–primer sequences (∼36 bp), allowing an error rate of 0.1 (--error-rate 0.1) and no indels (--no-indels). Reads were separated into sense and antisense orientations (--pair-adapters, --pair-filter = both) and processed independently. Given the use of 8-bp twin tags, amplicons recovered in both orientations were retained and analysed separately throughout the workflow.

Sequence inference was performed using DADA2 v1.12 (Callahan et al., 2016). Reads were quality-filtered (maxEE = 1, truncQ = 2), trimmed to marker-specific lengths (COI: 250/230 bp; 18S: 120/110 bp), denoised, and merged with a minimum overlap of 18 bp. Chimeric sequences were identified and removed using the *removeBimeraDenovo* algorithm. Amplicon sequence variants (ASVs) obtained from antisense reads were reverse-complemented and merged with those from sense reads to generate a unified sequence table.

ASV curation was conducted using LULU (Frøslev et al., 2017), implemented via a custom R function (ApplyLulu; doi:10.5281/zenodo.4671710), with a minimum similarity threshold of 98%. Sequence similarity and co-occurrence patterns were evaluated using *vsearch* (Rognes et al., 2016) in self-matching mode (--usearch_global, --self), applying a minimum identity threshold of 0.84 (--id 0.84, --iddef 1), minimum query coverage of 0.9 (--query_cov 0.9), and a maximum of ten hits per query (--maxhits 10).

Taxonomic assignment was performed using BLAST+ v2.14.1 (Camacho et al., 2009) against the NCBI *nt* database (March 2024), retaining up to 300 alignments per query (-num_alignments 300). BLAST outputs were parsed in R using a custom script (ParseTaxonomy; doi:10.5281/zenodo.4671710), applying identity and coverage thresholds and resolving multiple matches using a lowest common ancestor approach. Species-level assignments were considered reliable at ≥97% identity and ≥85% coverage for COI, and at 100% identity (≥98% coverage) for 18S. Ambiguous and environmental taxonomic assignments were rejected, while the corresponding ASVs were retained as unassigned sequences in analyses that did not require taxonomic classification.

Following taxonomic assignment, potential contamination was addressed using an ASV-specific maximum-subtraction procedure. For each ASV, the highest read count observed across all negative controls was subtracted from its count in every PCR replicate, and resulting negative values were set to zero. The datasets were subsequently filtered at the ASV and PCR-replicate levels. First, singleton and doubleton ASVs, defined as ASVs represented by only one or two reads across the entire dataset, were removed. Second, within each sediment horizon, ASVs detected in only one PCR replicate were excluded. Third, ASV counts below three reads within an individual PCR replicate were set to zero. Finally, PCR replicates with fewer than 1,000 remaining reads were removed from the dataset.

For downstream community analyses, counts were converted separately within each retained PCR replicate to relative abundances by dividing the abundance of each ASV by the total number of retained reads in that replicate. Replicates belonging to the same sediment horizon were then combined by calculating the mean relative abundance of each ASV across the retained PCR replicates. The sediment horizon, rather than the individual PCR replicate, was therefore the unit used in the temporal, taxonomic and community-composition analyses.

Eukaryotic ASV richness was calculated independently for COI and 18S as Hill diversity of order zero (q = 0), corresponding to the number of retained eukaryotic ASVs detected in each sediment horizon (Chao et al., 2014; Hill, 1973).

The sensitivity of richness estimates to sequencing depth and sampling completeness was evaluated using both sample-size- and coverage-based approaches (Fig. S8). PCR replicates were rarefied to 10,000 reads for COI and 20,000 reads for 18S, and rarefied richness was compared with observed richness and sequencing depth. Sample-size- and coverage-based rarefaction and extrapolation curves were additionally generated using *iNEXT* to assess sampling completeness across sediment records and markers. These analyses were used to evaluate whether variation in observed richness was dominated by unequal sequencing depth rather than ecological differences among samples.

### Relationships between eukaryotic ASV richness and anthropogenic enrichment

Relationships between eukaryotic ASV richness and the Anthropogenic Enrichment Index (AEI) were evaluated separately for each sediment record and metabarcoding marker (COI and 18S). To facilitate pairing of biological and geochemical observations, CCO, COL, HIT and HIO were aggregated into 20-year intervals, whereas 30-year intervals were used for DEE, STO, CSD and CMB. Interval widths were selected according to differences in record duration and temporal sampling resolution, balancing the preservation of temporal detail with the need to obtain sufficient paired biological and geochemical observations. Median richness and AEI were calculated within each interval, and only intervals containing both variables were retained.

Associations across each complete record were first assessed using Spearman’s rank correlation coefficient, calculated with cor.test in R (Spearman, 1961). Their magnitude was also estimated using linear models after richness and AEI had been standardised within each site and marker. In these models (richness^z^ ∼ AEI^z^), the AEI coefficient represented the expected change in richness standard deviations associated with a one-standard-deviation increase in AEI. Models were fitted when at least five paired bins were available.

Because richness and AEI could both increase or decrease systematically through time, their association across a complete sediment record could partly reflect shared or opposing long-term trends rather than coupled variation around those trends. To distinguish these possibilities, a second set of models included standardised calendar year as a linear covariate predictor (richness^z^ ∼ AEI^z^ + Year^z^). In these models, the AEI coefficient estimates the association between richness and AEI after accounting for their average linear change through calendar time. Models were fitted when at least six paired temporal bins were available. To assess whether observations close in time remained correlated after fitting these fixed effects, generalised least-squares models were fitted as a sensitivity analysis using the same predictors and a continuous first-order autoregressive residual structure (gls and corCAR1 in the *nlme* package; Pinheiro et al., 1999). These models required at least eight paired bins and were therefore not fitted for CCO or COL (Table S8).

Directional changes in median richness and AEI were calculated between successive retained bins within each sediment record and marker. Transitions were classified into four categories: both variables increased, both decreased, richness decreased while AEI increased, or richness increased while AEI decreased. Given the temporal dependence of consecutive differences and the small number of transitions in some records (minimum = 4), transition frequencies and proportions were evaluated descriptively (Table S11).

Temporal variation in the richness and AEI association was further examined using centred rolling Spearman correlations calculated with rollapply in the *zoo* package (Zeileis & Grothendieck, 2005). Each window contained four consecutive bins, corresponding to approximately 80 years for records using 20-year bins and 120 years for those using 30-year bins. Correlations required at least three complete pairs and at least two distinct values for each variable. Because adjacent windows overlapped and the shortest records produced only two estimates, rolling correlations were treated as exploratory descriptions of temporal non-stationarity and were not subjected to individual significance tests.

Nominal two-sided p-values were reported for the complete-record Spearman correlations and regression coefficients, using p < 0.05 as the significance threshold. Effect sizes and 95% confidence intervals were reported where applicable, and associations were not interpreted as evidence of causality.

### Taxonomic richness profiles

Taxonomic composition and taxon-specific richness were analysed separately for the COI and 18S markers in R (v4.3.1) (R Core Team, 2025). For each sample, total eukaryotic ASV richness was calculated together with richness assigned to Fungi, Metazoa, Plantae, and protists. “Protists” was used as an operational grouping for eukaryotic ASVs assigned to protistan lineages but not to Fungi, Metazoa or Plantae and was not treated as a formal monophyletic taxon. ASVs assigned to Eukaryota but lacking a reliable kingdom-level classification were classified as unassigned eukaryotes.

Temporal profiles show sample-level ASV richness within each taxonomic category. Richness was represented using a pseudo-logarithmic scale to allow total richness and individual taxonomic groups to be visualised simultaneously; this transformation was used only for graphical representation. For each site and marker, pie charts show the percentage contribution of each taxonomic category to the pooled number of unique ASVs detected throughout the corresponding sediment record. Percentages therefore represent pooled ASV richness rather than relative read abundance.

Metazoan composition was examined using the same procedure. For each site and marker, the five phyla containing the greatest pooled number of unique metazoan ASVs were identified independently, and the remaining phyla were combined into an “Other Metazoa” category.

### Beta diversity among sediment records

Community composition patterns were evaluated separately for COI and 18S using three datasets: all retained ASVs irrespective of taxonomic assignment, ASVs assigned to Eukaryota, and ASVs assigned to Metazoa. Samples containing no ASVs after taxonomic filtering and ASVs absent from all retained samples were excluded. ASV counts were converted to sample-wise relative abundances with decostand(method = “total”), and Bray– Curtis dissimilarities were calculated with vegdist (method = “bray”) in the *vegan* package.

Community dissimilarities were visualised using non-metric multidimensional scaling (nMDS) implemented with metaMDS. Two- and three-dimensional solutions were compared, and the three-dimensional solutions were retained because they reduced stress by approximately 30% relative to the corresponding two-dimensional configurations. The main figures show projections onto NMDS1 and NMDS2, while the NMDS1–NMDS3 and NMDS2–NMDS3 projections were examined as complementary views. Reported stress values refer to the complete three-dimensional configurations.

Compositional differences among sediment records were tested using permutational multivariate analysis of variance (PERMANOVA), with Site as the explanatory variable, implemented with adonis2 and 9,999 permutations (Anderson, 2001). Homogeneity of multivariate dispersion was evaluated from distances to the spatial median using betadisper function, followed by permutest with 9,999 permutations (Anderson et al., 2006). PERMANOVA and dispersion results were interpreted together because significant differences in dispersion can contribute to apparent differences among groups.

Because each site was represented by a dated sediment core, the analyses describe compositional differences among the sampled records rather than providing a spatially replicated test of general site effects (Baños et al., 2026).

### Identification of temporally coincident multiproxy change

Periods of temporally coincident change were identified by determining whether multiple proxy groups showed marked trends within the same centred temporal window at each site: eukaryotic ASV richness, δ¹³C, δ¹⁵N, C:N ratio, and the general AEI. Original values were retained for graphical display, with COI richness multiplied by 10 for visualisation only. All trend calculations used unscaled values standardised within each site, proxy and analytical series.

Local temporal trends were estimated by regressing each standardised proxy series against calendar year within centred moving windows using lm in R. Windows of 25 years were used for CCO, COL, HIT and HIO, and 100-year windows for DEE, STO, CSD and CMB. Windows were moved in one-year increments across each record and were truncated at record boundaries. A regression was fitted only when at least three observations from three distinct years and at least two distinct standardised values were available.

For each regression, the slope was multiplied by the corresponding window length to estimate local change over 25 or 100 years. A proxy was classified as undergoing marked change when the absolute estimated change was at least 0.25 within-series standard deviations. The sign of the estimate distinguished increasing and decreasing trajectories. The 0.25-standard-deviation threshold was defined a priori as an operational effect-size criterion rather than a general threshold of ecological significance.

A focal year was classified as a period of concurrent multivariate change when at least three of the five proxy groups met this criterion, irrespective of direction. Complete information for all five groups was not required, but at least three proxies needed valid local regressions and had to meet the effect-size threshold. Consecutive focal years assigned to the same category were grouped into continuous intervals.

Concurrent change was classified as potential anthropogenic intensification when AEI increased, richness decreased and at least one of δ¹³C, δ¹⁵N or C:N ratio also met the effect-size criterion. Potential anthropogenic relaxation was defined as decreasing AEI, increasing richness and marked change in at least one additional proxy. All remaining configurations meeting the three-proxy criterion were classified as other concurrent changes. No predefined direction was imposed on δ¹³C, δ¹⁵N or C:N ratio because their responses can indicate different organic-matter sources, trophic processes and local biogeochemical conditions.

Regression p-values were retained as diagnostic information but were not used to define marked change or concurrent intervals. The resulting categories were therefore interpreted as exploratory, effect-size-based descriptions of temporally coincident proxy configurations, rather than as evidence of causal relationships or simultaneous statistical significance.

## Results

### Sediment chronology and accumulation rates

Excess ^210^Pb profiles showed substantial variation in recent sediment accumulation rates among sediment cores (Table S4; Fig. S3). The lowest rates were estimated for STO and CSD (0.10 cm yr^-1^), DEE (0.11 cm yr^-1^), and CMB (0.15 cm yr^-1^). Below the unsupported ^210^Pb dating horizon estimated basal ages were of approximately 1134 CE for STO, 1135 CE for CSD, 1384 CE for DEE and 1419 CE for CMB. The estimates for STO and CSD were poorly constrained because each regression contained only two unsupported ^210^Pb observations (Table S4).

Higher accumulation rates were estimated for HIT (0.42 cm yr^-1^), CCO (0.53 cm yr^-1^), COL (0.63 cm yr^-1^), and HIO (0.63 cm yr^-1^). Unsupported ^210^Pb remained detectable almost to the bases of the 49-cm CCO and COL cores, which had basal ages of approximately 1939 and 1953 CE, respectively. The HIT and HIO cores had extrapolated basal ages of approximately 1812 and 1884 CE. The unsupported ^210^Pb dating horizon ranged from 3 cm in CSD, 10 cm in STO, 15 cm in DEE and 15.5 cm in CMB to 40 cm in HIT, 48 cm in CCO, 49 cm in COL and 65 cm in HIO (Fig. S3). Uncertainty in the estimated accumulation rates also varied among records, with particularly wide 95% ranges for CMB and HIO (Table S4).

AMS ^14^C determinations from the lower sections of five cores extended the chronological coverage beyond the unsupported ^210^Pb horizons (Table S5; Fig. S4). Calibrated ages are reported as 95.4% highest-probability-density ranges (conventionally, 2σ). The oldest dated material was recovered at 85 cm in CSD and yielded a calibrated age range predominantly between 933 and 809 Before Common Era (BCE). The basal DEE sample at 70 cm was dated to 424–564 CE, while the 85-cm samples from HIO and HIT were dated predominantly to 889– 995 CE and 1027–1159 CE, respectively. The CMB sample at 85 cm produced calibrated age ranges of 1305–1364 CE and 1383–1419 CE. For DEE, CSD, HIT, and HIO, the calibrated years were older than the basal ages predicted by extrapolating the recent CF:CS-derived accumulation rates, whereas the younger CMB age range overlapped the extrapolated basal age of approximately 1419 CE. Because the radiocarbon samples were located 20–82 cm below the corresponding unsupported-^210^Pb horizons, the two methods constrained different sections of the cores. Comparisons therefore relied on extrapolating the recent ^210^Pb-derived accumulation rates beyond their directly dated intervals and should be interpreted cautiously.

### Site-specific patterns of element enrichment

Site-specific element sets differed among sediment records (Fig. S5; Table S6). In the Ebro Delta, DEE retained Cr, Ni, P, Pb and Zn. Within the Santoña system, STO retained As, Cd, Cr, Hg, P, Pb and Sn, whereas the shorter CCO and COL records retained Cd, Hg, Ni and Pb, and Cd, Hg and P, respectively. The southern Atlantic records were also heterogeneous: CSD retained Hg and Pb; CMB retained Ni, Pb and Sn; HIT retained As, Cu, Hg, Ni, P, Pb and Zn; and HIO retained Cu, Hg, Ni, P, Pb and Sn.

Enrichment magnitude and recurrence did not always covary across elements (Fig. S5). Some elements showed high mean positive robust z-scores but exceeded the strong-enrichment threshold in relatively few observations, whereas others had lower mean positive scores but exceeded the threshold (z^robust^ > 3) more frequently. Magnitude and recurrence therefore provided complementary descriptions of element-specific enrichment, although element retention was based exclusively on the occurrence of strong enrichment in at least two samples.

### Sensitivity of the AEI to site-specific element selection

The general and site-specific AEIs showed broadly similar temporal patterns but differed in their absolute values (Fig. S6; Table S6). Across the 402 paired observations, the two indices had a strong rank association (ρ = 0.887), while their absolute concordance was lower (CCC = 0.623). Concordance increased to 0.913 after both indices were standardised within each site, showing greater agreement in temporal trajectories after differences in location and scale had been removed.

The mean site-specific AEI was higher than the mean general AEI (0.925 versus 0.623), with a mean paired difference of 0.302. Agreement was highest in HIO (mean difference approximately zero; CCC = 0.933) and HIT (mean difference = −0.014; CCC = 0.890), followed by STO (mean difference = 0.244; CCC = 0.857). The largest difference occurred in CMB, where the site-specific AEI exceeded the general AEI by an average of 1.123 units. Despite a strong rank association at this site (ρ = 0.928), absolute concordance was low (CCC = 0.411). Absolute concordance was also comparatively low in COL (CCC = 0.539), CSD (CCC = 0.581) and CCO (CCC = 0.634). CSD had the weakest rank association (ρ = 0.621), whereas DEE had the strongest (ρ = 0.964).

Relationships between richness and the element-specific Pb and Hg enrichment scores varied among sediment records and markers (Fig. S7; Table S9). At CMB, Pb was negatively associated with both COI and 18S richness in the unadjusted models, partly reproducing the negative relationships observed with the AEI, although both associations weakened when calendar year was included as an additional predictor. By contrast, neither Pb nor Hg reproduced the positive DEE–COI relationship identified using the AEI. Some additional element–richness coefficients had nominal p-values below 0.05, but none remained significant after correction for multiple testing. Overall, the analyses supported the absence of a consistent richness response to Pb or Hg across sediment records and indicated that the richness–AEI relationships could not generally be attributed to either element individually.

Overall, restricting the index to locally recurrent elements preserved the relative temporal patterns more consistently than their absolute magnitude.

### Sequencing output and data overview

High-throughput sequencing of the *seda*DNA metabarcoding libraries generated 1,377,919,936 raw reads across sediment samples and negative controls for both metabarcoding markers. Initial sequence inference recovered 549,798 ASVs for 18S and 632,966 ASVs for COI before completion of the quality-control, curation and contamination-filtering workflow.

Following quality filtering, denoising, read-pair merging, chimera removal, LULU curation, low-abundance filtering and control-based contaminant subtraction, the final 18S dataset contained 424,652 ASVs and 748,479,852 reads across 829 retained PCR replicates, representing 122 sediment horizons. The final COI dataset contained 270,136 ASVs and 399,543,516 reads across 832 retained PCR replicates, representing 118 sediment horizons. Mean sequencing depth was 902,871 reads per retained PCR replicate for 18S and 480,221 reads per retained PCR replicate for COI.

Rarefaction reduced sample-level ASV richness by an average of 23% for 18S and 43% for COI relative to the corresponding unrarefied datasets (Fig. S8). Nevertheless, observed and rarefied richness remained strongly correlated for both markers (18S: Spearman’s ρ = 0.954; COI: ρ = 0.856). Sequencing depth explained a relatively small proportion of variation in rarefied richness (18S: R^2^ = 0.076; COI: R^2^ = 0.048). Sample-size-based rarefaction curves approached an asymptote for most samples, and coverage-based estimates indicated generally high, although variable, sampling completeness across sites and markers (Fig. S8).

Before control-based subtraction, negative controls contained a mean of 385 reads for COI (SD = 408) and 519 reads for 18S (SD = 1,430), substantially fewer than sediment samples. Reads detected in controls were incorporated into the ASV-specific maximum-subtraction procedure. Overall, the 18S *seda*DNA dataset recovered more ASVs and greater sample-level richness than the corresponding COI dataset.

### Temporal biodiversity and anthropogenic enrichment dynamics

Temporal trajectories of eukaryotic ASV richness and anthropogenic enrichment varied among sediment records and between metabarcoding markers (Fig. 1). Analyses of median values within site-specific temporal bins included between five and fifteen paired observations per site and marker (Table S7).

**Fig. 1:**
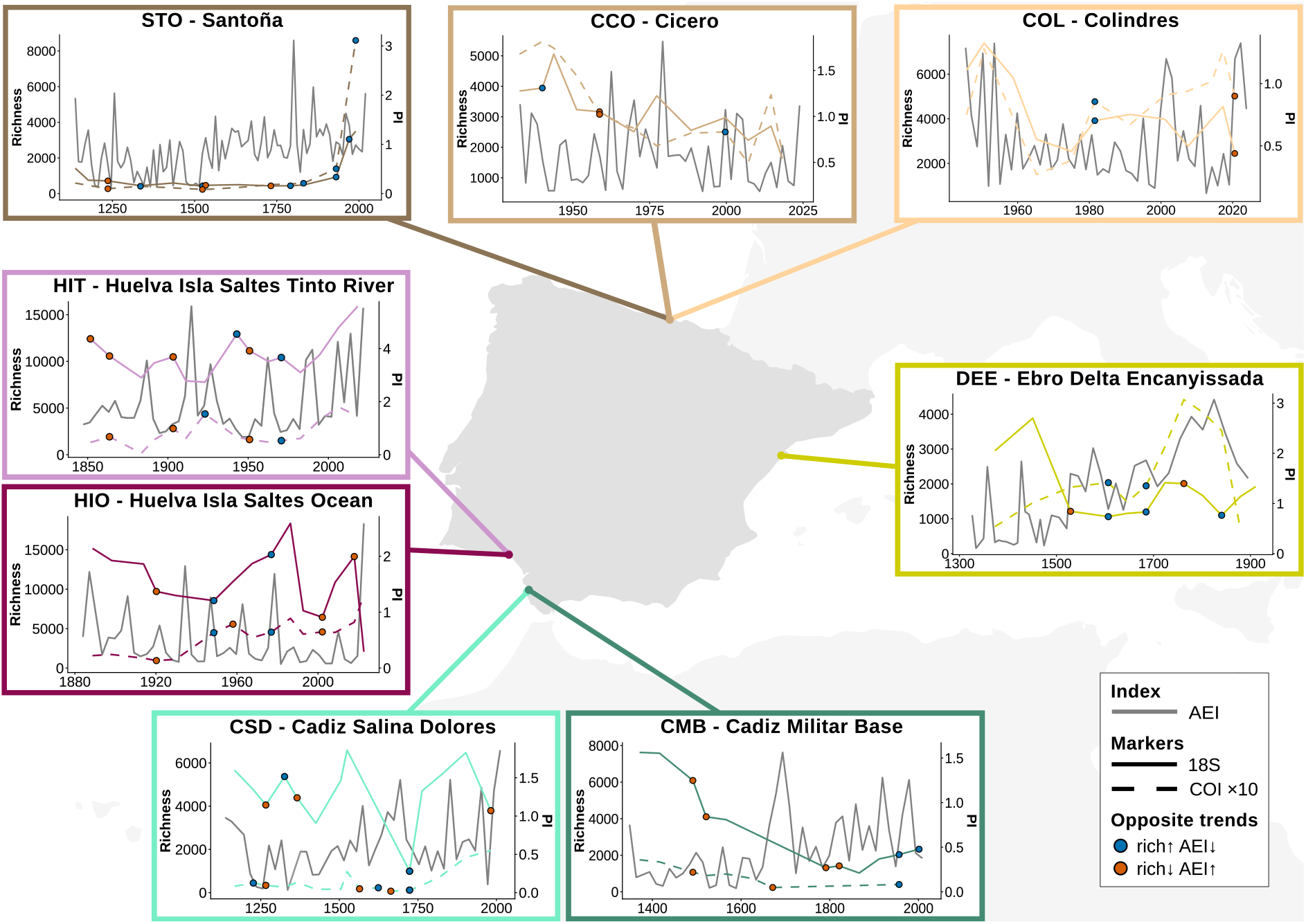
Geographical location of the eight coastal wetlands across the Iberian Peninsula with the temporal trajectories of sedimentary ancient DNA (sedaDNA) richness and the general Anthropogenic Enrichment Index (AEI). The northern Atlantic sites comprise Cicero (CCO), Colindres (COL), and Santoña (STO); the southern Atlantic sites comprise Saltés Island (HIT and HIO) and the Bay of Cádiz (CMB and CSD); and the Mediterranean site is located in the Ebro Delta (DEE). Richness was calculated independently for 18S and COI as the number of detected eukaryotic ASVs (Hill number q = 0). The general AEI was calculated as the equal-weight mean of positive robust enrichment scores across the ten candidate elements with available XRF measurements and is shown as a dotted black line. COI richness was multiplied by 10 for visualisation only. All statistical analyses used the original values. Filled circles mark richness values associated with consecutive-bin transitions in which richness and AEI changed in opposite directions (orange: richness decreased while AEI increased; blue: richness increased while AEI decreased).

The clearest negative relationships occurred at CMB, where richness was negatively correlated with AEI for both 18S (ρ = −0.654, p = 0.008) and COI (ρ = −0.718, p = 0.003). Standardised linear models produced corresponding negative slopes for 18S (β = −0.568, 95% CI: −1.061 to −0.075, p = 0.027) and COI (β = −0.613, 95% CI: −1.086 to −0.140, p = 0.015). DEE showed the opposite pattern for COI. Richness was positively correlated with AEI (ρ = 0.718, p = 0.013), and the standardised linear coefficient was also positive (β = 0.752, 95% CI: 0.255 to 1.249, p = 0.008). The corresponding 18S relationship was negative but not significant (ρ = −0.236, p = 0.484; β = −0.548, p = 0.081). Additional marker-specific relationships were detected by the linear models at HIT, where COI richness increased with AEI (β = 0.640, 95% CI: 0.061 to 1.220, p = 0.034), and at HIO, where 18S richness decreased with AEI (β = −0.760, 95% CI: −1.409 to −0.110, p = 0.029). The corresponding Spearman correlations were not significant at either HIT (ρ = 0.546, p = 0.083) or HIO (ρ = 0.095, p = 0.823). No significant relationships were detected at CCO, COL, CSD or STO.

The preceding models used dated observations but related richness only to AEI. We therefore fitted a second set of models that included calendar year as an additional predictor, testing whether richness remained associated with AEI after considering the linear change of both variables through time. In these models, only the positive DEE–COI relationship remained significant (β = 1.093, 95% CI: 0.139 to 2.047, p = 0.030; Table S8). The coefficients were no longer significant for CMB–18S (β = −0.308, p = 0.096), CMB–COI (β = −0.395, p = 0.056), HIT–COI (β = 0.514, p = 0.068) or HIO–18S (β = −0.622, p = 0.058). Models incorporating a continuous first-order autoregressive residual structure produced effectively identical estimates. Time-adjusted and autoregressive models were not fitted for CCO or COL because only five paired bins were available.

Overall temporal trends also differed among records and markers (Table S10). AEI increased significantly through time only at DEE (β = 0.461, 95% CI: 0.242–0.680, p = 0.001). Richness declined for both markers at CCO (18S: β = −3.099, p = 0.003; COI: β = −2.970, p = 0.018) and CMB (18S: β = −0.436, p < 0.001; COI: β = −0.394, p = 0.003), whereas COI richness increased at CSD (β = 0.230, p = 0.020) and HIO (β = 1.777, p = 0.005). No other overall linear trends were significant. The CCO coefficients were based on only five temporal bins and should therefore be interpreted cautiously.

Richness and AEI changed in opposite directions in 66 of 147 consecutive-bin transitions (44.9%; Table S11). The proportions were similar for 18S (34 of 76; 44.7%) and COI (32 of 71; 45.1%). Opposite-direction transitions were most frequent for 18S in COL (75.0%), HIT (60.0%) and for COI in CMB (57.1%), although the COL estimate was based on only four transitions.

Rolling correlations alternated between positive and negative values in most site–marker combinations (Fig. S9). DEE–COI was the main exception, with positive correlations across all available windows. Because the four-bin windows overlapped and the shortest records produced only two estimates, these profiles were treated as exploratory descriptions of temporal variation in the richness–AEI relationship.

### Taxonomic composition and taxon-specific richness patterns

The taxonomic distribution of ASV richness reconstructed from *seda*DNA differed between markers and among sediment records (Fig. 2). In the COI dataset, Fungi represented the largest proportion of pooled ASV richness in DEE, STO, CCO, COL, CSD and CMB, accounting for 28–52% of the unique ASVs in these records. Plantae was the largest component in HIT and HIO, accounting for 32% and 30%, respectively. Metazoa, protists and unassigned eukaryotes contributed smaller and more variable proportions.

**Fig. 2:**
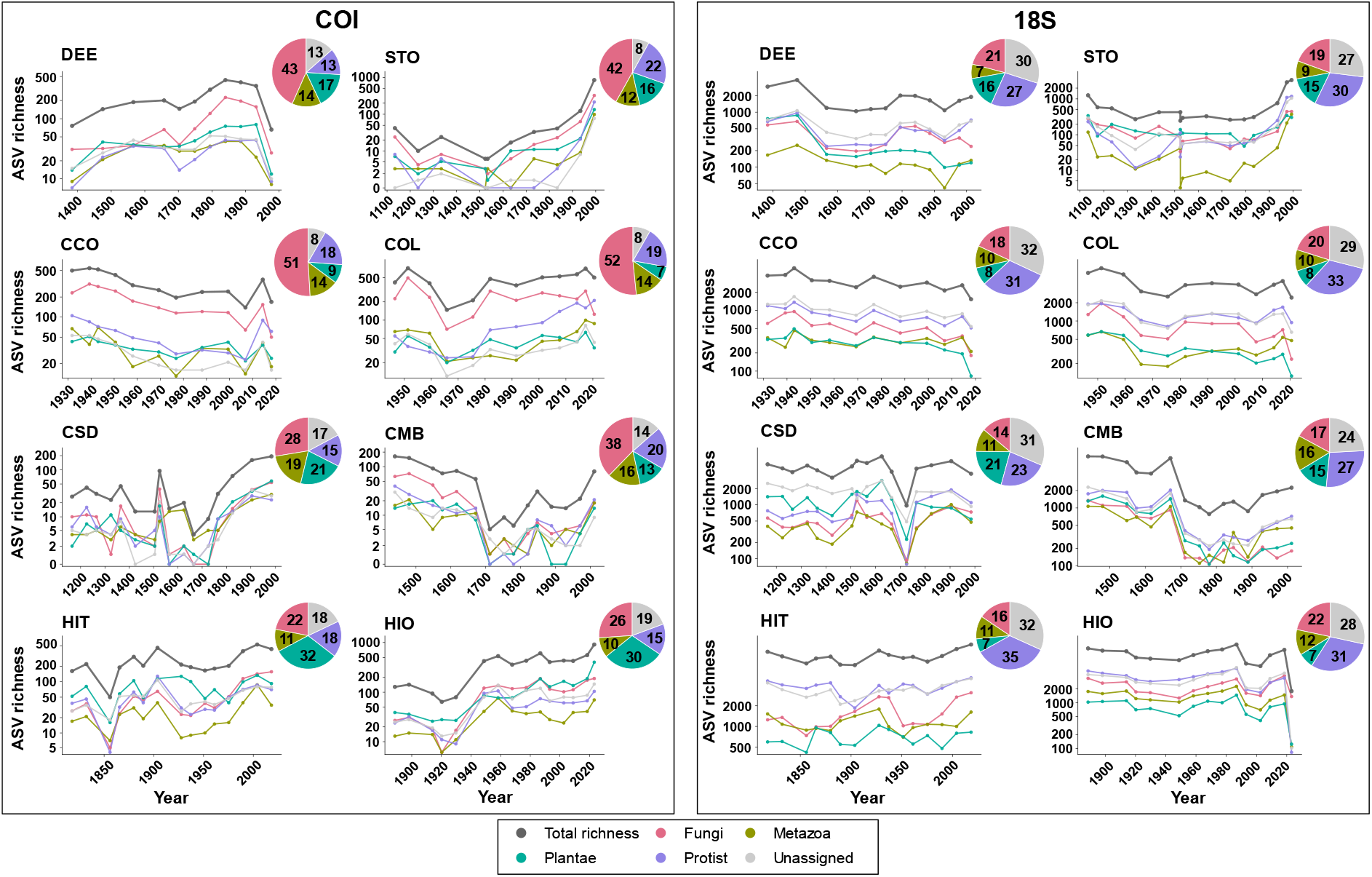
Kingdom-level distribution and temporal patterns of eukaryotic ASV richness reconstructed from sedimentary DNA across Iberian coastal wetlands. Panels show total eukaryotic ASV richness and richness assigned to Fungi, Metazoa, Plantae, protists and unassigned eukaryotes for COI and 18S at each site. Richness is displayed on a pseudo-logarithmic scale to allow total richness and less ASV-rich groups to be visualised simultaneously. All records are plotted against the calendar year. Pie charts show the percentage contribution of each taxonomic category to the pooled number of unique ASVs detected for the corresponding site and marker. Percentages do not represent relative read abundance. Colours identify taxonomic categories and are maintained across sites and markers.

The 18S dataset showed a different distribution. Protists and unassigned eukaryotes were the two largest components in every record, representing 23–35% and 24–32% of pooled richness, respectively. Fungi accounted for 14–22%, while Plantae and Metazoa generally represented smaller fractions. Total and kingdom-specific richness varied through time, and changes in total richness frequently coincided with changes in more than one kingdom-level group. Individual groups did not, however, follow identical trajectories across all records or periods.

Metazoan richness was also distributed differently between markers and sites (Fig. S10). Arthropoda was the largest component of pooled COI metazoan richness in six records, ranging from 26.8% in DEE to 54.5% in HIT; Mollusca was the largest component in STO and Annelida in COL. In the 18S dataset, Nematoda was the largest identified component in DEE, STO, CCO, COL, CMB and HIT, whereas the pooled “Other Metazoa” category was marginally largest in CSD and HIO. The five most ASV-rich phyla collectively accounted for approximately 74–97% of pooled metazoan richness, depending on the record and marker. Their temporal trajectories frequently varied alongside total metazoan richness, although the contribution of individual phyla was not constant through time.

### Community composition and beta diversity

Community composition differed among sediment records for both markers (Table S12; Fig. 3). For the complete ASV datasets, ordination stress decreased from 0.192 in two dimensions to 0.133 in three dimensions for COI and from 0.175 to 0.123 for 18S. The three-dimensional nMDS configurations showed compositional structuring among records, with differing degrees of overlap and within-record dispersion.

**Fig. 3:**
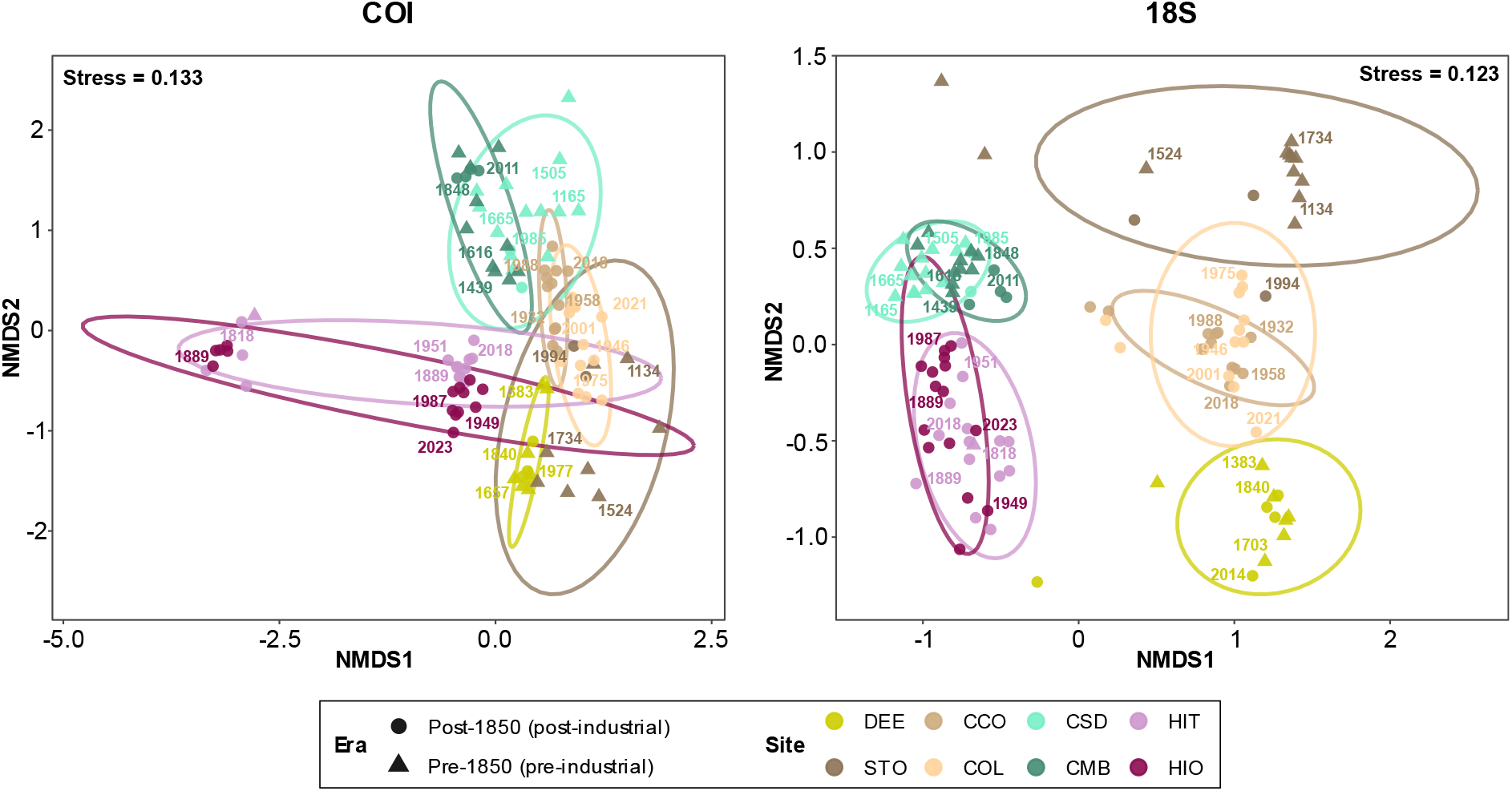
Community composition across Iberian sediment records reconstructed using COI and 18S metabarcoding. Non-metric multidimensional scaling (nMDS) ordinations were calculated from Bray–Curtis dissimilarities of sample-wise relative ASV abundances, including all retained ASVs irrespective of taxonomic assignment. Ordinations were fitted separately for COI (left) and 18S (right) in three dimensions and are shown as projections onto NMDS1 and NMDS2. Points represent individual sediment samples, colours identify sediment records, and point shapes distinguish pre-industrial (pre-1850) and post-industrial (post-1850) samples. Labels show four representative calendar years distributed across each record. Coloured lines are 95% data ellipses calculated under a multivariate t-distribution and summarise within-record dispersion. Reported stress values refer to the complete three-dimensional configurations.

PERMANOVA detected differences among records for both COI (pseudo-F = 4.652, R² = 0.253, p < 0.001) and 18S (pseudo-F = 5.970, R² = 0.291, p < 0.001). Site accounted for approximately 25% and 29% of compositional variation, respectively. Multivariate dispersion also differed among records for COI (F = 3.163, p = 0.005) and 18S (F = 3.377, p = 0.003); the PERMANOVA results therefore include potential contributions from both differences in centroid location and within-record dispersion.

Restricting the datasets to taxonomically assigned eukaryotes produced similar results (Table S12; Fig. S11A). Three-dimensional nMDS stress was 0.164 for COI and 0.120 for 18S. Site accounted for 23.0% of compositional variation in COI (pseudo-F = 4.091, p < 0.001) and 31.0% in 18S (pseudo-F = 6.562, p < 0.001). Dispersion differed among records for both COI (F = 2.137, p = 0.046) and 18S (F = 4.121, p < 0.001).

Among-record differentiation was weaker in the Metazoa datasets (Table S12; Fig. S11B). Three-dimensional nMDS stress was 0.183 for COI and 0.143 for 18S. Site accounted for 15.4% of variation in COI (pseudo-F = 2.442, p < 0.001) and 17.9% in 18S (pseudo-F = 3.179, p < 0.001). No difference in multivariate dispersion was detected for COI (F = 0.997, p = 0.445), whereas dispersion differed among records for 18S (F = 5.679, p < 0.001). The 18S PERMANOVA result may therefore reflect both compositional location and within-record variability, while no corresponding dispersion effect was detected for COI.

### Multiproxy sediment trajectories and concurrent change

Multiproxy trajectories integrating *seda*DNA-derived richness, stable-isotope composition, C:N and AEI varied in timing, magnitude and direction among sediment records (Fig. 4). Periods of marked change were detected in every record, but the proxies involved and their directional configurations differed among sites.

**Fig. 4:**
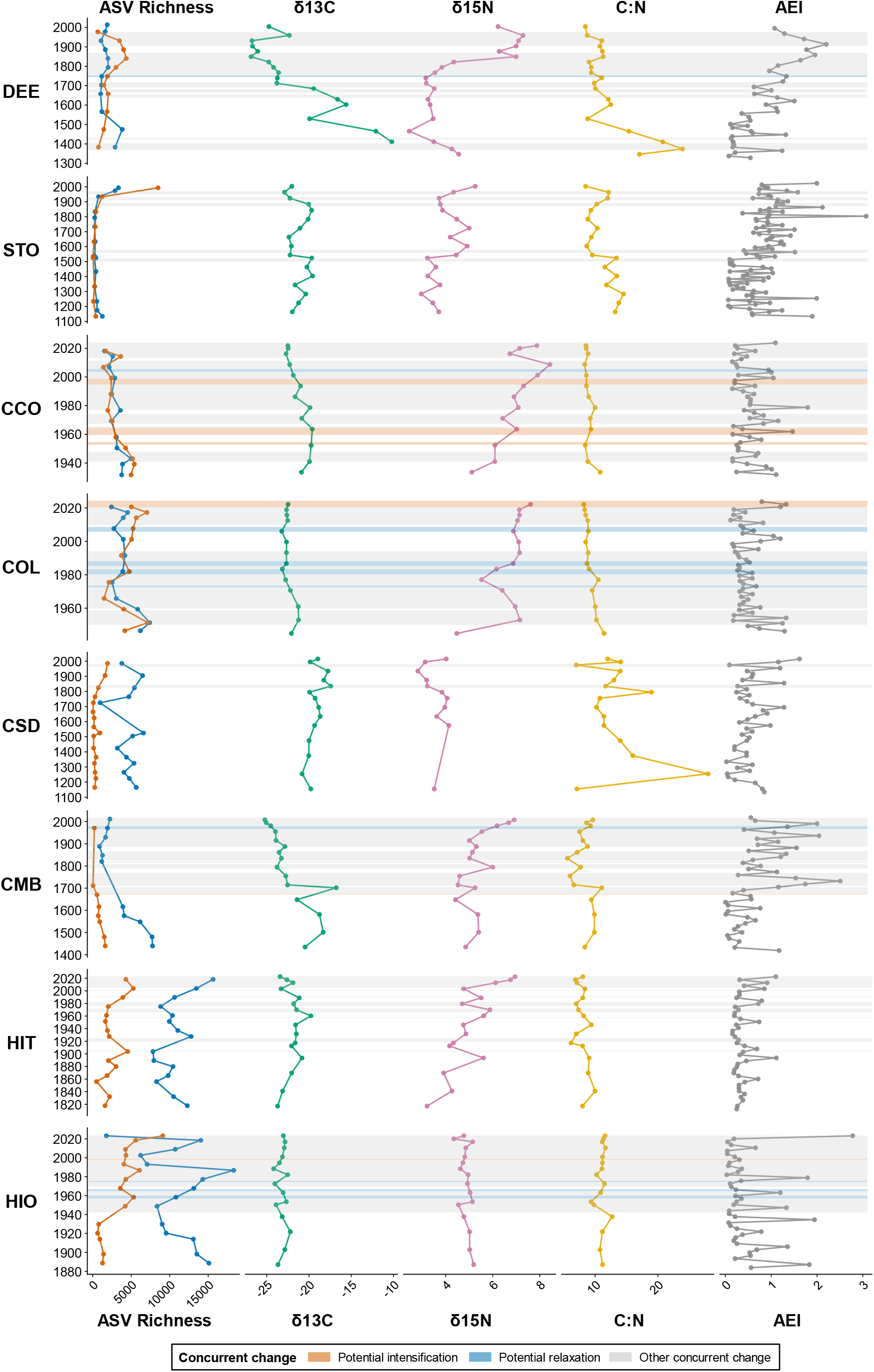
Multiproxy temporal trajectories and intervals of concurrent multivariate change across the eight wetland study sites. Rows represent individual sediment records, and columns show eukaryotic ASV richness, sedimentary δ¹³C, δ¹⁵N, C:N ratio and the general Anthropogenic Enrichment Index (AEI). Points represent individual observations, and connecting lines show their temporal trajectories. Within the richness panels, 18S richness is shown in blue and COI richness in orange; COI values were multiplied by 10 for visualisation only. Orange shading identifies potential intensification, defined as increasing AEI, decreasing ASV richness and marked change in at least one additional proxy. Blue shading identifies potential relaxation, defined as decreasing AEI, increasing richness and marked change in at least one additional proxy. Grey shading identifies other intervals in which at least three proxy groups met the effect-size criterion but the AEI–richness configuration did not satisfy either directional definition. Marked change was defined as an absolute estimated change of at least 0.25 within-series standard deviations. Local trends were calculated using 25-year moving windows for CCO, COL, HIT and HIO and 100-year windows for DEE, STO, CSD and CMB. Original proxy values are displayed on proxy-specific horizontal scales, and calendar year on site-specific vertical scales.

A total of 63 intervals met the operational criterion for concurrent multiproxy change (Table S13). Most were classified as other concurrent changes (47 intervals; 74.6%), indicating that at least three proxy groups showed marked local change without matching either predefined AEI–richness configuration. Six intervals (9.5%) were classified as potential intensification and ten (15.9%) as potential relaxation. The directions of change in δ¹³C, δ¹⁵N and C varied among sites and intervals.

Potential intensification was identified in four sediment records. Three intervals occurred at CCO (1952–1953, 1959–1963 and 1995–1998), while one occurred at CMB (1665–1666), HIO (1999) and COL (2021–2024). Each combined increasing AEI, decreasing richness and marked change in at least one additional proxy, although the corresponding stable-isotope and C directions differed. The CMB interval pre-dated industrialisation and is therefore reported only as an effect-size-defined directional configuration, not as evidence of industrial disturbance.

Potential relaxation was identified at DEE (1745–1753), CCO (2004–2005) and CMB (1968– 1979). Four intervals occurred at COL (1972, 1980–1982, 1985–1987 and 2006–2008), and three occurred at HIO (1957–1959, 1965–1966 and 1975). These intervals combined decreasing AEI, increasing richness and marked change in at least one additional stable-isotope or C proxy.

The number and classification of intervals also differed among records. CCO and COL each contained 11 intervals, CMB and HIO each contained nine, DEE and HIT each contained eight, STO contained five and CSD contained two. CCO, COL, CMB and HIO included both intensification and relaxation configurations; DEE contained one relaxation interval and seven other-change intervals. All intervals identified at STO, CSD and HIT were classified as other concurrent changes.

Overall, concurrent-change intervals were predominantly characterised by site-specific combinations of increasing and decreasing *seda*DNA richness, AEI and isotopic or biogeochemical proxies, rather than by a uniform regional sequence.

## Discussion

Our multiproxy sedimentary records reveal that long-term biodiversity patterns across Iberian coastal wetlands cannot be characterised by a uniform relationship between ASV richness and sedimentary elemental enrichment. Richness–enrichment associations differed among sediment records and metabarcoding markers, and most associations detected across complete records weakened after accounting for calendar time. Exploratory rolling correlations further indicated that the direction of these relationships could vary across historical periods within individual records. In parallel, COI and 18S recovered contrasting taxonomic profiles, while community composition differed markedly among sediment records. Geochemical, isotopic and biodiversity proxies also showed locally specific configurations of change. Together, these findings highlight the importance of temporal context, taxonomic coverage and environmental history when interpreting long-term biodiversity change, and demonstrate the limitations of inferring ecological responses from richness and sedimentary enrichment alone.

### Richness responses to anthropogenic enrichment varied among wetlands and through time

No common richness response to anthropogenic enrichment emerged across the eight sediment records. At CMB in Cádiz, higher AEI values were associated with lower richness for both 18S and COI, whereas the Ebro Delta record (DEE) showed a positive COI–AEI relationship. Marker-specific associations were also detected in Huelva, with a positive HIT– COI relationship and a negative HIO–18S relationship (Fig. 1; Tables S7 and S8). When calendar year was included as an additional predictor, however, only the positive DEE–COI association remained statistically significant. This indicates that several of the initial richness– AEI associations reflected shared or opposing linear trends through time rather than relationships independent of those trends (Li et al., 2019). The HIT–COI and HIO–18S associations were also not detected by the corresponding Spearman correlations and should therefore be interpreted as record-specific tendencies rather than strong evidence of enrichment-related biodiversity change, particularly given the limited number of temporal bins (Jenkins & Quintana-Ascencio, 2020).

The contrasting relationships observed among wetlands are not necessarily unexpected. Coastal ecosystems differ in hydrology, habitat structure, initial community composition and exposure history, all of which can influence how biodiversity responds to environmental change (Cloern, 2001; Mozdzer et al., 2020). Under some conditions, enrichment may favour opportunistic or disturbance-tolerant taxa, whereas under others, particularly under more intense or persistent exposure, it may contribute to the loss of sensitive taxa or the simplification of communities (King & Richardson, 2007; Odum et al., 1979). These changes in community composition need not produce equivalent changes in richness (Dornelas et al., 2014; Hillebrand et al., 2018; Suzzi et al., 2023). The AEI should therefore be interpreted as a relative measure of multielement enrichment rather than as a direct measure of nutrient availability, contaminant toxicity or ecological risk.

Temporal variability further characterised the richness-AEI relationships. Nearly half of all successive retained-bin transitions showed richness and AEI changing in opposite directions, and rolling correlations frequently switched between positive and negative values through time (Table S11; Fig. S9). DEE-COI was the main exception, retaining positive correlations throughout the available windows. Together, these patterns suggest that biodiversity responses to anthropogenic enrichment were not constant through time, even within individual wetlands.

The results indicate that similar levels of anthropogenic enrichment were associated with richness declines, richness increases, or little detectable change depending on the wetland, marker, and period considered. Stable or increasing *seda*DNA richness also does not necessarily indicate ecological stability because substantial taxonomic turnover can occur without a net change in richness (Dornelas et al., 2014; Hillebrand et al., 2018). The compositional differentiation among records (Fig. 3; Table S12) further suggests that local environmental histories played an important role in shaping long-term biodiversity trajectories across Iberian coastal wetlands.

### Spatially and taxonomically heterogeneous biodiversity change

Biodiversity patterns differed markedly among regions, taxonomic groups and metabarcoding markers (Fig. 2). In the COI records, Plantae contributed the largest share of pooled COI ASV richness in Huelva (HIT and HIO), whereas Fungi contributed the largest share in Santoña, Cádiz and the Ebro Delta. By contrast, protists and unassigned eukaryotes dominated the 18S datasets across all records. Marker-dependent differences were also evident within Metazoa: Arthropoda contributed most strongly to COI richness at most sites, whereas Nematoda predominated in most 18S records (Fig. S10). These contrasts partly reflect differences in primer affinity, marker resolution, gene-copy number and reference-database coverage (Casey et al., 2021; Clarke et al., 2017; van der Loos & Nijland, 2021). COI and 18S should therefore be regarded as complementary molecular records of different components of eukaryotic diversity rather than as interchangeable measures of the same community (Atienza et al., 2020; Casey et al., 2021; Sawaya et al., 2019; Zimmermann et al., 2024).

Community composition was also strongly differentiated among sediment records for both markers, including when the analysis was restricted to taxonomically assigned eukaryotes (Fig. 3; Table S12). This spatial structuring is consistent with the environmental heterogeneity of coastal wetlands, where salinity, tidal exchange, sediment characteristics, catchment inputs and habitat histories can vary over short distances (Dalrymple et al., 1992; Lorrain-Soligon et al., 2023). Because each locality was represented by a single core, however, these results describe differentiation among the sampled sediment records rather than a spatially replicated general site effect.

Differentiation persisted within Metazoa, although it was weaker than in the complete eukaryotic datasets (Fig. S11; Table S12), indicating that non-metazoan and unassigned ASVs contributed substantially to the broader spatial patterns. Together with the contrasting taxon-specific richness trajectories, these compositional differences show that long-term biodiversity change cannot be represented by total richness alone. Sediment samples with comparable ASV richness may contain compositionally distinct assemblages, and substantial taxonomic replacement can occur without a corresponding net loss of richness (Dornelas et al., 2014; Hillebrand et al., 2018). The patterns are consistent with heterogeneous community reorganisation across wetlands and taxonomic groups, rather than a uniform decline in biodiversity. Combining COI and 18S with both richness and compositional analyses provides a broader reconstruction of this ecological change than any single marker or diversity metric.

### Regional environmental histories shaped contrasting ecological trajectories

The multiproxy records show that biodiversity change occurred within distinct regional environmental histories rather than along a common trajectory of anthropogenic intensification (Fig. 4; Table S13). Periods of concurrent change were identified in all records, but most involved combinations of richness, elemental enrichment, stable isotopes and C:N that did not match the predefined intensification or relaxation configurations. The ecological meaning of these changes must therefore be interpreted in relation to the depositional setting, biological community and historical pressures represented by each wetland.

Within the Santoña estuary, STO, CCO and COL followed different multiproxy trajectories despite their geographical proximity. This heterogeneity is consistent with the history of salt-marsh reclamation, agricultural occupation and subsequent regeneration documented in the estuary (García-Artola et al., 2016). COL is located beside a dyke that was absent from the 1954 edition of the Spanish National Topographic Map but present by 2003, placing its construction within the second half of the twentieth century (Instituto Geográfico Nacional, MTN50 sheet 36). Altered tidal exchange and sediment delivery associated with this landscape modification provide a plausible context for the observed changes, although neither the chronological resolution nor the proxy evidence permits their attribution to the dyke itself. In Cádiz, both CSD and CMB recorded increasing elemental enrichment from approximately 1500 CE, broadly coinciding with the growing role of the Bay of Cádiz within the port network of the Carrera de Indias and with increasing maritime and military activity during the sixteenth and seventeenth centuries (Acedo Romero, 1988; Gullón Abao & Reyes, 2023; Hammer, 1997; Morales-Padrón, 2021). Nevertheless, their biological trajectories differed: CMB showed declining richness and negative richness–AEI relationships, whereas CSD did not show an equivalent response. This contrast indicates that a shared regional history of increasing human pressure did not produce the same ecological trajectory at neighbouring locations.

The Huelva records provide a further example of ecological divergence within the same estuarine system. HIT, facing the inner Tinto estuary, contained only other concurrent-change configurations, whereas HIO, located on the ocean-facing side of Saltés Island, contained both potential intensification and relaxation intervals during the twentieth century. Construction of the Juan Carlos I breakwater, completed in 1981, substantially modified sediment transport and coastal morphology around the mouth of the Tinto–Odiel estuary (Morales et al., 2004). Some of the multiproxy changes at HIO occurred broadly within this period of port expansion, although their temporal correspondence does not demonstrate that the infrastructure caused the biological or geochemical changes.

The Ebro Delta followed a different trajectory from the Atlantic records, with a persistent positive relationship between COI richness and the AEI. Intensive rice cultivation and the discharge of nutrient-rich agricultural drainage have altered nutrient dynamics in the delta and adjacent coastal waters (Calvo-Cubero et al., 2014; Cruzado et al., 2002). Phosphorus, an essential nutrient and an important contributor to the enrichment signal at DEE, may have increased resource availability and favoured opportunistic or disturbance-tolerant taxa. This provides a plausible ecological context for the positive COI richness–AEI relationship, although the AEI integrates several elements and sedimentary P does not directly represent bioavailable phosphorus. The observed association therefore cannot be attributed to nutrient enrichment alone.

Comparison of the general and site-specific AEIs showed that element selection affected the absolute magnitude of the index more strongly than its relative temporal patterns (Fig. S6; Table S6). The similarity between these patterns indicates that the main enrichment trajectories were not highly sensitive to whether a fixed or locally selected element set was used, although their magnitude remained dependent on element selection and reference conditions (Abrahim & Parker, 2008; Greco et al., 2019; Kowalska et al., 2018). The AEI should therefore be interpreted as a relative signal of multielement enrichment, rather than as a measure of contaminant concentration, bioavailability, toxicity or ecological risk. Accordingly, richness–AEI relationships represent historical associations within individual wetlands, not uniform dose–response relationships.

Together, these regional comparisons show that anthropogenic enrichment formed part of broader ecological reorganisations whose expression depended on local environmental history. Concurrent shifts in molecular, isotopic and geochemical proxies indicate periods of environmental change, but their direction and biological consequences differed among and within regions. The multiproxy records therefore support the central conclusion that coastal-wetland biodiversity change was historically contingent rather than a uniform response to an equivalent anthropogenic pressure.

### Chronological and methodological constraints

Several limitations should be considered when interpreting the reconstructed biodiversity trajectories. Each locality was represented by a single sediment core, providing a temporal record for regional comparison but not spatial replication within the corresponding wetland. Individual cores cannot capture heterogeneity in sedimentary conditions or DNA deposition across the wider wetland and its catchment (Baños, Knudsen, et al., 2026). Differences among records may therefore arise from local depositional and preservation conditions as well as broader ecological differences (Boere et al., 2011).

Differences in sediment accumulation and chronological uncertainty may also affect the estimated timing of ecological change and comparisons among records. This is particularly relevant below the unsupported ^210^Pb horizons, where ages were obtained by extrapolating recent accumulation rates. The deeper radiocarbon determinations provided independent chronological information but were not incorporated into the primary age–depth models. Changes in sediment accumulation, erosion and mixing can challenge the assumptions of ^210^Pb dating models in coastal environments (Abril, 2022; Andersen et al., 2000; Arias-Ortiz et al., 2018). The timing and duration of older changes should therefore be regarded as approximate rather than as precisely dated historical events.

*Seda*DNA records are additionally influenced by transport, sediment mixing, degradation and taxon-specific differences in preservation. Replicate sediment records can differ because of local depositional processes and sediment heterogeneity (Pearman et al., 2021). Detected DNA may originate from local organisms, the overlying water column or the surrounding catchment, while bioturbation and sediment reworking may alter its temporal resolution (Capo et al., 2021; Corinaldesi et al., 2011; Giguet-Covex et al., 2019; Nguyen et al., 2023). Sedimentary ASV richness should therefore be interpreted as a molecular proxy for past biodiversity rather than as a direct census of the organisms present at a particular place and time. Marker-specific amplification and taxonomic-assignment biases further influence the components of biodiversity detected by COI and 18S, reinforcing their interpretation as complementary rather than equivalent signals (Casey et al., 2021; Zimmermann et al., 2024).

Despite these constraints, high sequencing depth, the strong correspondence between observed richness and sampling-standardised richness estimates, and the recovery of compositional differentiation with both markers support the internal consistency of the principal patterns (Figs. S8 and 3). This robustness does not consist of identical biodiversity responses across records; rather, both markers provide evidence of spatially and taxonomically heterogeneous trajectories. Although *seda*DNA does not provide a direct census of former communities, it extends biodiversity observations beyond conventional ecological monitoring. Its integration with chronological, geochemical and isotopic evidence therefore provides long-term context for interpreting ecological change while acknowledging uncertainty in preservation, chronology and local depositional conditions.

### Conclusions

Our reconstruction of biodiversity change across eight Iberian coastal wetlands reveals contrasting long-term relationships between eukaryotic ASV richness and sedimentary multielement enrichment. These relationships differed among sediment records and metabarcoding markers, and most associations weakened after accounting for calendar time. Contrasting taxonomic profiles and marked differences in community composition among records further show that richness alone provides an incomplete representation of the biodiversity patterns preserved in coastal sediments. The integration of sedaDNA, geochemical and isotopic records also revealed locally specific configurations of environmental change rather than a common regional trajectory. Together, these findings demonstrate the value of historical multiproxy records for placing biodiversity change in its temporal and environmental context. They also underscore the importance of considering taxonomic composition, multiple biological groups and locally informed historical reference conditions when assessing ecological change and developing conservation strategies for coastal wetlands.

## CRediT (Contribution Roles Taxonomy)

**Elena Baños:** Conceptualization (equal); writing – original draft (lead); formal analysis (lead); data curation (lead); investigation (equal); methodology (lead); software (lead); visualization (lead); writing – review and editing (equal). **Sandra Nogué:** Conceptualization (equal); writing – original draft (supporting); funding acquisition (equal); project administration (equal); investigation (equal); resources (equal); supervision (lead); validation (equal); methodology (supporting); writing – review and editing (equal). **Javier de la Casa:** Conceptualization (supporting); writing – original draft (supporting); methodology (supporting); validation (equal); writing – review and editing (equal). **Marc Rius:** Funding acquisition (equal); project administration (equal); investigation (equal); resources (equal); writing – review and editing (equal). **Luke E. Holman:** Supervision (supporting); writing – review and editing (equal). **Andrew B. Cundy:** Investigation (supporting); methodology (supporting); resources (equal); writing – review and editing (equal). **Bastian Hambach:** Resources (equal); methodology (supporting); writing – review and editing (equal). **Enrique González-Ortegón:** Investigation (supporting); writing – review and editing (equal). **Erik J. de Boer:** Investigation (supporting); writing – review and editing (equal). **Jesús Zarcero:** Investigation (supporting); writing – review and editing (equal). **Xavier Benito:** Investigation (supporting); writing – review and editing (equal). **Xavier Turon:** Investigation (supporting); writing – review and editing (equal).

## Acknowledgments

This work was supported by the TEMPOINVASIONS grants TED2021-132228B-C21 and TED2021-132228B-C22, both funded by MCIN/AEI/10.13039/501100011033 and the European Union (“NextGenerationEU”/PRTR). We are also grateful to the Spanish Ministry of Science, Innovation and Universities for providing financial support with the BlueDNA PID2023-146307OB-C22 grant (MICIU/AEI/https://doi.org/10.13039/501100011033 and FEDER, UE). Pawel Gaca and Madeleine Cobbold at GAU-Radioanalytical, University of Southampton, are thanked for conducting ^210^Pb dating analyses. We thank Megan Wilding from the National Oceanography Centre Southampton in University of Southampton, for conducting the isotope stable analysis. We thank Antonio Moreno Muñoz, technician at the Institute of Marine Sciences of Andalusia (ICMAN-CSIC), for his assistance and collaboration during fieldwork in Cádiz and Huelva. Marc Rius and Xavier Turon are members of the research group SGR2021-00405 and Sandra Nogué is member of the research group SGR 01333, both funded by the Generalitat de Catalunya. Luke E. Holman is supported by the European Research Council through the SeaChange project (Grant agreement No. 856488).

## Data Availability Statement

Raw COI and 18S metabarcoding sequence data are available through the NCBI Sequence Read Archive under BioProject PRJNA1535453. The processed datasets, metadata, supporting tables and R scripts used to reproduce the statistical analyses and figures are publicly available on GitHub and archived on Zenodo under DOI 10.5281/zenodo.22646988.

## Supplementary Information

**Table S1:** Sampling locations and core characteristics for sedimentary DNA (*seda*DNA) analyses. Geographic coordinates, site names, and total core depths are reported for each sampling location.

| **Core ID** | **Site** | **Latitude** | **Longitude** | **Core Depth (cm)** |
|---|---|---|---|---|
| DEE | Ebro Delta Encanyissada | 40.653108 N | 0.685247 E | 70 |
| STO | Santoña | 43.446528 N | -3.466719 W | 90 |
| CCO | Cicero | 43.422497 N | -3.481806 W | 50 |
| COL | Colindres | 43.400247 N | -3.455278 W | 50 |
| CSD | Cadiz Salina Dolores | 36.464889 N | -6.223167 W | 90 |
| CMB | Cadiz Militar Base | 36.499139 N | -6.193056 W | 90 |
| HIT | Huelva Isla Saltes Tinto River | 37.181778 N | -6.937831 W | 90 |
| HIO | Huelva Isla Saltes Ocean | 37.180806 N | -6.938719 W | 90 |

**Table S2a:** Geochemical elements included in the anthropogenic enrichment index (AEI), their main anthropogenic sources or processes, rationale for inclusion, and supporting references.

| Element | Main anthropogenic sources / processes | Rationale for inclusion in the index | Supporting references |
|---|---|---|---|
| Lead (Pb) | Mining and smelting, industrial emissions, fossil-fuel combustion, urban runoff, historical leaded gasoline and antifouling activities. | Widely used element of anthropogenic metal enrichment. Pb is efficiently retained in sediments and has been used to reconstruct historical contamination in estuarine sediment cores. | (Adetunji et al., 2022; Alloway, 2013; Berto et al., 2012; O. Bradley et al., 2025; Liang & Mao, 2015; J. M. Pacyna & Pacyna, 2001) |
| Zinc (Zn) | Metallurgy, industrial and urban runoff, wastewater, traffic and infrastructure corrosion. | Common anthropogenic trace-metal contaminant in estuarine and coastal sediments. Sediment cores frequently record historical increases in Zn associated with urbanisation and industrialisation. | (Alloway, 2013; Araújo et al., 2017; Berto et al., 2012; O. Bradley et al., 2025; J. M. Pacyna & Pacyna, 2001) |
| Copper (Cu) | Mining and metallurgy, industrial discharge, urban runoff, agricultural inputs and antifouling activities. | Common component of multi-metal contamination in coastal and estuarine sediments and useful for tracking industrial and urban inputs. | (Adetunji et al., 2022; Alloway, 2013; Berto et al., 2012; O. Bradley et al., 2025; J. M. Pacyna & Pacyna, 2001) |
| Antimony (Sb) | Metallurgy, industrial processes, fossil-fuel combustion and traffic-related emissions. | Particularly useful as a complementary anthropogenic element. Its use as an indicator of human influence has been explicitly evaluated in estuarine sediments, including sites in NW Spain. | (Araújo et al., 2017; Cal-Prieto et al., 2001; J. M. Pacyna & Pacyna, 2001) |
| Cadmium (Cd) | Metal smelting, industrial emissions, phosphate fertilizers, wastewater and urban inputs. | Toxic trace metal frequently enriched above natural background in anthropogenically impacted sediments and commonly used in sediment contamination assessments. | (Adetunji et al., 2022; Alloway, 2013; J. M. Pacyna & Pacyna, 2001; Pan et al., 2010) |
| Mercury (Hg) | Mining, industrial activities, fossil-fuel combustion, wastewater and atmospheric deposition. | Sediment profiles preserve historical changes in Hg deposition and have been extensively used to reconstruct increasing anthropogenic Hg inputs through time. | (O. Bradley et al., 2025; E. G. Pacyna et al., 2010; J. M. Pacyna & Pacyna, 2001) |
| Tin (Sn) | Maritime activities, harbours and historical use of organotin compounds, particularly TBT, in antifouling paints. | Especially relevant in coastal environments. Total Sn and organotin compounds in sediments can record historical maritime contamination and antifouling inputs. Evidence is available specifically from estuarine sediments in northern Spain. | (Berto et al., 2012; O. Bradley et al., 2025; J. M. Pacyna & Pacyna, 2001) |
| Chromium (Cr) | Metallurgy, metal processing, industrial discharge and other anthropogenic sources; also substantial lithogenic contributions. | Frequently included in multi-element assessments of sediment contamination, but interpretation requires consideration of geological background. | (Adetunji et al., 2022; Berto et al., 2012; J. M. Pacyna & Pacyna, 2001) |
| Nickel (Ni) | Metallurgy, fossil-fuel combustion, industrial discharge and urban runoff. | Commonly used in multi-element sediment contamination assessments. | (Begum et al., 2022; J. M. Pacyna & Pacyna, 2001) |
| Arsenic (As) | Mining and smelting, industrial activities, combustion and historical agricultural inputs. | Potentially useful indicator of anthropogenic enrichment. | (Berto et al., 2012; O. Bradley et al., 2025; J. M. Pacyna & Pacyna, 2001) |
| Phosphorus (P) | Fertilizer and agricultural runoff, sewage, wastewater and other nutrient inputs. | Represents nutrient enrichment rather than trace-metal pollution. Sedimentary P has been used to reconstruct historical eutrophication and anthropogenic nutrient loading. | (Dominik et al., 2023; Mahowald et al., 2008; Mekonnen & Hoekstra, 2018) |

**Table S2b:**
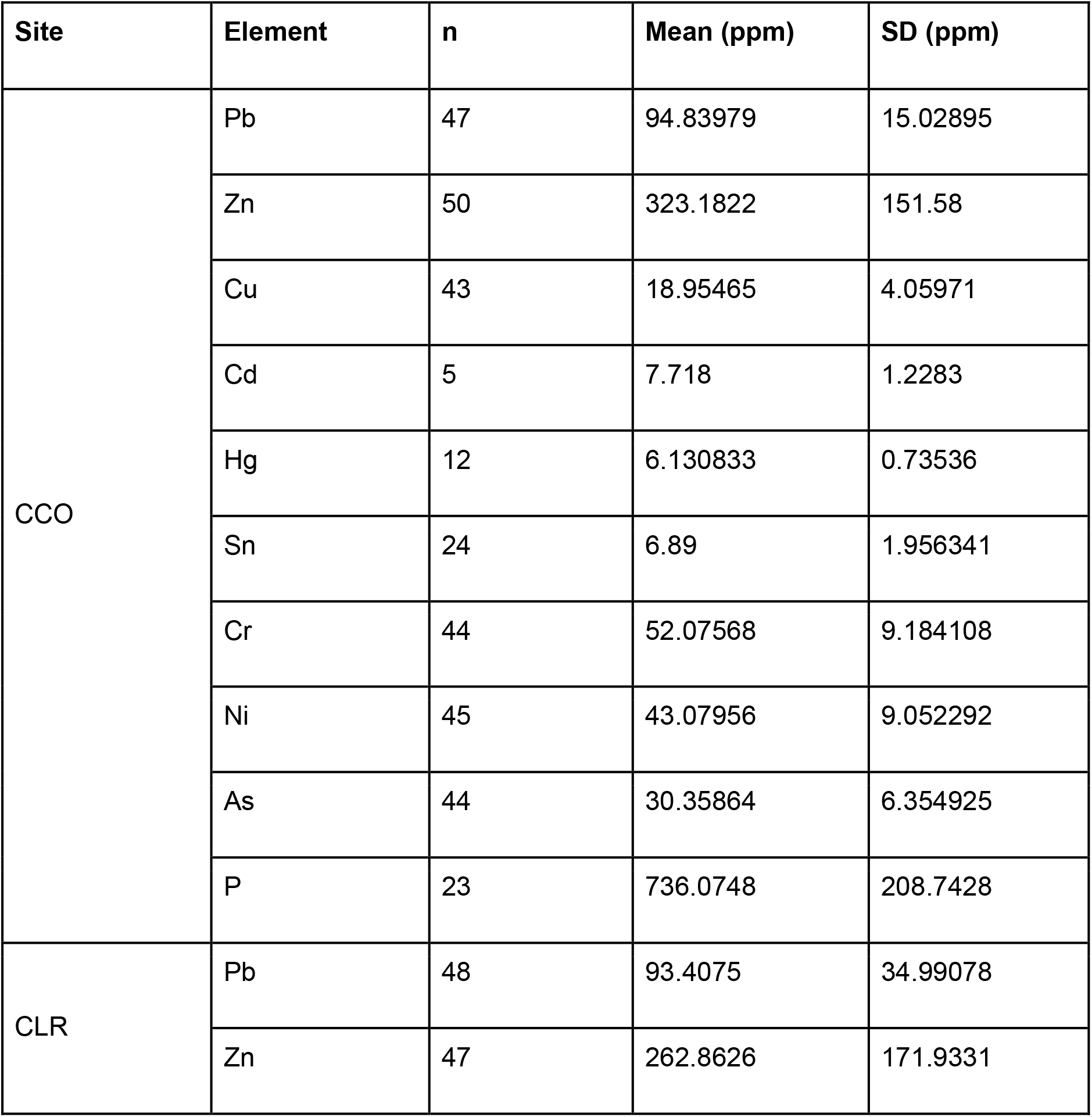
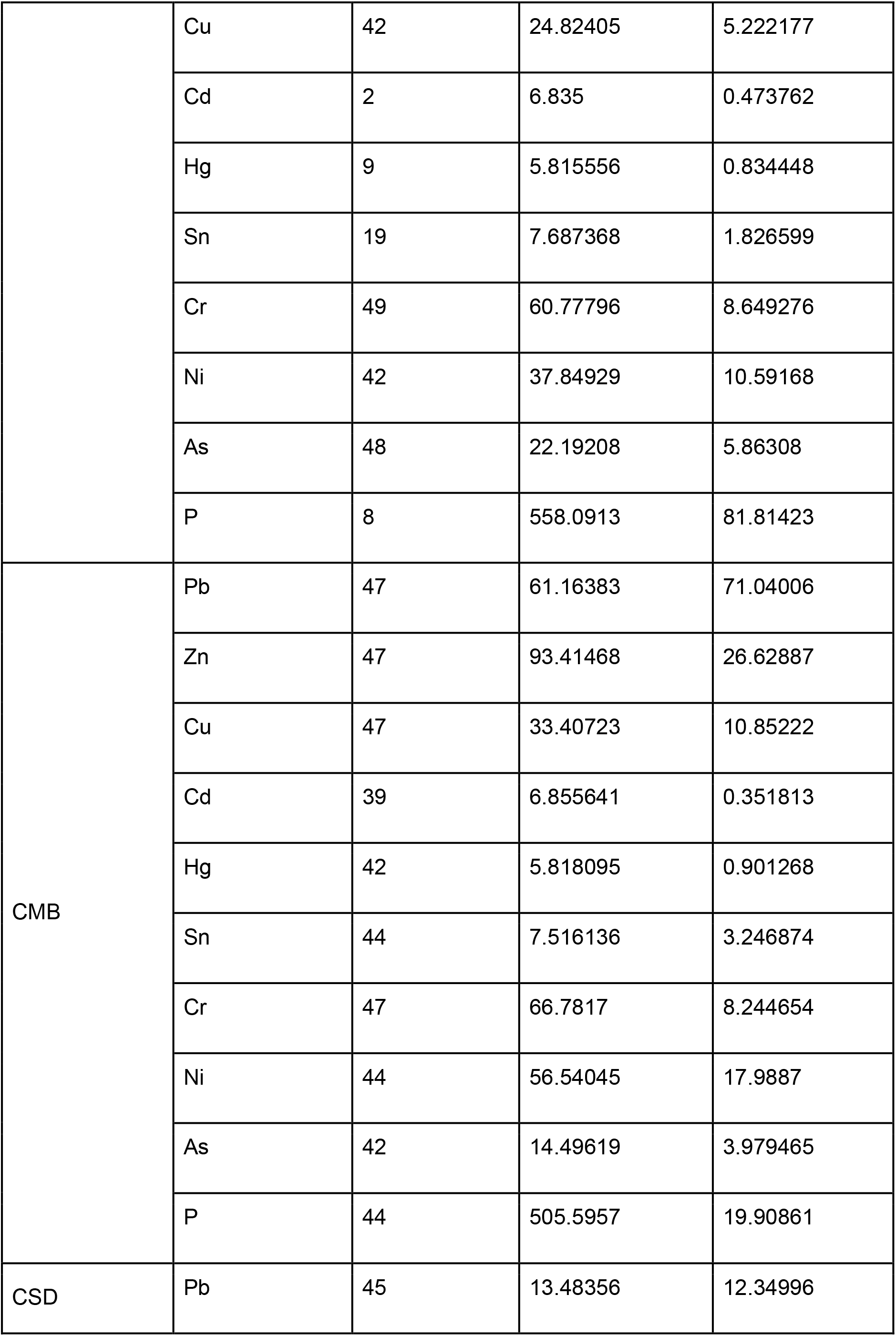
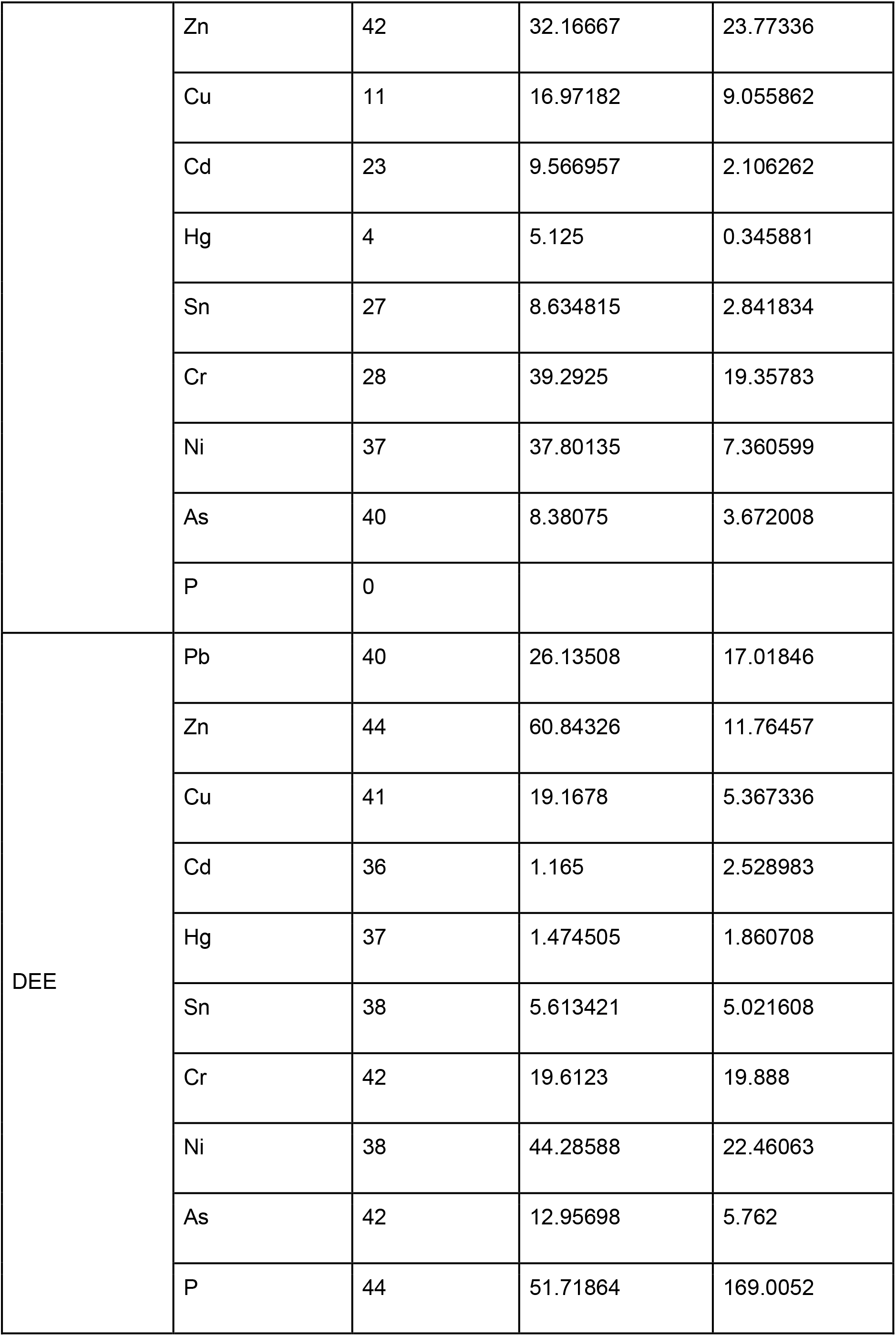
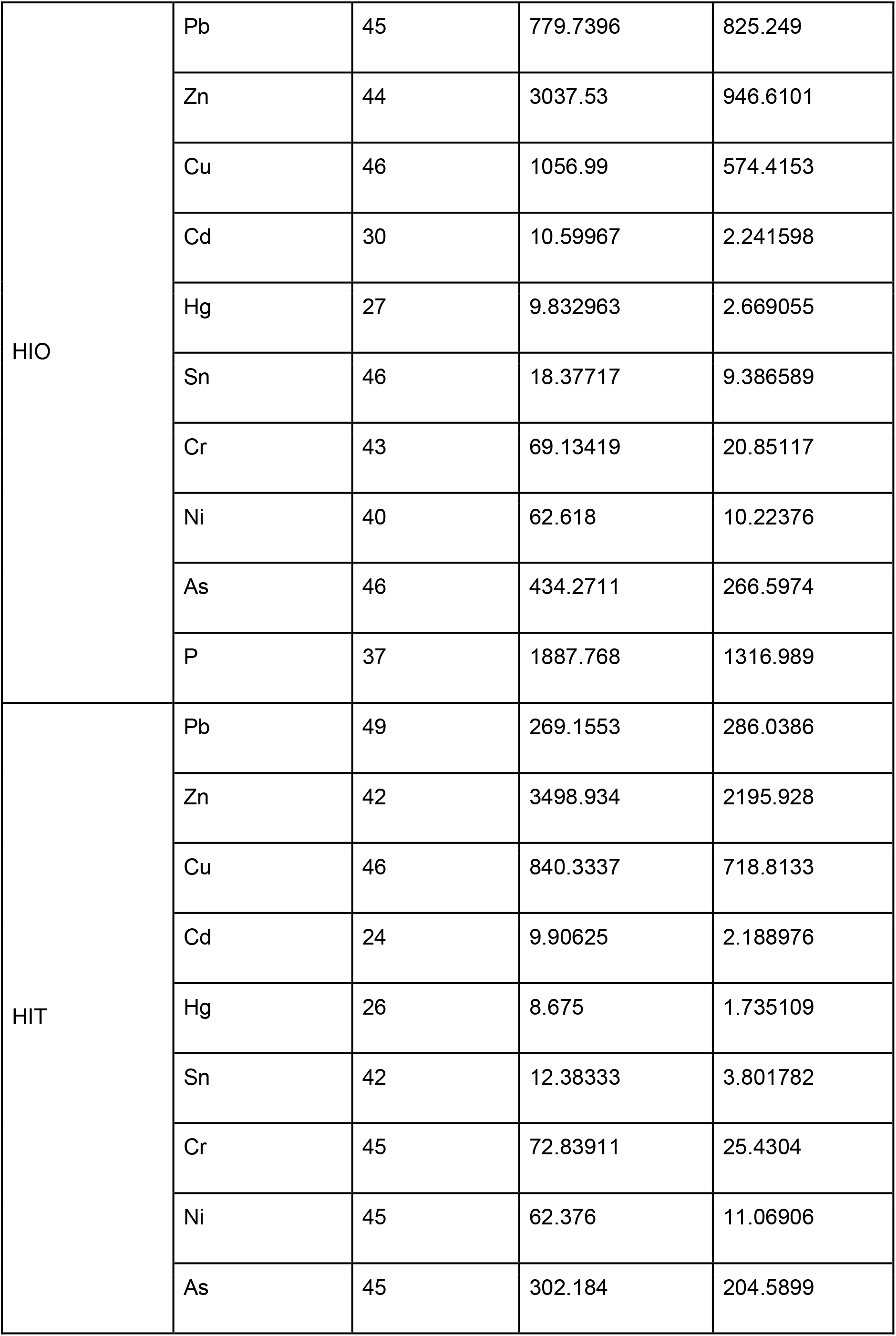
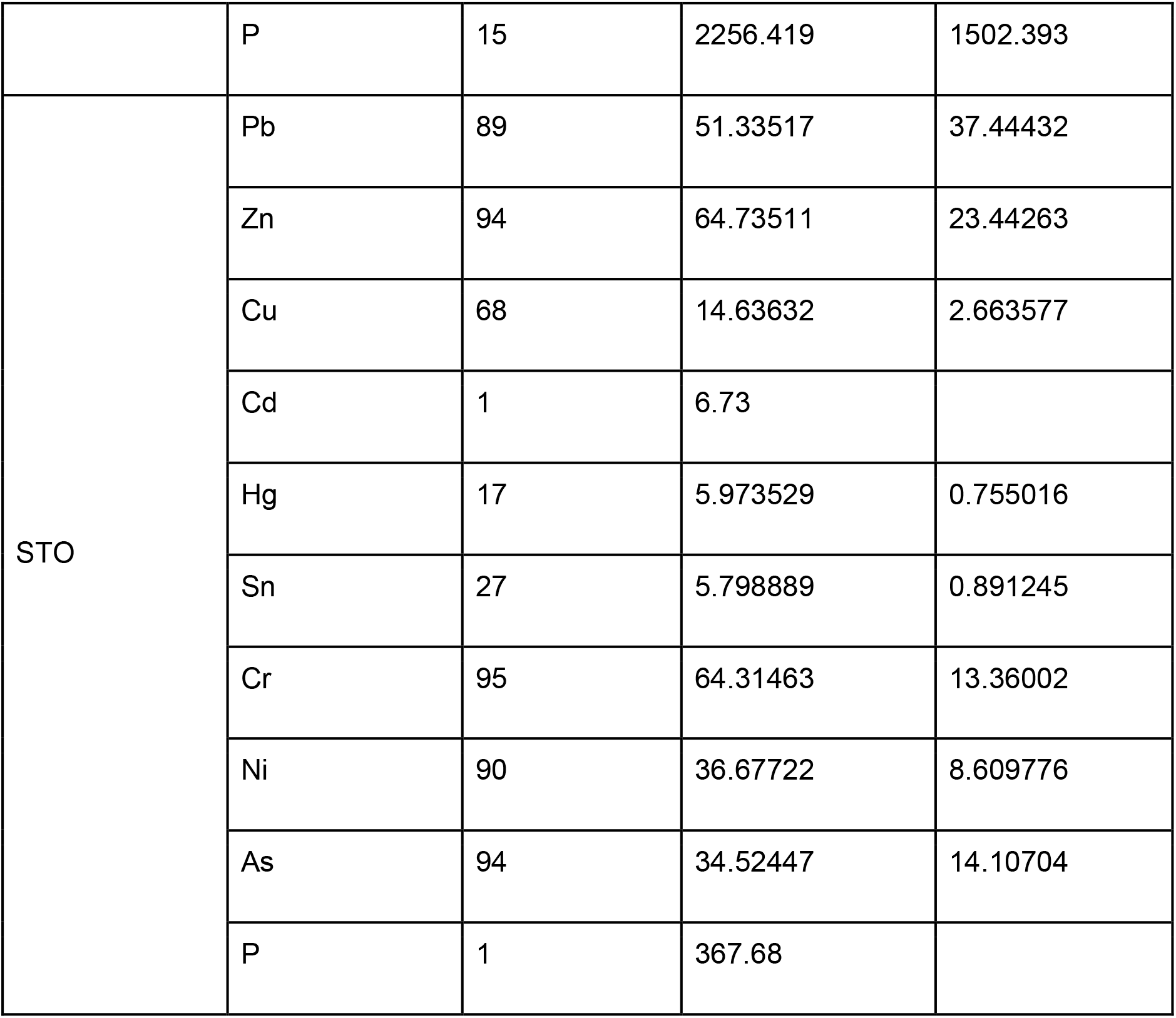
Site-specific descriptive statistics for the elemental measurements retained in the Anthropogenic Enrichment Index. Values represent the mean and standard deviation (SD) of cleaned XRF measurements, expressed in parts per million (ppm), for each element and sediment record. Measurements flagged as potential analytical artefacts or recorded as missing were excluded from the calculations; consequently, the number of valid measurements (n) may differ among elements and records. Sb is not included because measurements were unavailable for some sediment cores, preventing its consistent inclusion in cross-record comparisons.

**Table S3:**
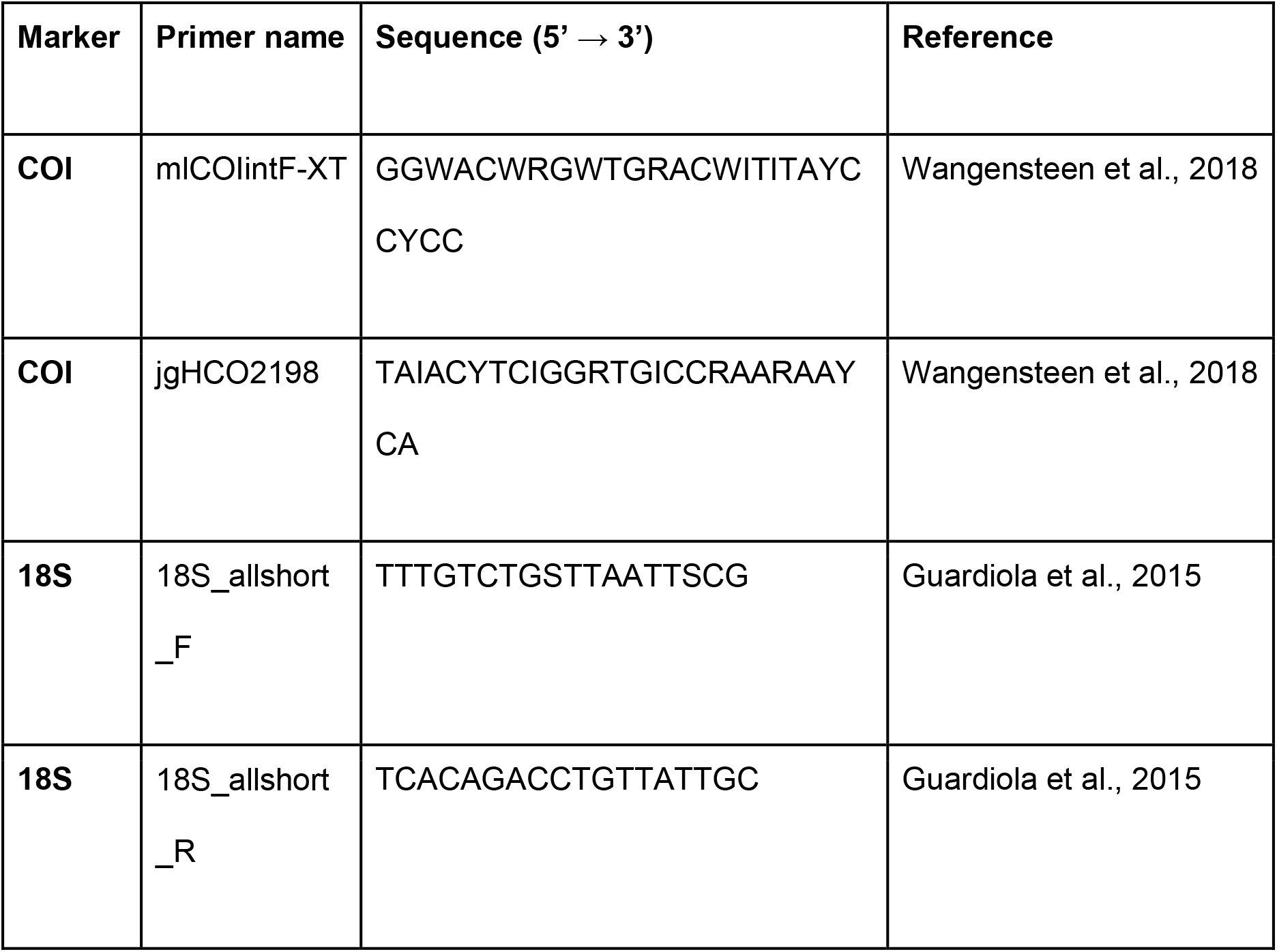
Primer sequences (5′–3′) used to amplify the COI (Leray-XT) and 18S (V7 region) markers in this study. Primers were dual-indexed using mirrored 8-bp tags to minimise tag-jumping artefacts.

**Table S4:** Recent sediment accumulation rates estimated from excess ^210^Pb profiles using the Constant Flux–Constant Sedimentation (CF:CS) model. Sediment accumulation rates and their 95% uncertainty ranges were derived from linear regressions of ln-transformed excess ^210^Pb activity against sediment depth. The unsupported-^210^Pb dating horizon indicates the deepest interval at which excess ^210^Pb remained distinguishable from supported background activity. Basal ages are extrapolations assuming that the recent CF:CS accumulation rate remained constant below the ^210^Pb dating horizon and should not be interpreted as independently dated ages. † Estimates for STO and CSD are poorly constrained because their regressions contained only two unsupported- ^210^Pb observations.

| Core ID | Site | Unsupported<sup>210</sup>Pb dating horizon (cm) | CF:CS SAR (cm yr<sup>-1</sup>) | 95% range (cm yr<sup>−1</sup>) | Core base (cm) | Extrapolated basal age assuming constant CF:CS SAR (CE) |
|---|---|---|---|---|---|---|
| DEE | Ebro Delta | 15 | 0.11 | 0.09 - 0.15 | 70 | ~1384 |
| STO | Santoña | 10 | 0.1 | 0.1† | 89 | ~1134 |
| CCO | Cicero | 48 | 0.53 | 0.42 - 0.73 | 49 | ~1939 |
| COL | Colindres | 49 | 0.63 | 0.50 - 0.87 | 49 | ~1953 |
| CSD | Cadiz Salina Dolores | 3 | 0.1 | 0.1† | 89 | ~1135 |
| CMB | Cadiz Militar Base | 15.5 | 0.15 | 0.08 - 0.58 | 89 | ~1419 |
| HIT | Huelva Tinto River | 40 | 0.42 | 0.3019 - 0.6850 | 89 | ~1812 |
| HIO | Huelva Saltes Island Ocean | 65 | 0.63 | 0.417 - 1.302 | 89 | ~1884 |

**Table S5:** AMS radiocarbon determinations and calibrated probability ranges for intertidal sediment samples. Conventional radiocarbon ages were calibrated using the IntCal20 Northern Hemisphere atmospheric calibration curve. Reported ranges encompass 95.4% of the calibrated probability distribution; the probability associated with each discontinuous interval is shown in parentheses. No marine-reservoir or old-carbon correction was applied.

| **Core ID** | **Depth (cm)** | **Conventional<sup>14</sup>C age (yr BP)** | **1σ error (yr)** | **Calibration curve** | **Calibrated age, 95.4% HPD (cal BCE/CE)** |
|---|---|---|---|---|---|
| DEE | 70 | 1570 | 30 | IntCal20 | 424–564 CE (95.6%) |
| CSD | 85 | 2735 | 35 | IntCal20 | 975–954 BCE (4.0%);<br>933–809 BCE (91.5%) |
| CMB | 85 | 580 | 30 | IntCal20 | 1305–1364 CE (64.3%);<br>1383–1419 CE (31.2%) |
| HIT | 85 | 955 | 30 | IntCal20 | 1027–1159 CE (95.5%) |
| HIO | 85 | 1095 | 30 | IntCal20 | 889–995 CE (90.2%);<br>1005–1018 CE (5.4%) |

**Table S6:**
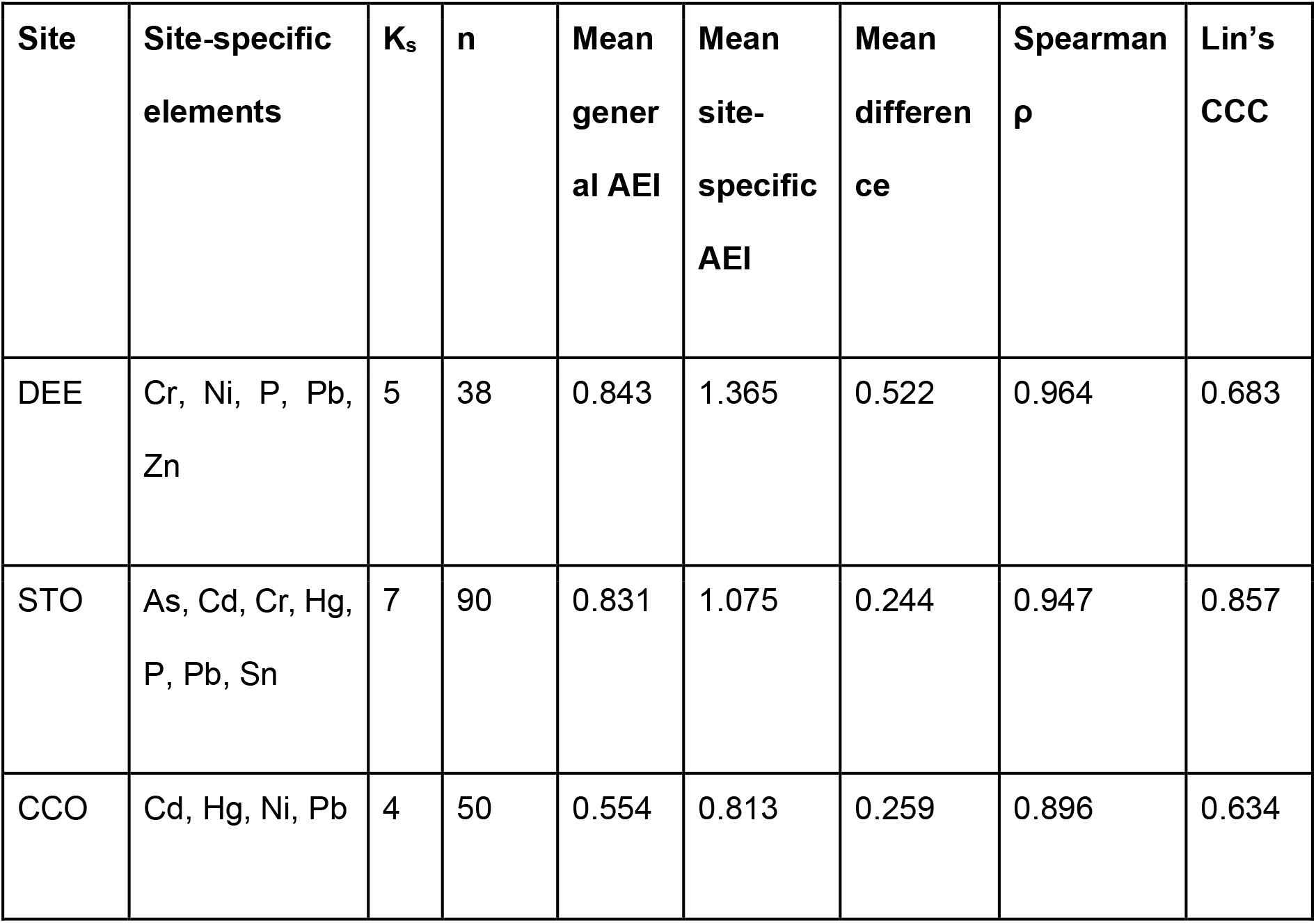
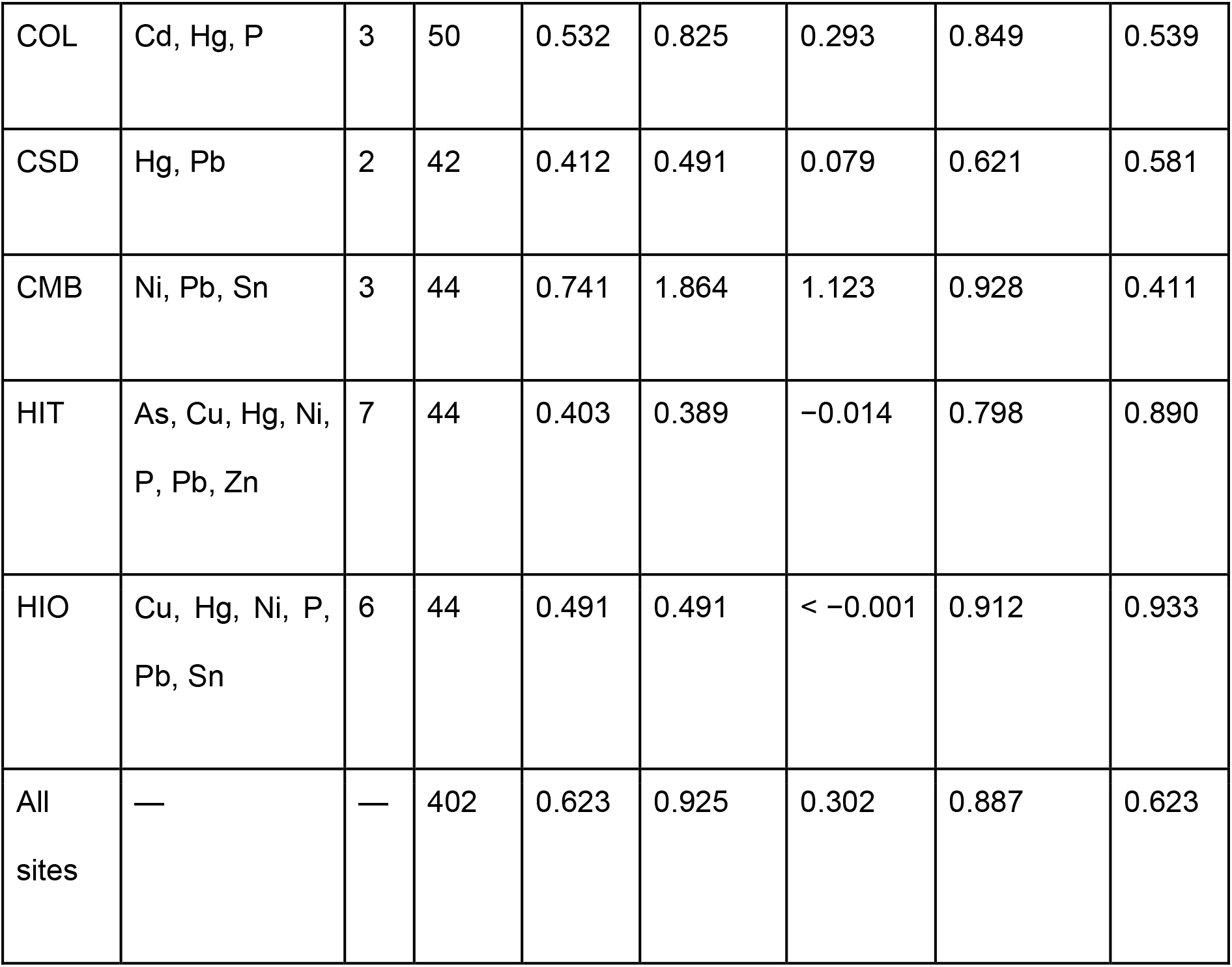
Composition of the site-specific Anthropogenic Enrichment Index (AEI) and statistical comparison with the general AEI. The general AEI was calculated as the equal-weight arithmetic mean of positive element-specific robust z-scores across the ten candidate elements with available XRF measurements. The site-specific AEI was calculated using the same aggregation procedure but was restricted to elements showing strong enrichment (z^robust^ > 3) in at least two samples from the corresponding sediment record. Ks indicates the number of elements included in each site-specific AEI, and n is the number of paired observations used in the comparison. Mean difference was calculated as site-specific AEI minus general AEI; positive values therefore indicate higher site-specific estimates. Spearman’s rank correlation coefficient (ρ) measures similarity in the ordering of samples, whereas Lin’s concordance correlation coefficient (CCC) evaluates absolute agreement with the line of identity. The pooled “All sites” row summarises the comparison across all paired observations and should be interpreted alongside the site-level results because observations are grouped within sediment records.

**Table S7:**
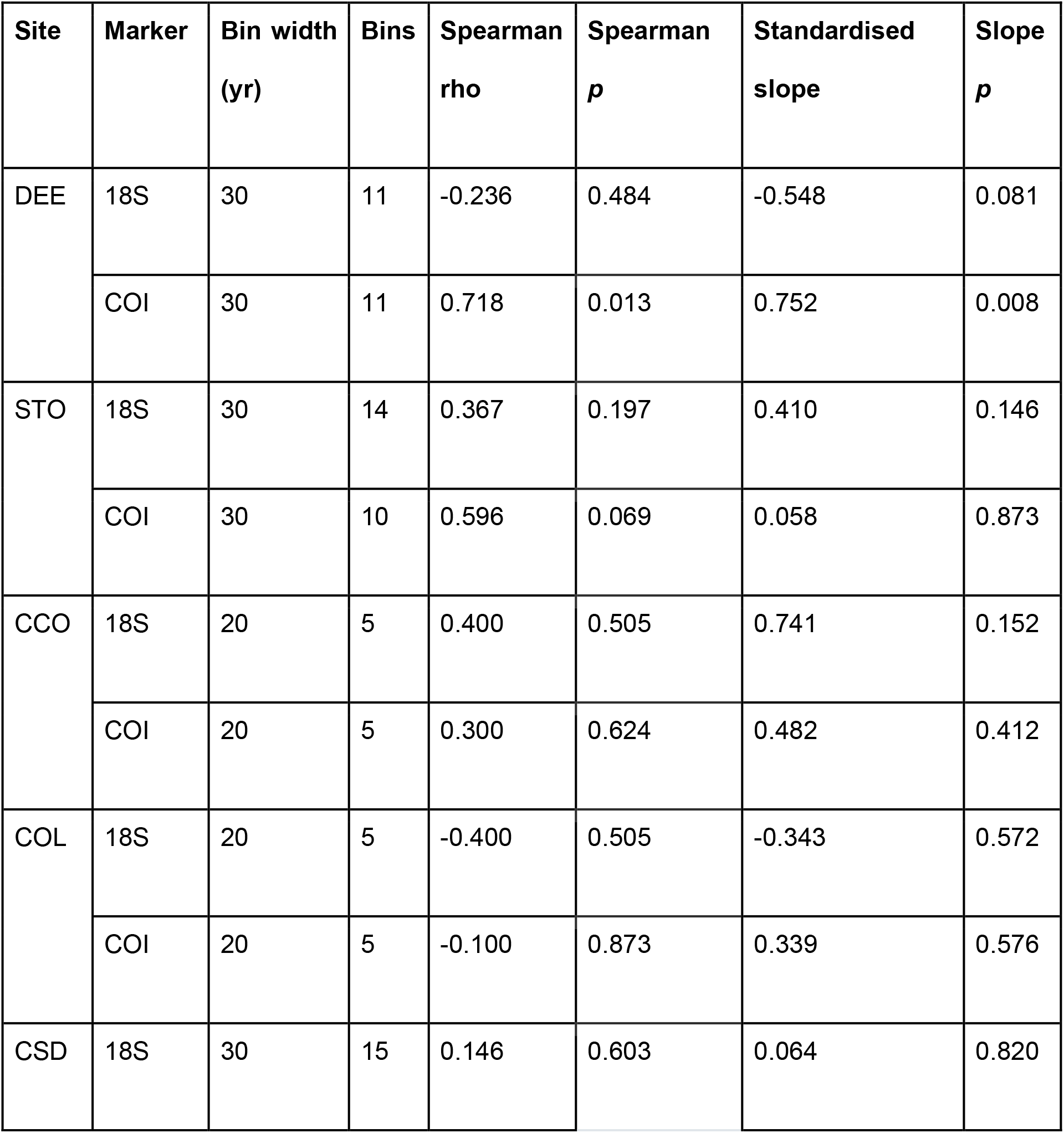
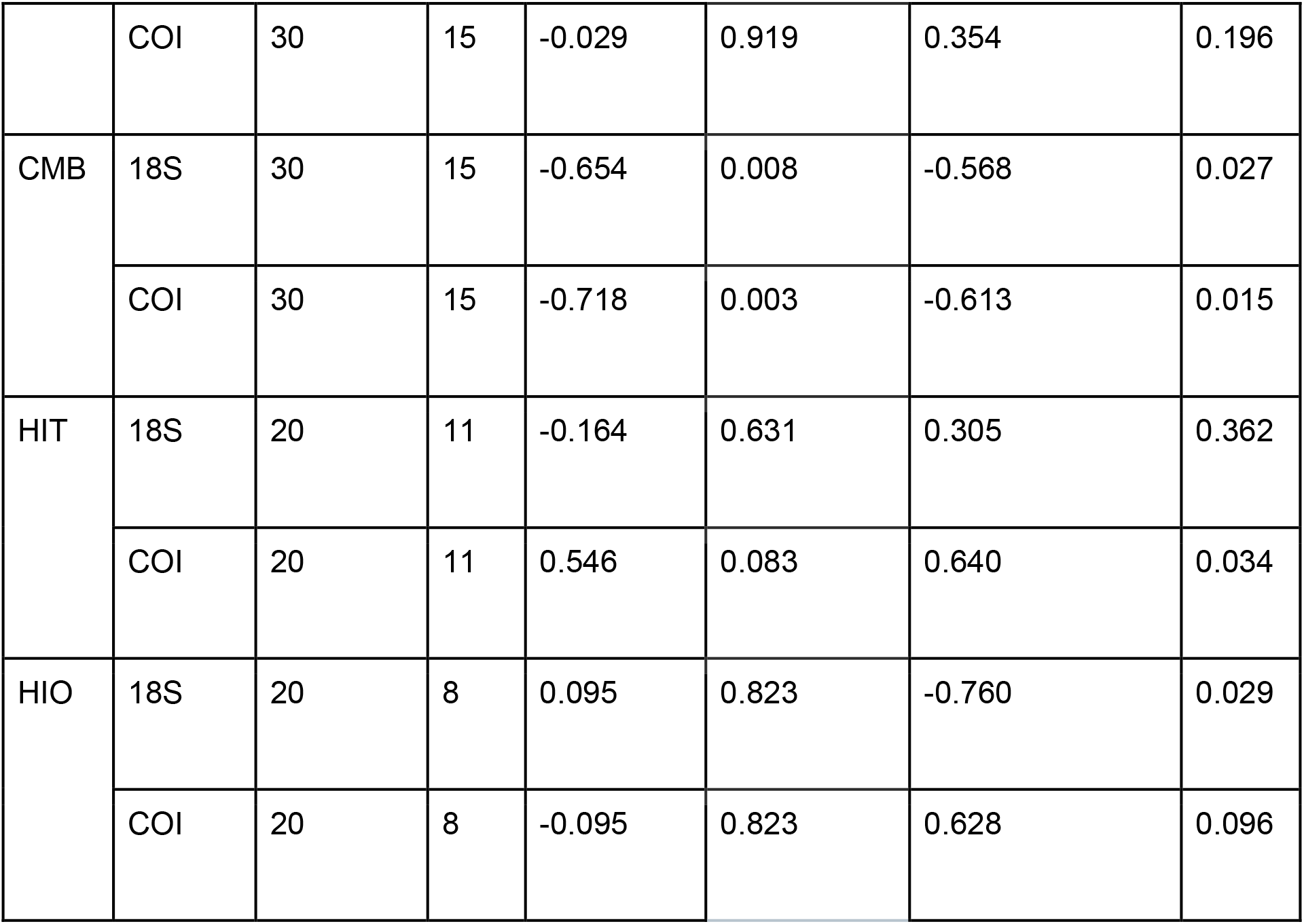
Site-specific associations between eukaryotic ASV richness and the general Anthropogenic Enrichment Index (AEI). Analyses were based on median richness and AEI values within 20-year bins for CCO, COL, HIT and HIO and 30-year bins for DEE, STO, CSD and CMB. Spearman’s rank correlation coefficient describes monotonic association. Standardised slopes were obtained from linear models relating within-series standardised richness to standardised AEI. Nominal p-values are reported for the 16 site-by-marker comparisons. COI richness was analysed in its original units.

| **Site** | **Marker** | **Bin width (yr)** | **Bins** | **Spearman rho** | **Spearman *p*** | **Standardised slope** | **Slope *p*** |
|---|---|---|---|---|---|---|---|
| DEE | 18S | 30 | 11 | -0.236 | 0.484 | -0.548 | 0.081 |
|  | COI | 30 | 11 | 0.718 | 0.013 | 0.752 | 0.008 |
| STO | 18S | 30 | 14 | 0.367 | 0.197 | 0.410 | 0.146 |
|  | COI | 30 | 10 | 0.596 | 0.069 | 0.058 | 0.873 |
| CCO | 18S | 20 | 5 | 0.400 | 0.505 | 0.741 | 0.152 |
|  | COI | 20 | 5 | 0.300 | 0.624 | 0.482 | 0.412 |
| COL | 18S | 20 | 5 | -0.400 | 0.505 | -0.343 | 0.572 |
|  | COI | 20 | 5 | -0.100 | 0.873 | 0.339 | 0.576 |
| CSD | 18S | 30 | 15 | 0.146 | 0.603 | 0.064 | 0.820 |
|  | COI | 30 | 15 | -0.029 | 0.919 | 0.354 | 0.196 |
| CMB | 18S | 30 | 15 | -0.654 | 0.008 | -0.568 | 0.027 |
|  | COI | 30 | 15 | -0.718 | 0.003 | -0.613 | 0.015 |
| HIT | 18S | 20 | 11 | -0.164 | 0.631 | 0.305 | 0.362 |
|  | COI | 20 | 11 | 0.546 | 0.083 | 0.640 | 0.034 |
| HIO | 18S | 20 | 8 | 0.095 | 0.823 | -0.760 | 0.029 |
|  | COI | 20 | 8 | -0.095 | 0.823 | 0.628 | 0.096 |

**Table S8:** Time-adjusted and autocorrelation-sensitive models of the relationship between eukaryotic ASV richness and AEI. Both richness and AEI were standardised within each site and marker. Time-adjusted models included standardised calendar year as a covariate. CAR(1) sensitivity models used the same fixed effects and a continuous first-order autoregressive residual structure. CAR(1) models were fitted only for records containing at least eight paired temporal bins; blank entries indicate insufficient temporal replication. The time-adjusted and CAR(1) models produced identical coefficient estimates and nominal p-values; therefore, a single p-value is shown. Models were not fitted for CCO and COL because only five paired temporal bins were available, providing insufficient residual degrees of freedom for models containing both AEI and calendar year and for estimation of a CAR(1) residual structure.

| **Site** | **Marker** | **n bins** | **Adjusted slope** | **CAR(1) slope** | ***p*** |
|---|---|---|---|---|---|
| DEE | 18S | 11 | -0.268 | -0.268 | 0.635 |
|  | COI | 11 | 1.093 | 1.093 | 0.030 |
| STO | 18S | 14 | 0.354 | 0.354 | 0.188 |
|  | COI | 10 | 0.096 | 0.096 | 0.771 |
| CCO | 18S | 5 | — | — | — |
|  | COI | 5 | — | — | — |
| COL | 18S | 5 | — | — | — |
|  | COI | 5 | — | — | — |
| CSD | 18S | 15 | 0.092 | 0.092 | 0.776 |
|  | COI | 15 | 0.131 | 0.131 | 0.614 |
| CMB | 18S | 15 | -0.308 | -0.308 | 0.096 |
|  | COI | 15 | -0.395 | -0.395 | 0.056 |
| HIT | 18S | 11 | 0.240 | 0.240 | 0.507 |
|  | COI | 11 | 0.514 | 0.514 | 0.068 |
| HIO | 18S | 8 | -0.622 | -0.622 | 0.058 |
|  | COI | 8 | 0.371 | 0.371 | 0.076 |

**Table S9:** Associations between eukaryotic ASV richness and element-specific Pb and Hg enrichment across sediment records. Pb and Hg enrichment was expressed using element-specific robust z-scores and summarised as medians within the same site-specific temporal bins used for the richness–AEI analyses (20 years for CCO, COL, HIT and HIO; 30 years for DEE, STO, CSD and CMB). E indicates the element, n indicates the number of paired bins. Spearman’s ρ measures rank association, whereas β represents the standardised richness response to a one-standard-deviation increase in the corresponding element score. Year-adjusted models included standardised calendar year as an additional predictor to separate potentially shared linear temporal trends. Parentheses contain the nominal p-value followed by the Benjamini–Hochberg-adjusted q-value. Nominal p-values below 0.05 are shown in bold; no association remained significant after multiple-testing correction. † indicates an element not retained in the corresponding site-specific AEI but included here for completeness. Em dashes indicate models that were not fitted because fewer than six paired bins were available.

| Site | Marker | E | n | Spearman rho (p; q) | Unadjusted beta (p; q) | Year-adjusted beta (p; q) |
|---|---|---|---|---|---|---|
| DEE | 18S | Pb | 11 | -0.091 (0.790; 0.872) | -0.303 ( 0.366; 0.650) | 0.013 (0.971; 0.974) |
|  |  | Hg<sup>†</sup> | 11 | -0.618 (**0.043**; 0.319) | -0.486 ( 0.130; 0.416) | -0.354 (0.241; 0.453) |
|  | COI | Pb | 11 | 0.336 (0.312; 0.629) | 0.318 (0.340; 0.650) | 0.039 (0.918; 0.974) |
|  |  | Hg<sup>†</sup> | 11 | -0.327 (0.326; 0.629) | -0.288 (0.391; 0.659) | -0.478 (0.110; 0.453) |
| STO | 18S | Pb | 14 | -0.090 (0.759; 0.872) | 0.038 (0.896; 0.900) | -0.520 (0.159; 0.453) |
|  |  | Hg | 14 | 0.279 (0.334; 0.629) | 0.091 (0.757; 0.849) | 0.346 (0.245; 0.453) |
|  | COI | Pb | 10 | 0.304 (0.393; 0.629) | 0.119 (0.744; 0.849) | -0.567 (0.205; 0.453) |
|  |  | Hg | 10 | 0.012 (0.973; 0.973) | -0.226 (0.530; 0.771) | 0.116 (0.771; 0.974) |
| CCO | 18S | Pb | 5 | -0.200 (0.747; 0.872) | -0.227 (0.714; 0.849) | — |
|  |  | Hg | 5 | -0.500 (0.391; 0.629) | -0.116 (0.852; 0.900) | — |
|  | COI | Pb | 5 | -0.500 (0.391; 0.629) | -0.314 (0.607; 0.809) | — |
|  |  | Hg | 5 | -0.600 (0.285; 0.629) | -0.526 (0.362; 0.650) | — |
| COL | 18S | Pb<sup>†</sup> | 5 | -0.900 (**0.037**; 0.319) | -0.847 (0.070; 0.330) | — |
|  |  | Hg | 5 | -0.800 (0.104; 0.476) | -0.527 (0.361; 0.650) | — |
|  | COI | Pb<sup>†</sup> | 5 | -0.100 (0.873; 0.931) | -0.650 (0.235; 0.626) | — |
|  |  | Hg | 5 | 0.500 (0.391; 0.629) | 0.381 (0.527; 0.771) | — |
| CSD | 18S | Pb | 15 | -0.032 (0.909; 0.939) | -0.131 (0.641; 0.820) | -0.147 (0.652; 0.956) |
|  |  | Hg | 15 | 0.111 (0.694; 0.872) | 0.269 (0.333; 0.650) | 0.288 (0.344; 0.550) |
|  | COI | Pb | 15 | 0.241 (0.386; 0.629) | 0.463 (0.082; 0.330) | 0.252 (0.330; 0.550) |
|  |  | Hg | 15 | -0.150 (0.593; 0.863) | -0.272 (0.326; 0.650) | -0.095 (0.704; 0.956) |
| CMB | 18S | Pb | 15 | -0.514 (**0.050**; 0.319) | -0.663 (**0.007**; 0.113) | -0.307 (0.149; 0.453) |
|  |  | Hg<sup>†</sup> | 15 | -0.521 (**0.046**; 0.319) | -0.540 (**0.038**; 0.302) | -0.428 (**0.006**; 0.142) |
|  | COI | Pb | 15 | -0.689 (**0.004**; 0.143) | -0.694 (**0.004**; 0.113) | -0.421 (0.076; 0.453) |
|  |  | Hg<sup>†</sup> | 15 | -0.225 (0.420; 0.640) | -0.184 (0.512; 0.771) | -0.075 (0.717; 0.956) |
| HIT | 18S | Pb | 11 | 0.373 (0.259; 0.629) | 0.555 (0.077; 0.330) | 1.787 (**0.012**; 0.142) |
|  |  | Hg | 11 | -0.127 (0.709; 0.872) | -0.043 (0.900; 0.900) | 0.030 (0.933; 0.974) |
|  | COI | Pb | 11 | 0.364 (0.272; 0.629) | 0.516 (0.104; 0.370) | -0.050 (0.947; 0.974) |
|  |  | Hg | 11 | 0.473 (0.142; 0.568) | 0.185 (0.587; 0.809) | 0.348 (0.237; 0.453) |
| HIO | 18S | Pb | 8 | 0.119 (0.779; 0.872) | 0.494 (0.213; 0.619) | 0.636 (**0.034**; 0.275) |
|  |  | Hg | 8 | -0.643 (0.086; 0.456) | -0.764 (**0.027**; 0.290) | -0.715 (0.173; 0.453) |
|  | COI | Pb | 8 | 0.119 (0.779; 0.872) | -0.124 (0.770; 0.849) | -0.302 (0.152; 0.453) |
|  |  | Hg | 8 | 0.452 (0.260; 0.629) | 0.674 (0.067; 0.330) | 0.012 (0.974; 0.974) |

**Table S10:**
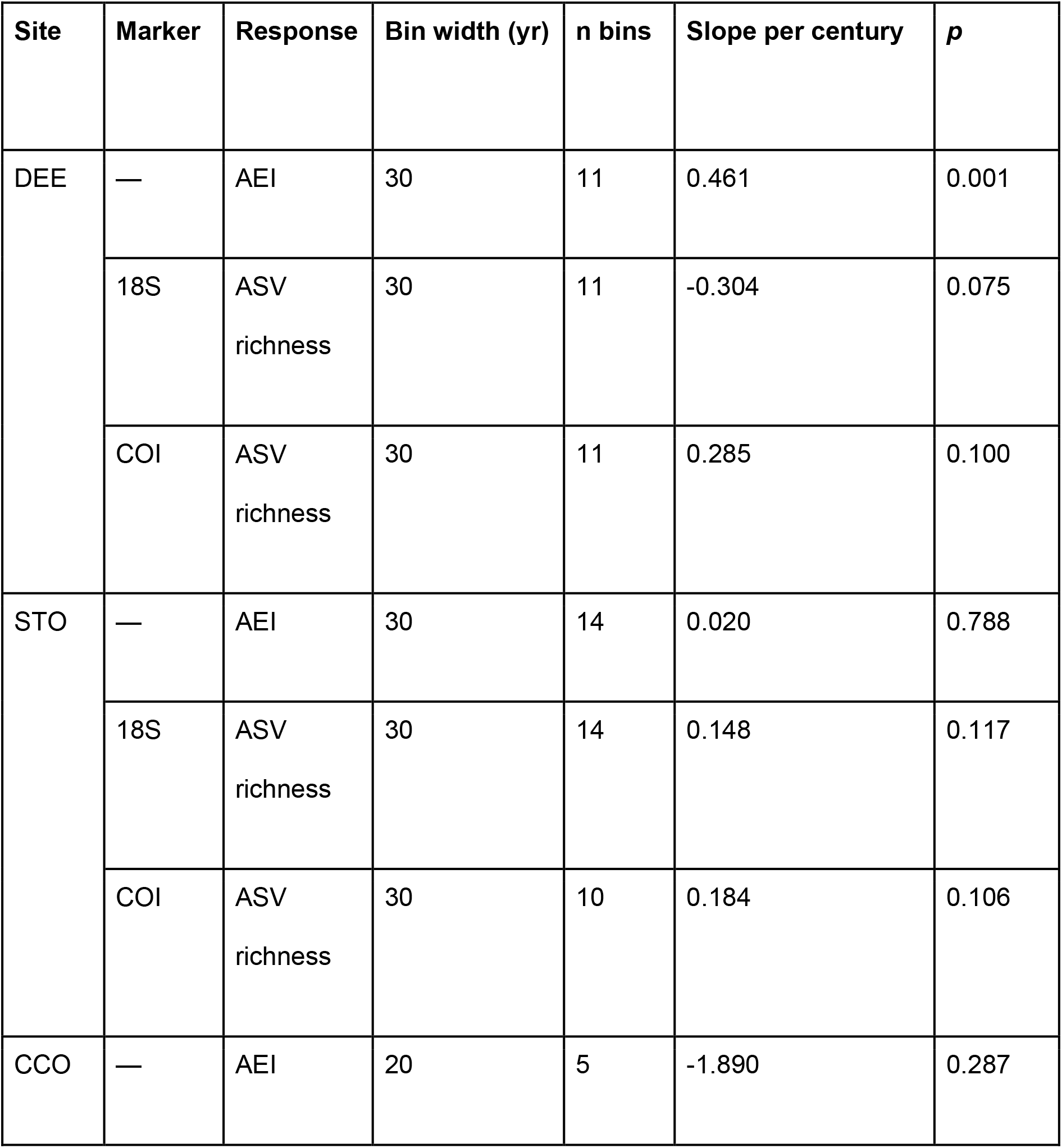
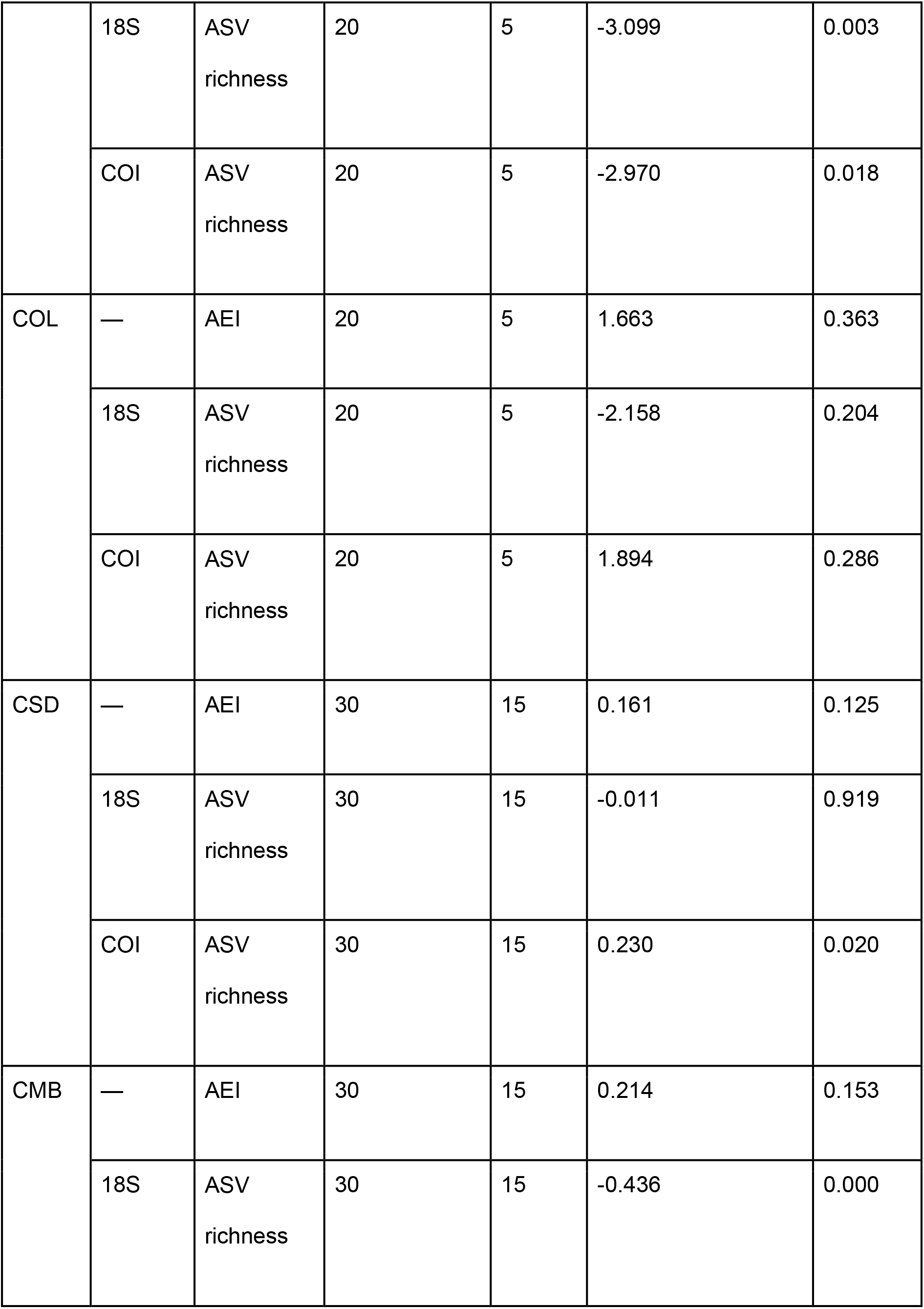
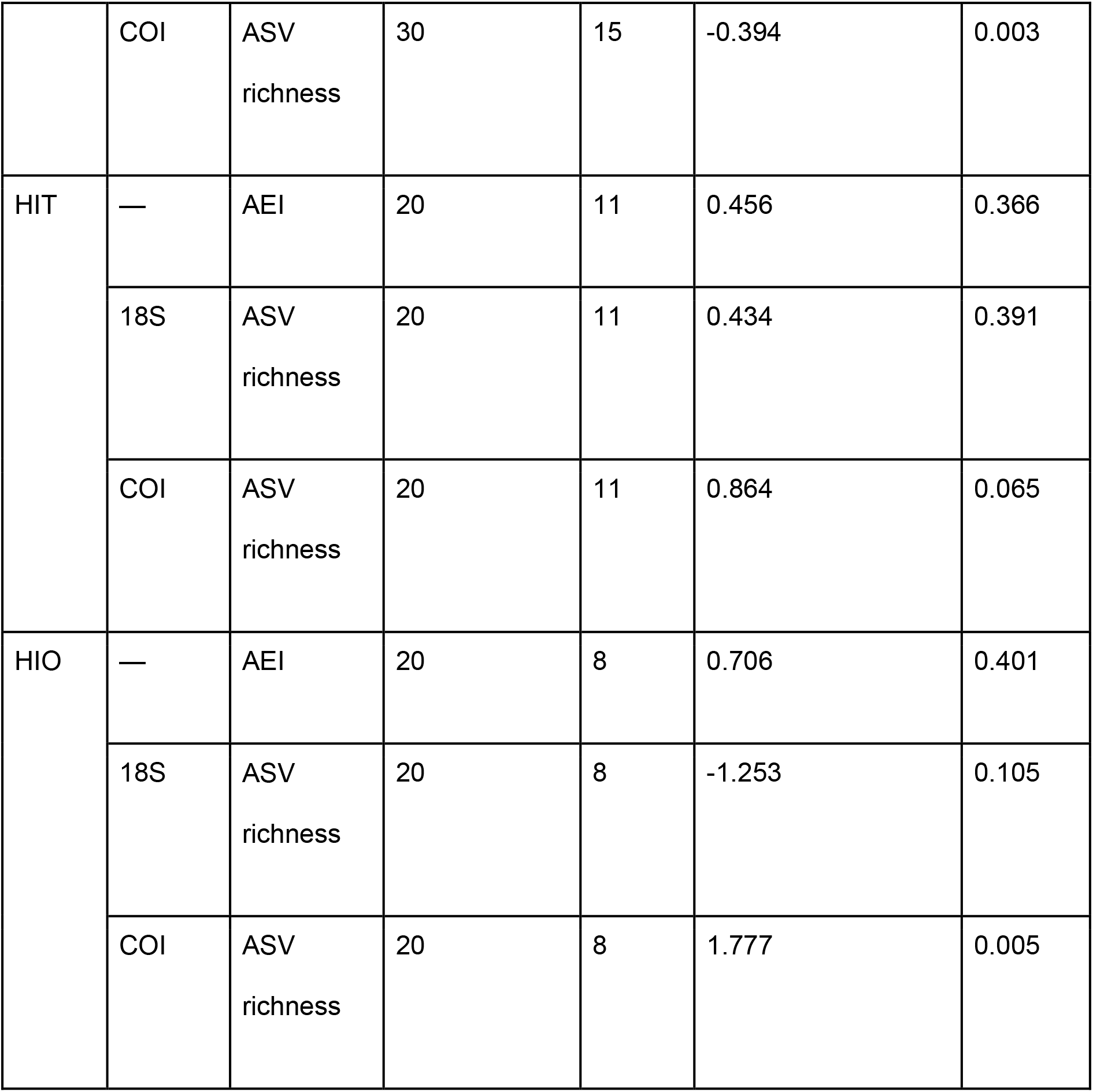
Overall temporal trends in eukaryotic ASV richness and the general AEI. Linear slopes were estimated from standardised median-binned values and are expressed as within-series standard deviations per century. Richness models were fitted separately for each site and metabarcoding marker, whereas AEI models were fitted once per site. These coefficients summarise the average linear tendency across each complete record and do not imply monotonic change within shorter intervals.

**Table S11:** Direction of consecutive changes in eukaryotic ASV richness and AEI. Values show the number and percentage of transitions in which richness and AEI both decreased, both increased, or changed in opposite directions. Opposite-direction transitions comprise richness decreases accompanied by AEI increases and richness increases accompanied by AEI decreases. These summaries are descriptive because consecutive transitions are temporally dependent and the shortest records contain only four transitions.

| **Site** | **Marker** | **Total** | **Both ↓ %** | **Both ↑ %** | **Richness ↓ / AEI ↑ %** | **Richness ↑ / AEI ↓%** | **Opposite %** |
|---|---|---|---|---|---|---|---|
| DEE | 18S | 10 | 30.0 | 20.0 | 20.0 | 30.0 | 50.0 |
|  | COI | 10 | 40.0 | 40.0 | 0.0 | 20.0 | 20.0 |
| STO | 18S | 13 | 38.5 | 30.8 | 7.7 | 23.1 | 30.8 |
|  | COI | 8 | 25.0 | 25.0 | 0.0 | 50.0 | 50.0 |
| CCO | 18S | 4 | 75.0 | 0.0 | 25.0 | 0.0 | 25.0 |
|  | COI | 4 | 50.0 | 0.0 | 25.0 | 25.0 | 50.0 |
| COL | 18S | 4 | 25.0 | 0.0 | 50.0 | 25.0 | 75.0 |
|  | COI | 4 | 25.0 | 25.0 | 25.0 | 25.0 | 50.0 |
| CSD | 18S | 14 | 21.4 | 28.6 | 28.6 | 21.4 | 50.0 |
|  | COI | 14 | 14.3 | 35.7 | 21.4 | 28.6 | 50.0 |
| CMB | 18S | 14 | 35.7 | 21.4 | 21.4 | 21.4 | 42.9 |
|  | COI | 14 | 28.6 | 14.3 | 28.6 | 28.6 | 57.1 |
| HIT | 18S | 10 | 20.0 | 20.0 | 40.0 | 20.0 | 60.0 |
|  | COI | 10 | 20.0 | 40.0 | 20.0 | 20.0 | 40.0 |
| HIO | 18S | 7 | 57.1 | 14.3 | 14.3 | 14.3 | 28.6 |
|  | COI | 7 | 42.9 | 14.3 | 14.3 | 28.6 | 42.9 |

**Table S12:**
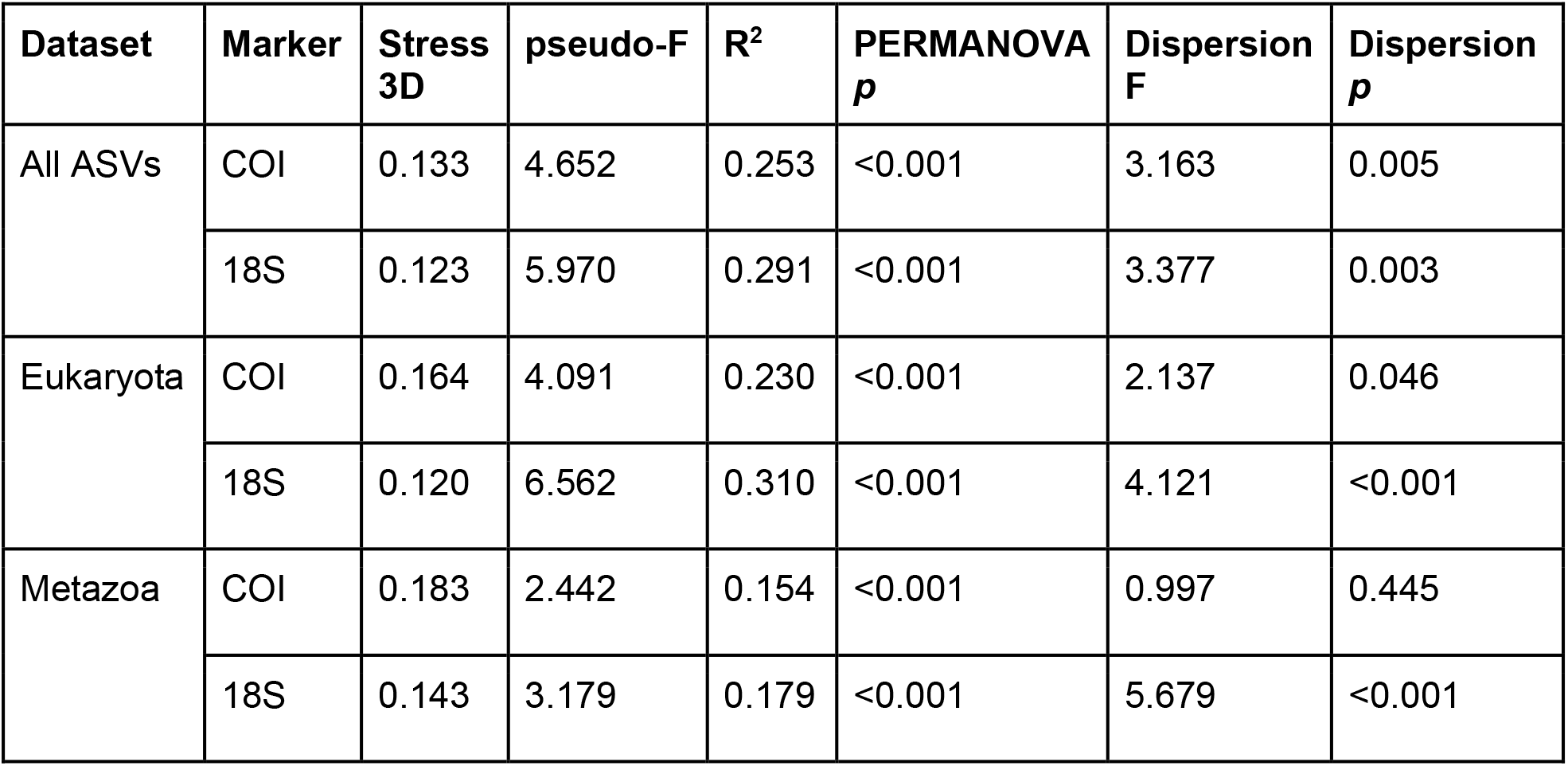
Non-metric multidimensional scaling performance, PERMANOVA results and homogeneity of multivariate dispersion for community-composition datasets. Analyses were conducted separately for the COI and 18S markers using three taxonomic datasets: all retained ASVs irrespective of taxonomic assignment, ASVs assigned to Eukaryota, and ASVs assigned to Metazoa. n indicates the number of sediment samples included after data filtering. Stress values correspond to the complete three-dimensional nMDS solutions based on Bray– Curtis dissimilarities of sample-wise relative ASV abundances. Differences in community composition among sediment records were evaluated using PERMANOVA with Site as the explanatory factor and 9,999 permutations. The pseudo-F statistic describes the ratio of among-record to within-record variation, and R^2^ represents the proportion of compositional variation associated with Site. Homogeneity of multivariate dispersion was evaluated using distances to the spatial median and a permutation test with 9,999 permutations. Significant dispersion results indicate that PERMANOVA outcomes may reflect differences in within-record variability as well as differences in multivariate centroid location.

**Table S13:** Intervals of concurrent multiproxy change identified within each sediment record using moving-window regressions. Start and end years delimit consecutive focal years during which at least three of the five proxy groups (eukaryotic ASV richness, δ¹³C, δ¹⁵N, C:N ratio and the general Anthropogenic Enrichment Index (AEI)) showed an absolute estimated change of at least 0.25 within-series standard deviations over the corresponding temporal window. Maximum active proxies indicates the greatest number of proxy groups simultaneously meeting this criterion within each interval. The representative directional signature corresponds to the year with maximum proxy concurrence; when several years were tied, the year closest to the interval midpoint was selected. Potential intensification denotes increasing AEI accompanied by decreasing richness and marked change in at least one additional proxy, whereas potential relaxation denotes decreasing AEI accompanied by increasing richness and marked change in at least one additional proxy. Remaining configurations were classified as other concurrent change. Moving windows spanned 25 years for CCO, COL, HIT and HIO and 100 years for DEE, STO, CSD and CMB. These classifications are exploratory and do not demonstrate causal relationships or simultaneous statistical significance among proxies.

| **Site** | **Change class** | **Start year (CE)** | **End year (CE)** | **Max. active proxies** | **Representative year (CE)** | **Representative directional signature** |
|---|---|---|---|---|---|---|
| DEE | Other | 1361 | 1396 | 3 | 1378 | ASV Richness – \| δ<sup>13</sup>C – \| δ<sup>15</sup>N ↓ \| C:N ↑ \| AEI ↓ |
|  | Other | 1416 | 1424 | 3 | 1420 | ASV Richness – \| δ<sup>13</sup>C – \| δ<sup>15</sup>N ↓ \| C:N ↓ \| AEI ↑ |
|  | Other | 1635 | 1652 | 3 | 1643 | ASV Richness – \| δ<sup>13</sup>C ↓ \| δ<sup>15</sup>N – \| C:N ↓ \| AEI ↓ |
|  | Other | 1663 | 1680 | 3 | 1671 | ASV Richness – \| δ<sup>13</sup>C ↓ \| δ<sup>15</sup>N – \| C:N ↓ \| AEI ↓ |
|  | Other | 1690 | 1717 | 4 | 1703 | ASV Richness – \| δ<sup>13</sup>C ↓ \| δ<sup>15</sup>N ↓ \| C:N ↑ \| AEI ↑ |
|  | Relaxation | 1745 | 1753 | 4 | 1749 | ASV Richness ↑ \| δ<sup>13</sup>C – \| δ<sup>15</sup>N ↑ \| C:N ↓ \| AEI ↓ |
|  | Other | 1754 | 1871 | 4 | 1812 | ASV Richness – \| δ<sup>13</sup>C ↓ \| δ<sup>15</sup>N ↑ \| C:N ↑ \| AEI ↑ |
|  | Other | 1910 | 1981 | 5 | 1935 | ASV Richness ↓ \| δ<sup>13</sup>C ↑ \| δ<sup>15</sup>N ↑ \| C:N ↓ \| AEI ↓ |
| STO | Other | 1494 | 1513 | 4 | 1504 | ASV Richness – \| δ<sup>13</sup>C ↓ \| δ<sup>15</sup>N ↑ \| C:N ↓ \| AEI ↑ |
|  | Other | 1554 | 1573 | 4 | 1563 | ASV Richness – \| δ<sup>13</sup>C ↓ \| δ<sup>15</sup>N ↑ \| C:N ↓ \| AEI ↑ |
|  | Other | 1874 | 1893 | 4 | 1883 | ASV Richness – \| δ<sup>13</sup>C ↓ \| δ<sup>15</sup>N ↓ \| C:N ↑ \| AEI ↓ |
|  | Other | 1914 | 1933 | 3 | 1923 | ASV Richness – \| δ<sup>13</sup>C ↓ \| δ<sup>15</sup>N ↑ \| C:N ↑ \| AEI – |
|  | Other | 1954 | 1973 | 3 | 1963 | ASV Richness ↑ \| δ<sup>13</sup>C – \| δ<sup>15</sup>N ↑ \| C:N ↓ \| AEI – |
| CCO | Other | 1940 | 1946 | 5 | 1943 | ASV Richness ↓ \| δ<sup>13</sup>C ↑ \| δ<sup>15</sup>N ↑ \| C:N ↓ \| AEI ↓ |
|  | Intensification | 1952 | 1953 | 5 | 1952 | ASV Richness ↓ \| δ<sup>13</sup>C ↑ \| δ<sup>15</sup>N ↑ \| C:N ↑ \| AEI ↑ |
|  | Intensification | 1959 | 1963 | 5 | 1961 | ASV Richness ↓ \| δ<sup>13</sup>C ↓ \| δ<sup>15</sup>N ↑ \| C:N ↑ \| AEI ↑ |
|  | Other | 1964 | 1964 | 4 | 1964 | ASV Richness – \| δ<sup>13</sup>C ↓ \| δ<sup>15</sup>N ↑ \| C:N ↑ \| AEI ↑ |
|  | Other | 1967 | 1973 | 3 | 1970 | ASV Richness – \| δ<sup>13</sup>C ↓ \| δ<sup>15</sup>N – \| C:N ↑ \| AEI ↑ |
|  | Other | 1977 | 1994 | 5 | 1987 | ASV Richness ↓ \| δ<sup>13</sup>C ↓ \| δ<sup>15</sup>N ↑ \| C:N ↓ \| AEI ↓ |
|  | Intensification | 1995 | 1998 | 5 | 1996 | ASV Richness ↓ \| δ<sup>13</sup>C ↓ \| δ<sup>15</sup>N ↑ \| C:N ↓ \| AEI ↑ |
|  | Other | 1999 | 2003 | 4 | 2000 | ASV Richness ↓ \| δ<sup>13</sup>C ↓ \| δ<sup>15</sup>N ↑ \| C:N ↓ \| AEI – |
|  | Relaxation | 2004 | 2005 | 5 | 2004 | ASV Richness ↑ \| δ<sup>13</sup>C ↓ \| δ<sup>15</sup>N ↓ \| C:N ↑ \| AEI ↓ |
|  | Other | 2006 | 2011 | 5 | 2008 | ASV Richness ↓ \| δ<sup>13</sup>C ↓ \| δ<sup>15</sup>N ↓ \| C:N ↑ \| AEI ↓ |
|  | Other | 2014 | 2024 | 4 | 2019 | ASV Richness – \| δ<sup>13</sup>C ↓ \| δ<sup>15</sup>N ↓ \| C:N ↑ \| AEI ↑ |
| COL | Other | 1949 | 1957 | 5 | 1953 | ASV Richness ↓ \| δ<sup>13</sup>C ↑ \| δ<sup>15</sup>N ↑ \| C:N ↓ \| AEI ↓ |
|  | Other | 1959 | 1971 | 5 | 1964 | ASV Richness ↓ \| δ<sup>13</sup>C ↓ \| δ<sup>15</sup>N ↓ \| C:N ↓ \| AEI ↓ |
|  | Relaxation | 1972 | 1972 | 5 | 1972 | ASV Richness ↑ \| δ<sup>13</sup>C ↓ \| δ<sup>15</sup>N ↓ \| C:N ↓ \| AEI ↓ |
|  | Other | 1973 | 1979 | 4 | 1976 | ASV Richness ↑ \| δ<sup>13</sup>C ↓ \| δ<sup>15</sup>N ↑ \| C:N ↓ \| AEI – |
|  | Relaxation | 1980 | 1982 | 5 | 1981 | ASV Richness ↑ \| δ<sup>13</sup>C ↓ \| δ<sup>15</sup>N ↑ \| C:N ↓ \| AEI ↓ |
|  | Other | 1983 | 1984 | 4 | 1983 | ASV Richness ↑ \| δ<sup>13</sup>C ↓ \| δ<sup>15</sup>N ↑ \| C:N ↓ \| AEI – |
|  | Relaxation | 1985 | 1987 | 5 | 1986 | ASV Richness ↑ \| δ<sup>13</sup>C ↑ \| δ<sup>15</sup>N ↑ \| C:N ↓ \| AEI ↓ |
|  | Other | 1988 | 1993 | 4 | 1990 | ASV Richness – \| δ<sup>13</sup>C ↑ \| δ<sup>15</sup>N ↑ \| C:N ↓ \| AEI ↑ |
|  | Relaxation | 2006 | 2008 | 4 | 2007 | ASV Richness ↑ \| δ<sup>13</sup>C ↑ \| δ<sup>15</sup>N – \| C:N ↓ \| AEI ↓ |
|  | Other | 2010 | 2020 | 5 | 2015 | ASV Richness ↑ \| δ<sup>13</sup>C ↑ \| δ<sup>15</sup>N ↑ \| C:N ↓ \| AEI ↑ |
|  | Intensification | 2021 | 2024 | 4 | 2022 | ASV Richness ↓ \| δ<sup>13</sup>C – \| δ<sup>15</sup>N ↑ \| C:N ↓ \| AEI ↑ |
| CSD | Other | 1825 | 1844 | 4 | 1834 | ASV Richness – \| δ<sup>13</sup>C ↑ \| δ<sup>15</sup>N ↓ \| C:N ↓ \| AEI ↑ |
|  | Other | 1965 | 1984 | 3 | 1974 | ASV Richness – \| δ<sup>13</sup>C ↓ \| δ<sup>15</sup>N ↑ \| C:N – \| AEI ↑ |
| CMB | Intensification | 1665 | 1666 | 5 | 1665 | ASV Richness ↓ \| δ<sup>13</sup>C ↑ \| δ<sup>15</sup>N ↑ \| C:N ↓ \| AEI ↑ |
|  | Other | 1667 | 1698 | 4 | 1682 | ASV Richness – \| δ<sup>13</sup>C ↑ \| δ<sup>15</sup>N ↑ \| C:N ↓ \| AEI ↑ |
|  | Other | 1705 | 1764 | 4 | 1737 | ASV Richness – \| δ<sup>13</sup>C ↓ \| δ<sup>15</sup>N ↓ \| C:N ↓ \| AEI ↓ |
|  | Other | 1785 | 1804 | 4 | 1794 | ASV Richness – \| δ<sup>13</sup>C ↓ \| δ<sup>15</sup>N ↑ \| C:N ↓ \| AEI ↓ |
|  | Other | 1812 | 1818 | 3 | 1815 | ASV Richness – \| δ<sup>13</sup>C – \| δ<sup>15</sup>N ↓ \| C:N ↓ \| AEI ↑ |
|  | Other | 1823 | 1864 | 4 | 1843 | ASV Richness – \| δ<sup>13</sup>C ↑ \| δ<sup>15</sup>N ↓ \| C:N ↑ \| AEI ↑ |
|  | Other | 1885 | 1967 | 4 | 1926 | ASV Richness – \| δ<sup>13</sup>C ↓ \| δ<sup>15</sup>N ↑ \| C:N ↓ \| AEI ↓ |
|  | Relaxation | 1968 | 1979 | 5 | 1973 | ASV Richness ↑ \| δ<sup>13</sup>C ↓ \| δ<sup>15</sup>N ↑ \| C:N ↑ \| AEI ↓ |
|  | Other | 1980 | 2018 | 4 | 1999 | ASV Richness – \| δ<sup>13</sup>C ↓ \| δ<sup>15</sup>N ↑ \| C:N ↑ \| AEI ↓ |
| HIT | Other | 1905 | 1905 | 4 | 1905 | ASV Richness – \| δ<sup>13</sup>C ↓ \| δ<sup>15</sup>N ↓ \| C:N ↓ \| AEI ↓ |
|  | Other | 1920 | 1925 | 4 | 1922 | ASV Richness – \| δ<sup>13</sup>C ↑ \| δ<sup>15</sup>N ↑ \| C:N ↓ \| AEI ↓ |
|  | Other | 1958 | 1958 | 4 | 1958 | ASV Richness – \| δ<sup>13</sup>C ↑ \| δ<sup>15</sup>N ↑ \| C:N ↓ \| AEI ↓ |
|  | Other | 1967 | 1972 | 4 | 1969 | ASV Richness – \| δ<sup>13</sup>C ↓ \| δ<sup>15</sup>N ↓ \| C:N ↓ \| AEI ↑ |
|  | Other | 1977 | 1982 | 4 | 1979 | ASV Richness – \| δ<sup>13</sup>C ↑ \| δ<sup>15</sup>N ↓ \| C:N ↑ \| AEI ↑ |
|  | Other | 1991 | 1991 | 3 | 1991 | ASV Richness – \| δ<sup>13</sup>C ↓ \| δ<sup>15</sup>N – \| C:N ↑ \| AEI ↓ |
|  | Other | 2001 | 2001 | 4 | 2001 | ASV Richness – \| δ<sup>13</sup>C ↓ \| δ<sup>15</sup>N ↑ \| C:N ↓ \| AEI ↑ |
|  | Other | 2006 | 2023 | 4 | 2016 | ASV Richness – \| δ<sup>13</sup>C ↓ \| δ<sup>15</sup>N ↑ \| C:N ↑ \| AEI ↑ |
| HIO | Other | 1942 | 1956 | 4 | 1947 | ASV Richness – \| δ<sup>13</sup>C ↑ \| δ<sup>15</sup>N ↑ \| C:N ↓ \| AEI ↓ |
|  | Relaxation | 1957 | 1959 | 5 | 1958 | ASV Richness ↑ \| δ<sup>13</sup>C ↑ \| δ<sup>15</sup>N ↑ \| C:N ↑ \| AEI ↓ |
|  | Other | 1960 | 1964 | 5 | 1960 | ASV Richness ↑ \| δ<sup>13</sup>C ↑ \| δ<sup>15</sup>N ↑ \| C:N ↑ \| AEI ↑ |
|  | Relaxation | 1965 | 1966 | 5 | 1965 | ASV Richness ↑ \| δ<sup>13</sup>C ↓ \| δ<sup>15</sup>N ↓ \| C:N ↑ \| AEI ↓ |
|  | Other | 1970 | 1974 | 5 | 1970 | ASV Richness ↑ \| δ<sup>13</sup>C ↑ \| δ<sup>15</sup>N ↓ \| C:N ↓ \| AEI ↑ |
|  | Relaxation | 1975 | 1975 | 5 | 1975 | ASV Richness ↑ \| δ<sup>13</sup>C ↑ \| δ<sup>15</sup>N ↓ \| C:N ↓ \| AEI ↓ |
|  | Other | 1976 | 1998 | 5 | 1987 | ASV Richness ↓ \| δ<sup>13</sup>C ↓ \| δ<sup>15</sup>N ↓ \| C:N ↑ \| AEI ↓ |
|  | Intensification | 1999 | 1999 | 5 | 1999 | ASV Richness ↓ \| δ<sup>13</sup>C ↑ \| δ<sup>15</sup>N ↑ \| C:N ↑ \| AEI ↑ |
|  | Other | 2000 | 2024 | 5 | 2012 | ASV Richness ↑ \| δ<sup>13</sup>C ↑ \| δ<sup>15</sup>N ↓ \| C:N ↑ \| AEI ↑ |

**Fig. S1:**
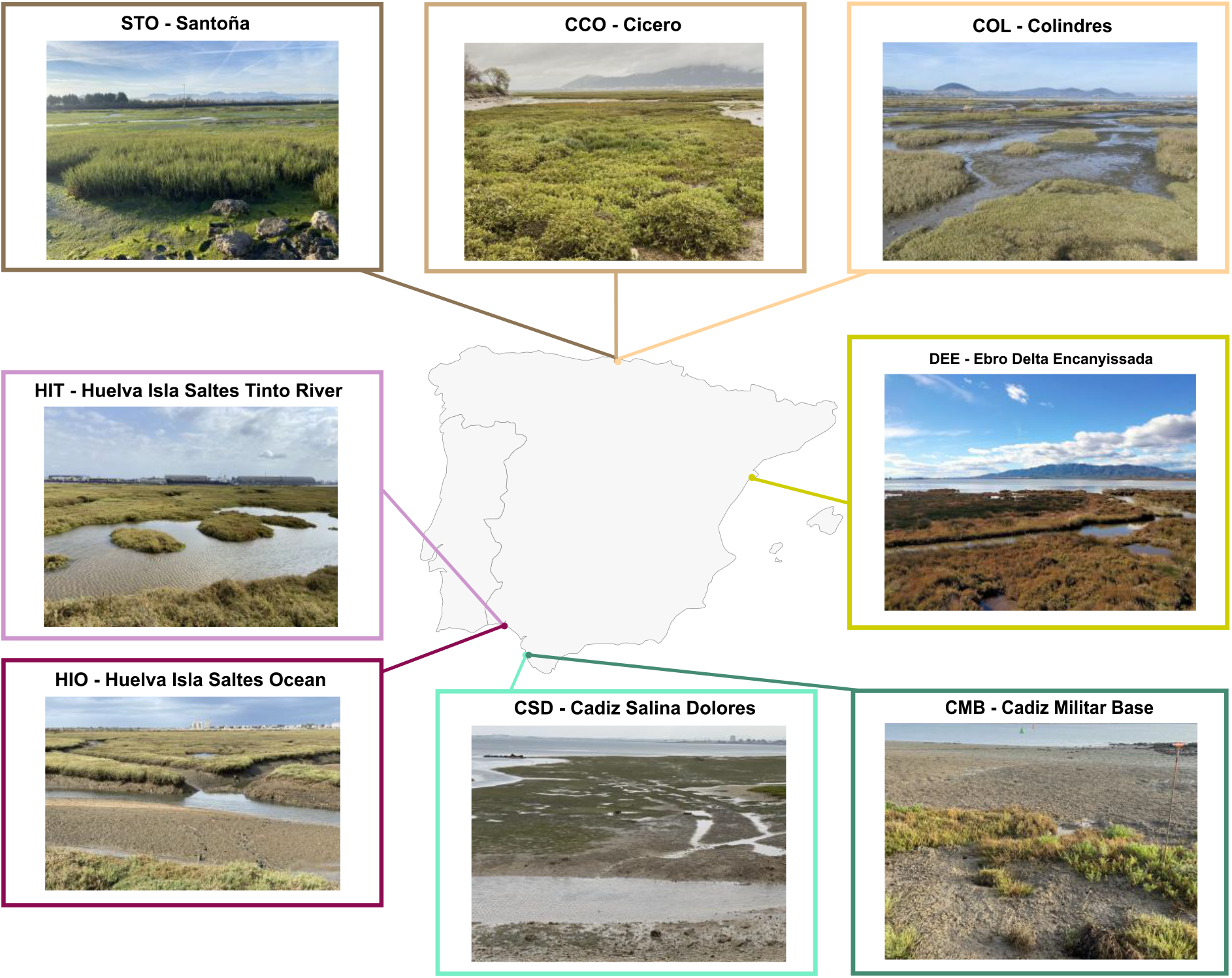
Location and overview of the eight coastal wetland study sites across the Iberian Peninsula. The map shows the geographical location of each sampling site together with photographs of the sampling sites: Santoña (STO), Cicero (CCO), Colindres (COL), Cádiz Salina Dolores (CSD), Cádiz Militar Base (CMB), Huelva Isla Saltés–Tinto River (HIT), Huelva Isla Saltés–Ocean (HIO), and Ebro Delta Encanyissada (DEE).

**Fig. S2:**
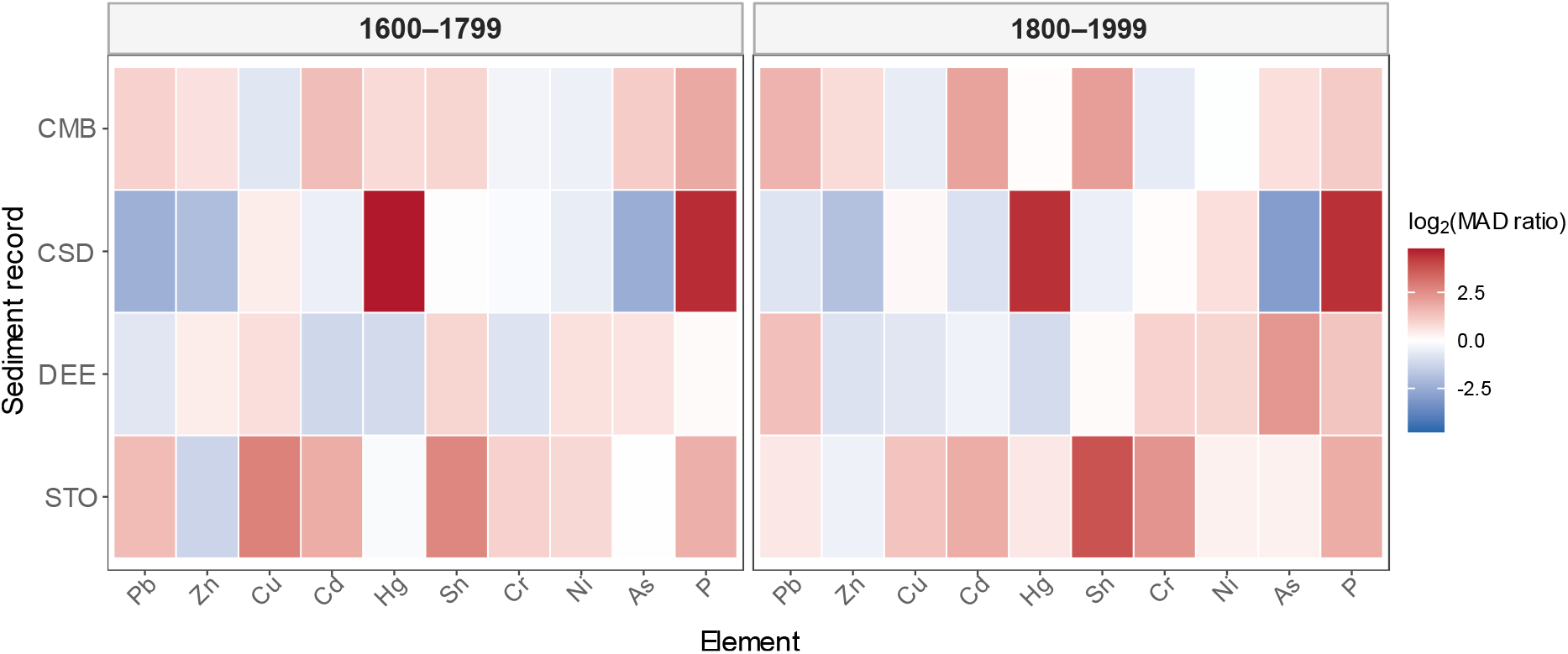
Elemental variability across consecutive 200-year windows. Tiles show the log₂ ratio between the scaled median absolute deviation (MAD) of CLR-transformed elemental values in each post-1600 window and the corresponding variability during 1400–1599. Positive values indicate greater variability than during 1400–1599, whereas negative values indicate lower variability. Only sediment records extending across the three temporal windows are shown, and windows containing fewer than five observations were excluded. The comparison was used as a descriptive diagnostic of variability around the operational 1600 CE cutoff and was not interpreted as evidence of a common regional geochemical breakpoint.

**Fig. S3:**
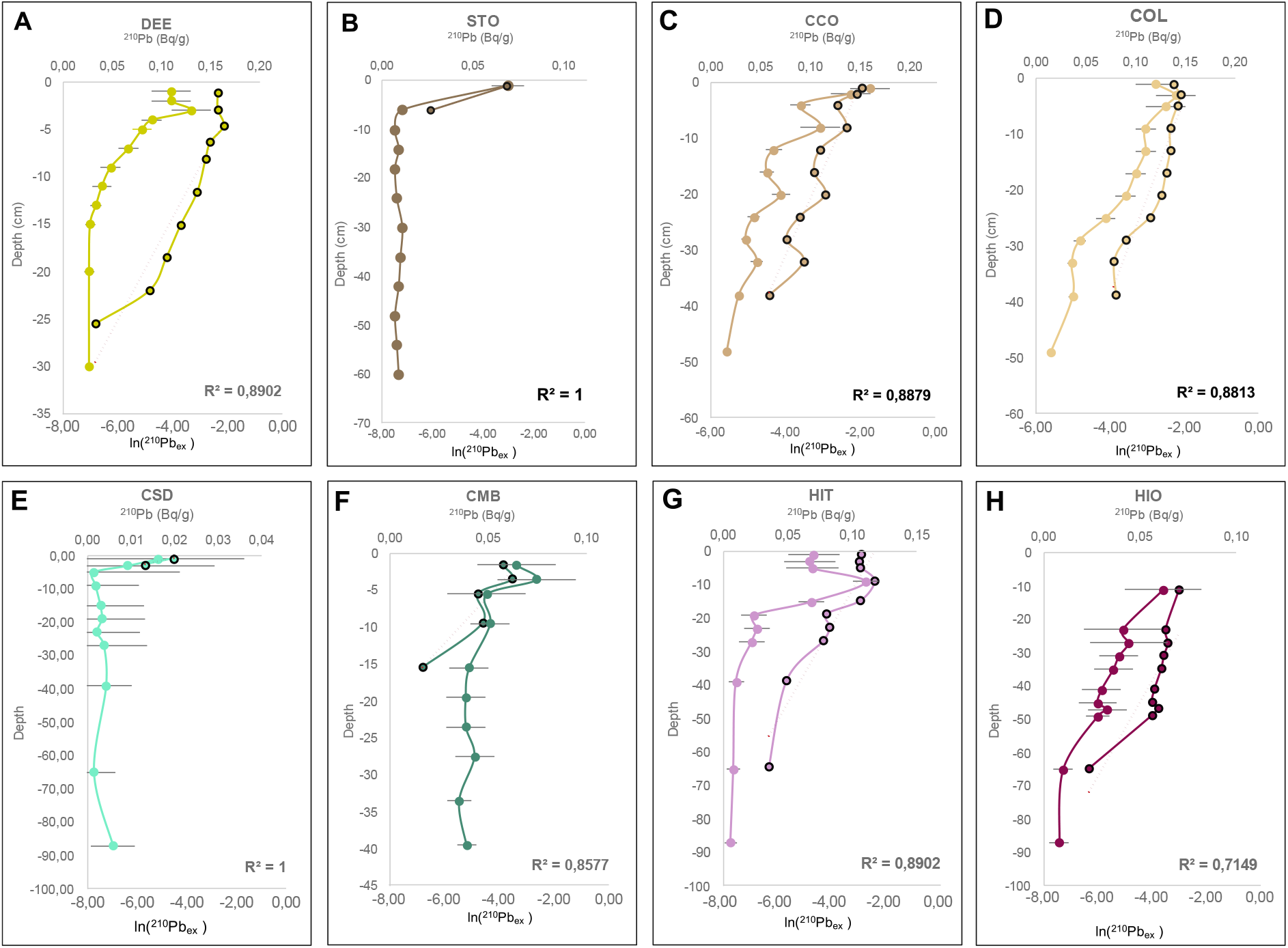
Excess ^210^Pb dating models for Iberian coastal sediment cores based on the Constant Flux–Constant Sedimentation (CF:CS) model. Panels show the relationship between the natural logarithm of unsupported ^210^Pb activity and sediment depth for each dated core. Linear regressions were used to estimate recent sediment accumulation rates (SARs) under the assumptions of constant sediment accumulation and ^210^Pb flux through time. Dashed lines represent the fitted regression models, and shaded areas show the corresponding regression uncertainty intervals, where estimable. The unsupported-^210^Pb dating horizon was defined as the deepest interval at which excess ^210^Pb remained distinguishable from supported background activity. Estimated SARs, uncertainty ranges and basal ages extrapolated under the assumption of constant recent accumulation are summarised in Table S4.

**Fig. S4:**
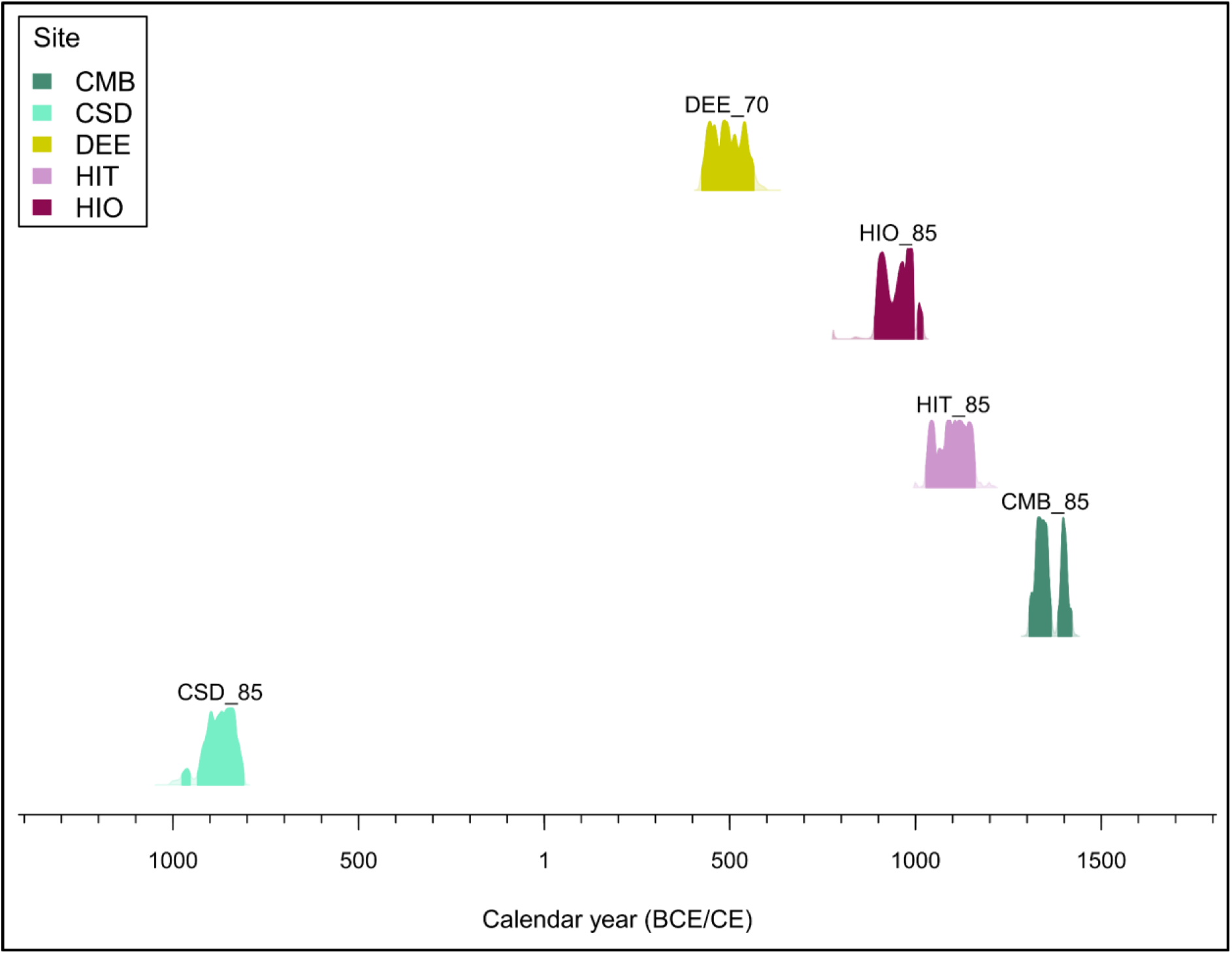
Calibrated AMS radiocarbon ages for intertidal sediment samples from five sediment cores. Conventional ^14^C ages were calibrated using the IntCal20 Northern Hemisphere atmospheric calibration curve. Coloured distributions represent the calibrated probability density for each sample, and horizontal bars indicate the intervals encompassing 95.4% of the calibrated probability distribution. Colours identify the corresponding sediment cores. No marine-reservoir or old-carbon correction was applied. The resulting age distributions represent the estimated ages of the dated carbon fractions and were not assumed a priori to correspond exactly to the depositional ages of the surrounding sediments. Conventional radiocarbon determinations and calibrated age ranges are summarised in Table S5.

**Fig. S5:**
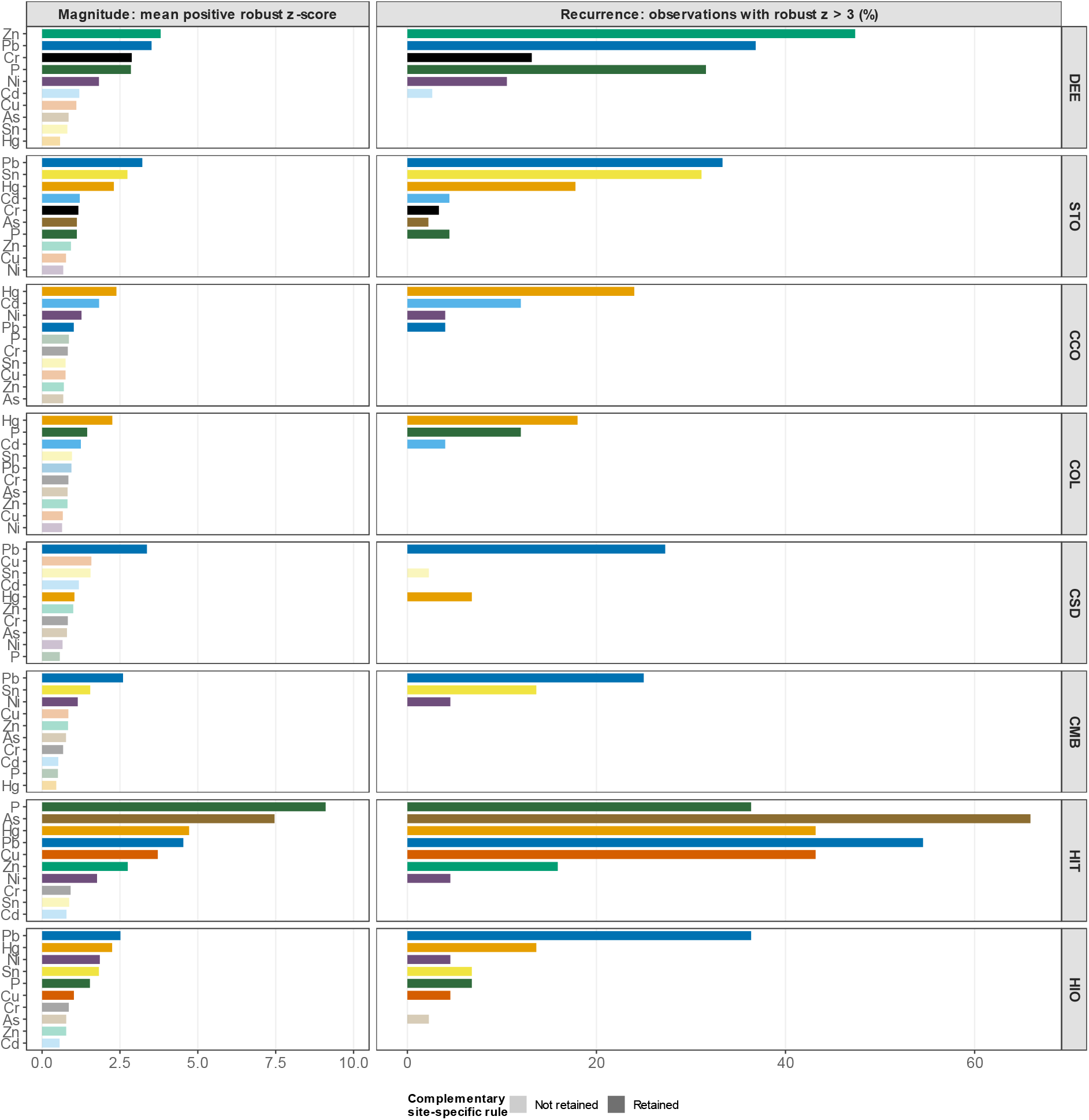
Magnitude and recurrence of enrichment among candidate anthropogenic elements at each study site. The left panels show enrichment magnitude, calculated as the mean robust z-score among observations with positive enrichment (z^robust^ > 0). The right panels show enrichment recurrence, calculated as the percentage of valid observations exhibiting strong enrichment (z^robust^ > 3). Elements are ordered within each site according to decreasing enrichment magnitude, and the same order is retained in both panels. Fully coloured bars identify elements retained for the complementary site-specific analysis, defined as elements with z^robust^ > 3 in at least two samples; faded bars indicate elements that did not meet this operational criterion. Colours identify individual elements and do not represent enrichment intensity. All enrichment scores were calculated relative to the corresponding site- and element-specific baseline.

**Fig. S6:**
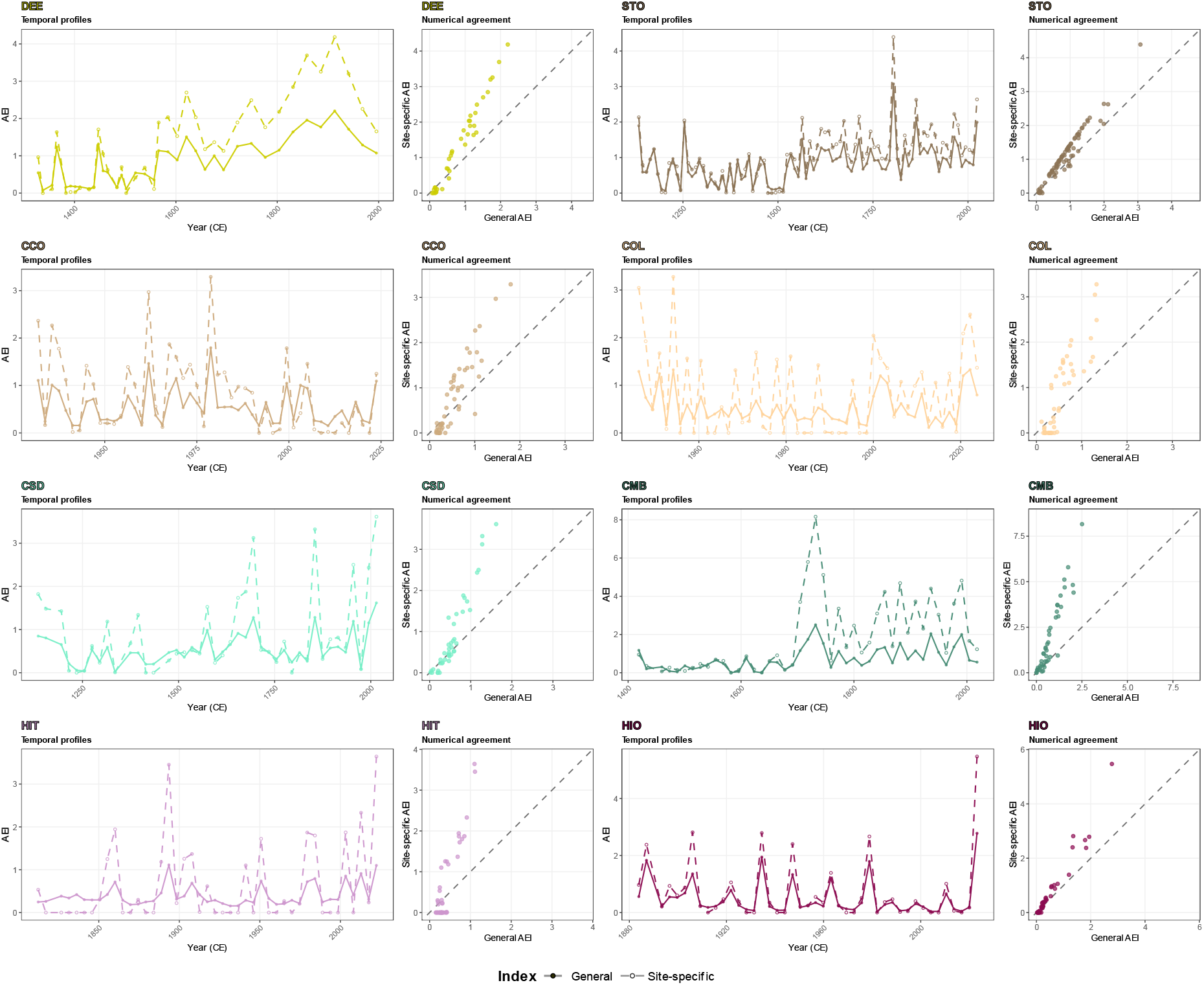
Temporal correspondence and numerical agreement between the general and site-specific Anthropogenic Enrichment Indices (AEIs). For each site, the left panel shows the temporal trajectories of the general AEI, calculated from all ten candidate elements with available XRF measurements, and the site-specific AEI, calculated from elements showing strong enrichment (z^robus^t > 3) in at least two samples. The adjacent right panel compares paired general and site-specific AEI values; the dashed diagonal denotes numerical identity. Points above this line indicate samples for which the site-specific AEI exceeded the general AEI, whereas points below it indicate higher general AEI values. Colours identify study sites. Note that axis ranges differ among sites to display within-record variation.

**Fig. S7:**
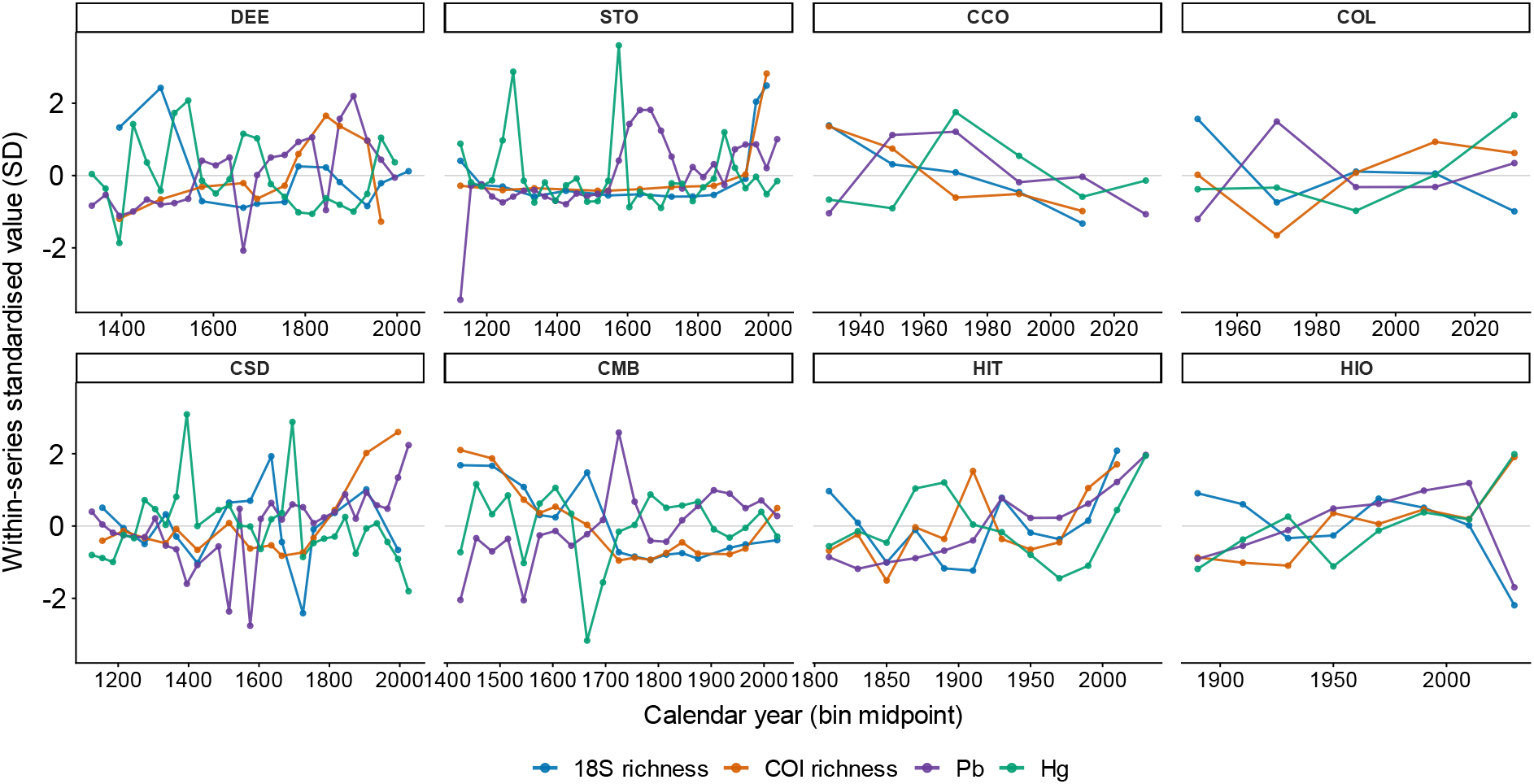
Temporal variation in eukaryotic ASV richness and element-specific Pb and Hg enrichment across the eight sediment records. Values represent medians calculated within 20-year intervals for CCO, COL, HIT and HIO and 30-year intervals for DEE, STO, CSD and CMB. Pb and Hg enrichment was expressed using element-specific robust z-scores. For visual comparison of temporal trajectories, 18S richness, COI richness, Pb and Hg were standardised separately within each sediment record and series to a mean of zero and standard deviation of one. Points represent retained temporal bins, and solid lines connect consecutive available bins. Standardised values therefore show relative temporal variation rather than differences in absolute richness or elemental concentration. Pb in COL and Hg in DEE and CMB did not meet the criterion for inclusion in the corresponding site-specific AEI but are shown for completeness.

**Fig. S8:**
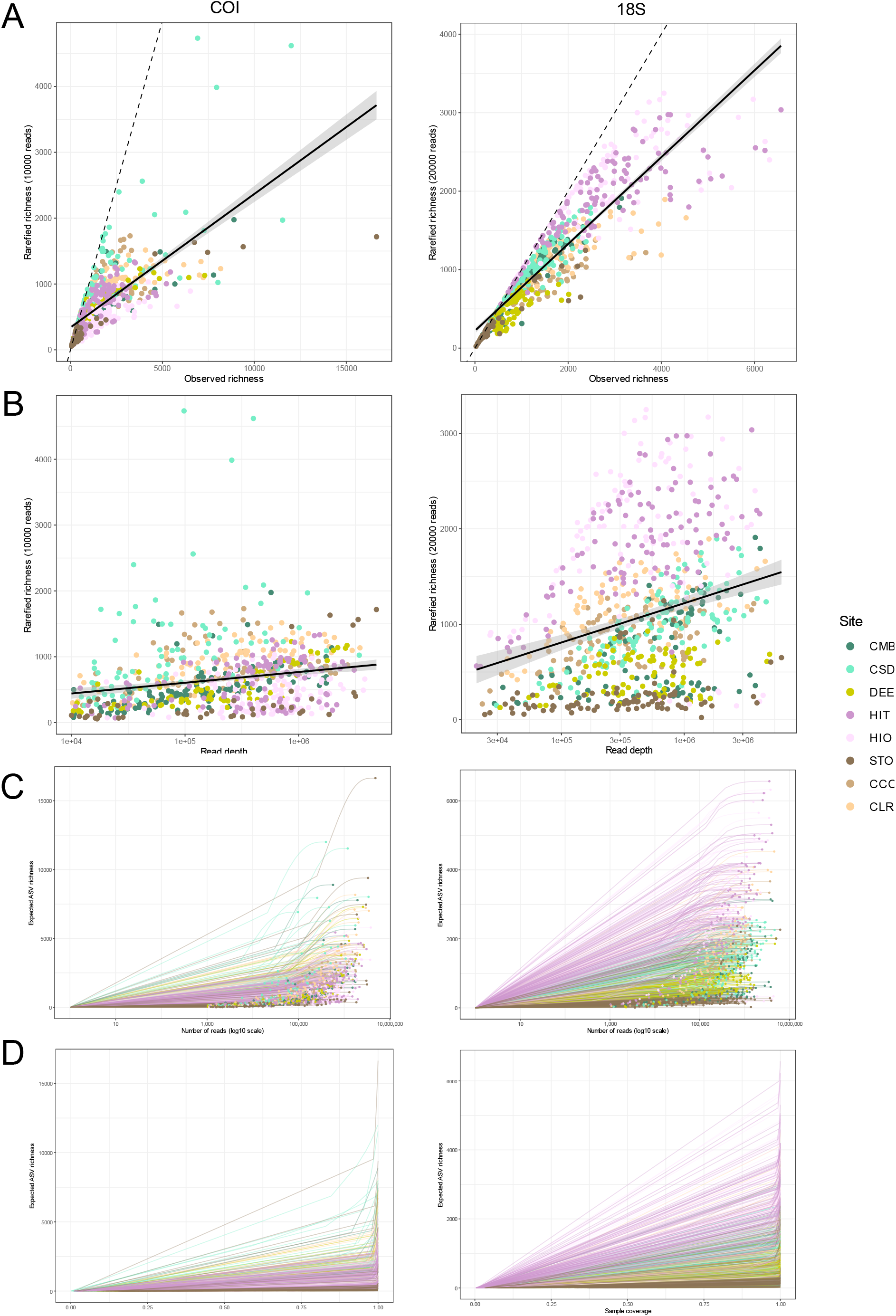
Sensitivity analysis of sequencing-depth standardisation and rarefaction/extrapolation analyses for sedimentary DNA metabarcoding datasets. (A) Relationship between observed and sample-size-rarefied ASV richness across PCR replicates for the COI and 18S markers after rarefaction to 10,000 reads (COI) and 20,000 reads (18S). (B) Relationship between rarefied ASV richness and sequencing depth, assessing the influence of read depth on richness estimates after standardisation. (C) Sample-size-based rarefaction and extrapolation curves generated with iNEXT for the COI and 18S datasets. Solid lines indicate interpolation, dashed lines indicate extrapolation, and shaded areas represent 95% confidence intervals. (D) Coverage-based rarefaction and extrapolation curves generated with iNEXT for the COI and 18S datasets, illustrating sample completeness and allowing diversity comparisons at equivalent levels of sampling coverage.

**Fig. S9:**
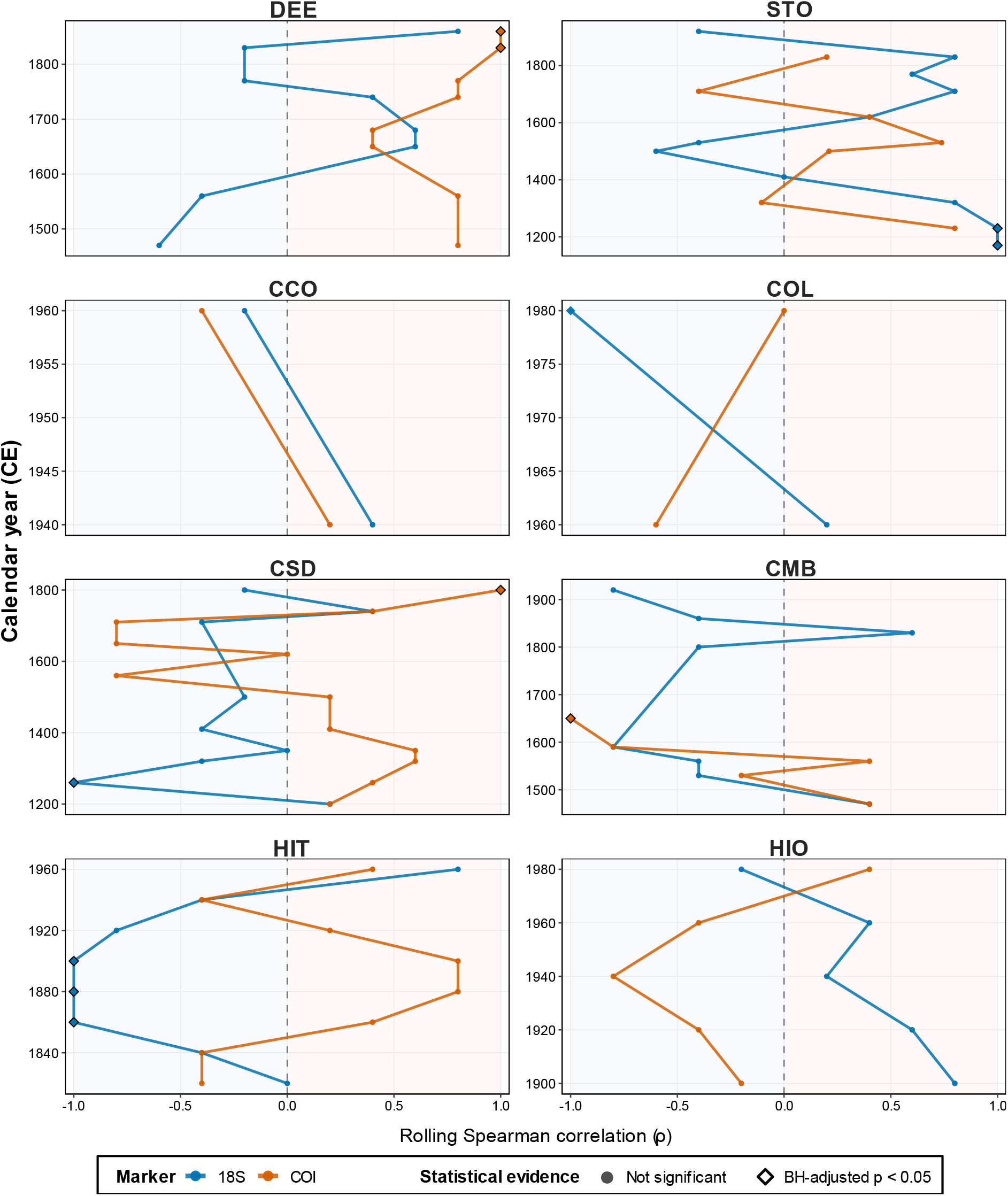
Temporal variation in the local association between eukaryotic ASV richness and the general Anthropogenic Enrichment Index (AEI). Lines show four-bin rolling Spearman correlations calculated separately for 18S and COI within each sediment record. Median richness and AEI values were calculated using 20-year bins for CCO, COL, HIT and HIO and 30-year bins for DEE, STO, CSD and CMB; rolling windows therefore span approximately 80 and 120 years, respectively. Positive values indicate that richness and AEI covaried in the same direction within a window, whereas negative values indicate opposing variation. Consecutive estimates are not independent because windows overlap. The rolling correlations were used as an exploratory description of temporal non-stationarity and were not interpreted as independent significance tests.

**Fig. S10:**
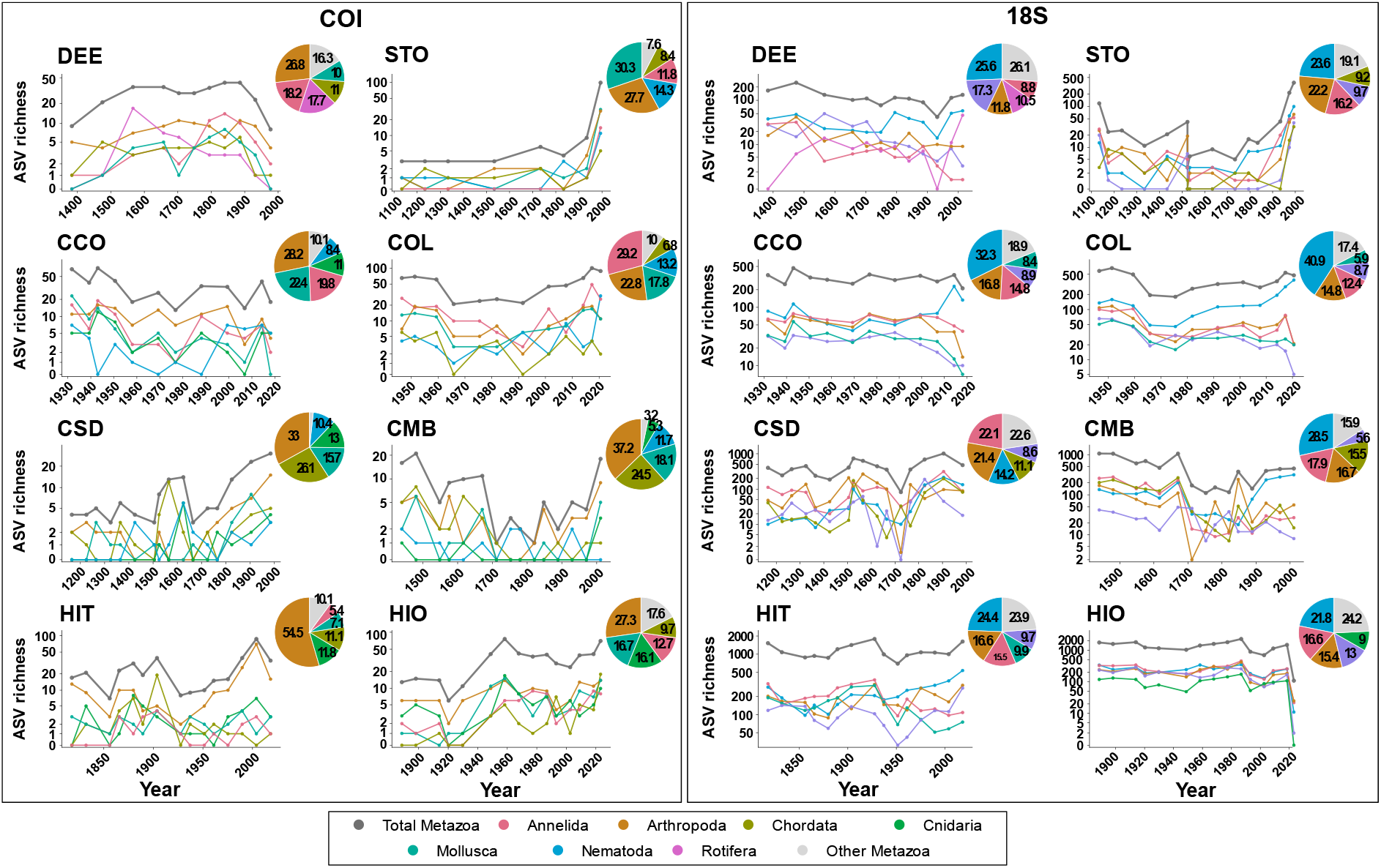
Phylum-level distribution and temporal patterns of metazoan ASV richness reconstructed from sedimentary DNA across Iberian coastal wetlands. Panels show total metazoan ASV richness and richness attributed to the five most ASV-rich metazoan phyla within each site and marker. Phyla not included among the corresponding five most ASV-rich groups were combined as “Other Metazoa”. Richness is displayed on a pseudo-logarithmic scale, and all records are plotted against calendar year. Pie charts show the percentage contribution of the five selected phyla and the pooled “Other Metazoa” category to the total number of unique metazoan ASVs detected across the corresponding site and marker. Pie-chart percentages represent pooled ASV richness rather than relative read abundance. Colours identify phyla and are maintained across panels whenever the same phylum occurs.

**Fig. S11:**
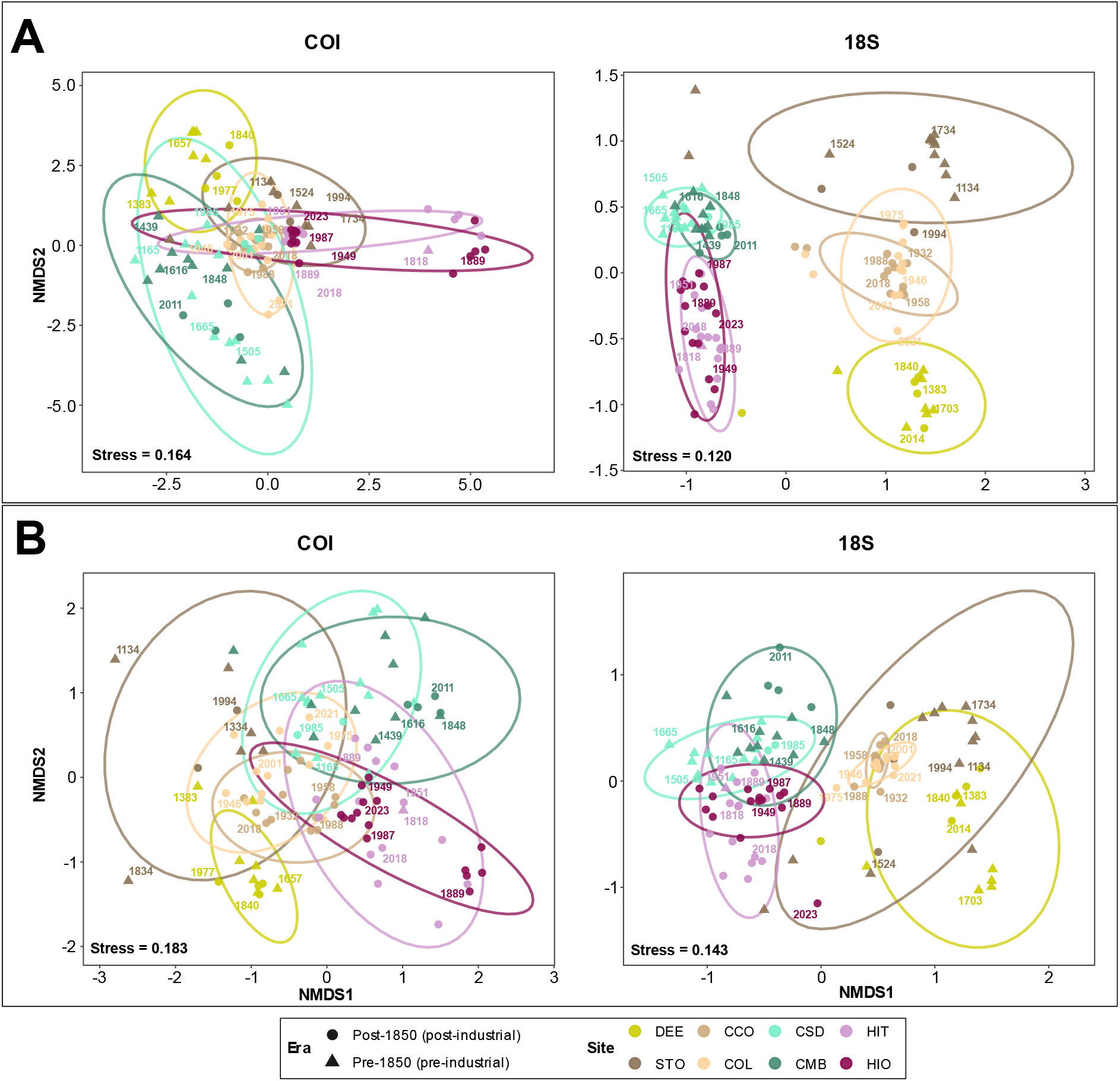
Community composition of eukaryotic and metazoan assemblages across Iberian coastal sediment records. Non-metric multidimensional scaling (nMDS) ordinations were calculated from Bray–Curtis dissimilarities of sample-wise relative ASV abundances after restricting the datasets to ASVs assigned to Eukaryota (A) or Metazoa (B). Ordinations were fitted separately for COI (left) and 18S (right) in three dimensions and are shown as projections onto NMDS1 and NMDS2. Points represent individual sediment samples, colours identify sediment records, and point shapes distinguish pre-industrial (pre-1850) and post-industrial (post-1850) samples. Labels show four representative calendar years distributed across each record. Coloured lines are 95% data ellipses calculated under a multivariate t-distribution and summarise within-record dispersion. Reported stress values correspond to the complete three-dimensional configurations.

## Supplementary methods

**Method S1.** Radionuclide measurements and radiocarbon calibration

Unsupported ^210^Pb activity was determined indirectly through measurement of its granddaughter radionuclide ^210^Po by alpha spectrometry following Flynn (1968). Sediment aliquots were spiked with ^209^Po as a yield element and digested in aqua regia. Polonium isotopes were subsequently auto-deposited onto silver discs and measured using ORTEC Alpha Octete spectrometers equipped with passivated implanted planar silicon detectors. The analytical detection limit was 0.1 Bq kg^−1^. Analyses were conducted at the GAU-Radioanalytical Laboratories, University of Southampton, UK.

Supported ^210^Pb activity was estimated from background activities measured in the deepest analysed sections of each core (Chepstow-Lusty et al., 2007). Unsupported, or excess, ^210^Pb activity was calculated by subtracting this supported component from total ^210^Pb activity. Sediment accumulation rates were estimated using the Constant Flux–Constant Sedimentation model (Krishnaswamy et al., 1971), implemented in Microsoft Excel by linear regression of the natural logarithm of unsupported ^210^Pb activity against sediment depth. The depth at which unsupported ^210^Pb declined to background activity was taken as the lower limit of the ^210^Pb dating horizon. Calendar ages were calculated from the core-specific sediment accumulation rates relative to the corresponding core-collection date. Below the unsupported-^210^Pb dating horizon, the same rates were extrapolated linearly to the core base.

Five sediment samples from DEE, CSD, CMB, HIT and HIO were analysed by accelerator mass spectrometry at the Poznań Radiocarbon Laboratory, Poland. Conventional ^14^C determinations were calibrated in R using the rcarbon package (v1.5.2; Crema & Bevan, 2021) and the IntCal20 Northern Hemisphere atmospheric calibration curve (Reimer et al., 2020). Calibrated ages were summarised using 95.4% probability ranges. No marine-reservoir correction or empirical correction for old or reworked carbon was applied. IntCal20 was selected under the operational assumption that the dated carbon was predominantly atmospheric or terrestrial in origin. Because the dated carbon fraction was not independently characterised, these calibrated ages were not incorporated into the CF age–depth models.

**Method S2.** XRF quality control and preprocessing

Raw X-ray fluorescence measurements were screened for isolated positive spikes using a moving-window Hampel-type filter based on the local median and median absolute deviation (MAD) (Pearson et al., 2016). For each elemental profile, a symmetric window containing the focal observation and up to two neighbouring observations on either side was used (k = 2). An observation was flagged as a potential positive spike when it exceeded the local median by more than 2.5 MAD-based scale units (t₀ = 2.5).

Because missing or non-detect values prevented consistent application of the Hampel criterion in some sections of the profiles, a supplementary nearest-neighbour rule was applied. An observation was flagged when it exceeded five times the median of the nearest valid neighbouring observations. Values identified by either criterion were treated as potential analytical artefacts and replaced by the median of the adjacent valid observations. Observations that did not meet either criterion were not modified.

Measurements identified as potential analytical artefacts were treated as missing rather than replaced by a local estimate. The window width and detection thresholds were defined as operational quality-control parameters and applied consistently across elements and sediment cores; they were not interpreted as universal geochemical outlier criteria. Before centred log-ratio transformation, missing and non-positive measurements were replaced separately for each element by half the minimum positive value observed for that element across the dataset, as described in Method S3.

**Method S3.** Construction and sensitivity analysis of the Anthropogenic Enrichment Index

### Selection and log-ratio transformation of candidate elements

Within each sediment record, potential isolated analytical artefacts were identified separately for each element using a moving-window Hampel-type filter. The local median and median absolute deviation (MAD) were calculated within a symmetric window containing the focal observation and up to two neighbouring observations on either side. Measurements differing from the local median by more than 2.5 MAD-based scale units were flagged. Because missing values prevented this criterion from being applied consistently in some parts of the profiles, an additional nearest-neighbour criterion was used: a measurement was flagged when it exceeded five times the median of the nearest valid observations preceding and following it. Measurements identified by either criterion were treated as missing. These thresholds were applied consistently across sediment records as operational quality-control criteria and were not interpreted as general geochemical thresholds for identifying anomalous enrichment.

Eleven candidate elements were defined a priori as potential indicators of metal contamination or nutrient enrichment: lead (Pb), zinc (Zn), copper (Cu), antimony (Sb), cadmium (Cd), mercury (Hg), tin (Sn), chromium (Cr), nickel (Ni), arsenic (As) and phosphorus (P). Their documented associations with industrial, urban, mining, maritime, agricultural and wastewater-related inputs, together with the rationale and supporting references for their inclusion, are summarised in Table S2a. These elements were considered collectively because anthropogenic modification of coastal wetlands may involve multiple and overlapping sources of elemental inputs rather than a single contaminant or enrichment pathway. However, their ecological effects are not equivalent: some elements may be associated with potentially harmful contamination, whereas others, particularly P, may reflect nutrient enrichment with context-dependent biological consequences. The index was therefore designed to summarise relative multielement enrichment rather than to represent a single ecological stressor or predict a uniform direction of biodiversity response. Sb was excluded because measurements were unavailable for some sediment cores, preventing its consistent inclusion in cross-record comparisons. The remaining ten elements were used to calculate the enrichment indices, and site-specific means and standard deviations of their cleaned XRF measurements are reported in Table S2b.

The retained elemental measurements were treated as compositional information and transformed using the centred log-ratio transformation (Aitchison, 1982; Weltje & Tjallingii, 2008):

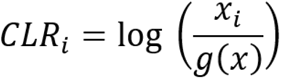

where x_ᵢ_ is the value of element i and g(x) is the geometric mean of the candidate elements within the same sample. Because the CLR transformation requires complete and strictly positive compositions, missing and non-positive measurements were replaced separately for each element by half the minimum positive value observed for that element across the complete analysed dataset. This procedure was applied consistently to all samples and sediment records as an operational preprocessing step and should not be interpreted as estimating the true elemental concentrations of missing or non-detect measurements (Martín-Fernández et al., 2003).

### Diagnostic assessment of elemental variability around the historical cutoff

To examine whether elemental variability changed around the operational 1600 CE cutoff, the robust variability of CLR-transformed values was calculated separately for each element and sediment record within fixed 200-year windows. Comparisons focused on 1400–1599, 1600– 1799 and 1800–1999, thereby maintaining an equivalent nominal duration before and after the cutoff. Robust variability was quantified as 1.4826 times the median absolute deviation (MAD), consistent with the scale estimator used to calculate element-specific enrichment scores. Windows containing fewer than five observations were excluded. This analysis was treated as a descriptive diagnostic and not as a test of whether 1600 CE represented a common geochemical breakpoint.

The two reference strategies therefore differed in their historical interpretation. For records with pre-1600 observations, enrichment was evaluated relative to an early historical reference, whereas for shorter records it represented deviations from the corresponding within-record elemental distribution. Although both approaches allowed changes in relative multielement enrichment to be characterised within individual records, AEI values were not interpreted as directly comparable measures of anthropogenic pressure intensity among wetlands.

### Magnitude, recurrence and site-specific element selection

The magnitude and recurrence of enrichment were evaluated separately for each candidate element and sediment record. Enrichment magnitude was calculated as the mean robust z-score among observations showing positive enrichment:

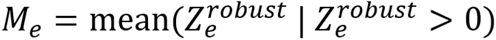

where Me is the magnitude of positive enrichment for element e.

Enrichment recurrence was calculated as the percentage of all valid observations showing strong enrichment:

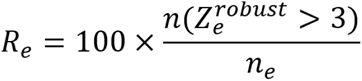

where Re is the recurrence of strong enrichment and ne is the total number of valid observations for element (e) in the corresponding sediment record.

Magnitude and recurrence were not combined into a composite importance score. This allowed elements showing large enrichment in a small number of observations to be distinguished from those showing repeated strong enrichment. An element was retained in the site-specific set when (z^robust^ > 3) occurred in at least two observations from the corresponding sediment record. This count-based criterion was operational and was not interpreted as a universal boundary for sediment contamination.

### Sensitivity of the AEI to site-specific element selection

The general and site-specific AEIs were compared to evaluate whether restricting the index to locally recurrent elements materially altered the reconstructed enrichment histories. Temporal profiles of both formulations were first compared graphically within each site.

Spearman’s rank correlation coefficient (Spearman, 1961) was used to quantify whether the two indices produced similar temporal rankings of samples without assuming numerical equivalence. Absolute agreement was evaluated using Lin’s concordance correlation coefficient (CCC), which incorporates both precision and deviation from the line of identity (Lin, 1989). Systematic differences in magnitude were additionally described using the paired difference:

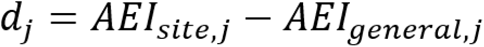

Because differences in scale can reduce raw concordance even when temporal trajectories are similar, both AEIs were additionally converted to within-site standardised scores, and their concordance was recalculated as a sensitivity analysis. This standardised comparison evaluated similarity in temporal trajectories after removing site-specific differences in location and scale; it was not interpreted as evidence of agreement in the original AEI units.

The general AEI was defined a priori as the primary formulation for subsequent cross-record analyses because it maintained a common element set, weighting scheme and denominator. The site-specific formulation was used only as a sensitivity analysis.

### Sensitivity analysis using Pb and Hg

To determine whether the richness–AEI relationships were reproduced using individual well-established pollutants, the analyses were repeated separately for Pb and Hg (Fig. S7; Table S9). Element-specific robust z-scores were aggregated using the same site-specific temporal intervals and median summaries applied in the richness–AEI analyses. Spearman correlations and standardised linear models were calculated for each sediment record and marker. Additional models included standardised calendar year as a predictor to separate the element–richness association from their potentially shared linear temporal trends, and models with a continuous first-order autoregressive residual structure were fitted when sufficient temporal bins were available. Results were interpreted as a complementary sensitivity analysis rather than as pollutant-specific dose–response relationships.

